# A Neural Arming Niche in Tumor-Draining Lymph Nodes Programs CD8⁺T Cell Cytotoxicity via GZMB Norepinephrinylation

**DOI:** 10.64898/2026.02.25.707881

**Authors:** Yan Yang, Xue-Li Zhang, Aziguli Tulamaiti, Shu-Yu Xiao, Yun-Zhen Qian, Jia-Mei Luo, Guang-hong Su, Rui Lu, Jun-Jie Wang, Hong-Tai Ma, Xia-Qing Li, Wen-Tao Shi, Yuan-Xin Hong, Jing-Li Hou, Li-Peng Hu, Xin Xing, Qing Li, Dong-Xue Li, Zhi-Gang Zhang

## Abstract

Lymph nodes are densely innervated neuro-immune hubs where naive T cells are primed and “armed” with cytotoxic machinery, yet how local neural cues in tumor-draining lymph nodes (tdLNs) set CD8⁺ T cell effector reserves remains unclear. Here we show that tdLN-local sympathetic norepinephrine (NE) instructs CD8⁺ T cell cytotoxic capacity by stabilizing the granzyme B (GZMB) pool through covalent norepinephrinylation. Exercise, used as a physiological perturbation, increases GZMB selectively in tdLN and tumor CD8⁺ T cells without altering T cell abundance or cytokine output; tdLN sympathectomy abolishes these effects and reduces tdLN NE enrichment. Mechanistically, NE enters CD8⁺ T cells and norepinephrinylates GZMB at Gln43, limiting UHRF1-mediated ubiquitination and proteasomal degradation. Genetic attenuation of norepinephrinylation destabilizes GZMB, impairs cytotoxicity, and accelerates tumor growth. Brief *ex vivo* NE conditioning increases GZMB reserves and improves adoptive tumor control.

## Introduction

CD8⁺ cytotoxic T lymphocytes (CTLs) are principal effectors of antiviral and antitumor immunity, and their capacity to execute target-cell killing is a major determinant of protective immune control^1,2^. Yet cytotoxic competence is not hardwired at the moment of antigen encounter. Even under comparable antigenic stimulation, effector CD8⁺ T cells can differ substantially in the magnitude of their cytotoxic effector reserves, implying that physiological inputs beyond antigen and co-stimulation shape the effector program^3–7^. Identifying such inputs, particularly during early priming, is therefore essential for understanding how cytotoxic potential is set and how it might be therapeutically enhanced.

The draining lymph node (dLN) is a key anatomical site where naïve CD8⁺ T cells are primed, clonally expanded, and instructed to acquire effector functions^8–10^. Upon productive interactions with antigen-presenting cells, newly activated CD8⁺ T cells rapidly initiate transcriptional and translational programs that build the cytotoxic machinery in dLNs^9,11,12^. Among these components, granzyme B (GZMB) is a core effector molecule required for CTL-mediated killing^13,14^. Notably, GZMB is subject to stringent post-translational control: after synthesis, it must be efficiently routed into lysosome-like secretory granules, whereas unincorporated cytosolic GZMB is potentially harmful and is therefore prone to rapid ubiquitination and proteasomal degradation^15^. Consequently, the ability of newly primed CD8⁺ T cells to stabilize and accumulate GZMB during the priming window in the dLNs is expected to set the “ceiling” of their cytotoxic effector reserves, thereby constraining subsequent cytotoxic potency. However, the local physiological cues and molecular mechanisms within the dLNs that govern GZMB protein stability during early effector programming remain poorly defined.

Lymph nodes are richly innervated by sympathetic fibers, and norepinephrine (NE) levels within lymphoid tissues fluctuate with physiological states such as exercise and stress^16–18^. These observations raise the possibility that sympathetic neurotransmission provides a dynamic, compartment-restricted signal that can tune effector programming in the dLNs, potentially by regulating the balance between synthesis, storage, and degradation of cytotoxic molecules in CD8⁺ T cells. NE is classically viewed as acting through adrenergic receptors to engage G protein-coupled signaling pathways^19,20^. In parallel, accumulating evidence indicates that NE can also exert receptor-independent effects through covalent post-translational modification of proteins, a process termed “norepinephrinylation”^21–24^, yet whether such “norepinephrinylation” occurs in CD8⁺ T cells and contributes to cytotoxic programming has remained unclear.

Here we identify a sympathetic-NE axis in the dLNs that directly controls CD8⁺ T cell cytotoxic effector reserves by stabilizing GZMB protein. We show that NE can enter CD8⁺ T cells and promote a covalent modification of GZMB at Gln43 mediated by transglutaminase 2 (TGM2). This modification limits UHRF1-dependent ubiquitination and proteasomal degradation of GZMB, thereby increasing the available pool of cytotoxic effector molecules without broadly amplifying inflammatory cytokine programs. By linking a tdLNs-local neural cue to a defined post-translational mechanism that sets cytotoxic capacity, our study reveals a neuro-immune checkpoint at the level of effector molecule homeostasis and suggests a minimal, controllable strategy to potentiate CD8⁺ T cell function through *ex vivo* NE conditioning for adoptive cell therapy.

## Results

### tdLNs-local sympathetic norepinephrine signaling as an instructive cue for CD8⁺ T cell cytotoxic effector reserves

Tumor-draining lymph nodes (tdLNs) are a critical site where tumor-specific CD8⁺ T cells are primed and where cytotoxic effector molecule reserves are established^11^. We reasoned that a tdLNs-local cue might control the magnitude of this reserve and thereby set cytotoxic potency. To expose such a cue *in vivo*, we used exercise as a tractable physiological perturbation that elevates sympathetic activity^25^. In a mouse subcutaneous tumor model, two exercise regimens were applied: a low-intensity^26,27^ treadmill-running protocol (5 sessions/week, 30 min/session, 15 cm/sec) and voluntary wheel running. In a subcutaneous tumor implantation model, Evans blue^28^ mapping identified the inguinal lymph node as the dominant tdLNs (Extended Data Fig. 1a). Exercise robustly suppressed tumor growth (Fig. 1a-d, and Extended Data Fig.1 b and c), providing a physiological entry point to interrogate upstream regulatory signals.

**Fig. 1.**
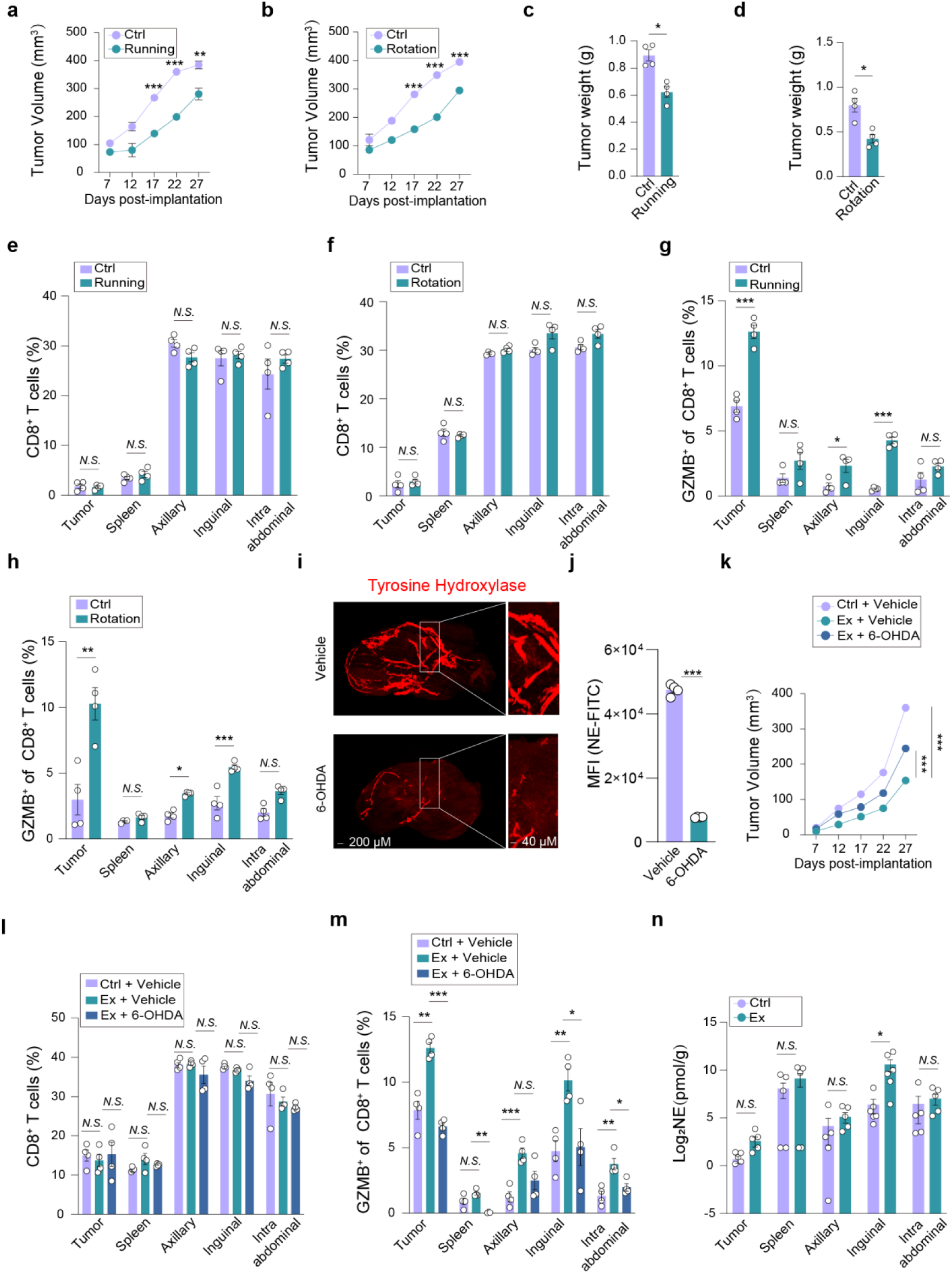
Tumor suppression requires norepinephrine release from sympathetic nerves mainly in draining lymph node. **a-d,** KPC1199 tumor cells were subcutaneously implanted into mice. Tumor volume (**a, b**) was monitored every 5 days, and tumor weight (**c, d**) was measured at 27days after implantation (n = 4 per group, 3 independent experiments). **e-f,** Percentage of CD8⁺ T cells were detected at 27 days after KPC1199 implantation. Runnig model (**e**), and voluntary wheel running model (**f**). Intra-abdominal lymph nodes include mesenteric and renal hilar/para-aortic lymph nodes (n = 4 per group, 3 independent experiments). **g-h,** Intracellular GZMB expression of CD8⁺ T cells were detected at 27 days after KPC1199 implantation. Runnig model (**g**), and voluntary wheel running model (**h**). (n = 4 per group, 3 independent experiments). **i,** 3D images of inguinal lymph nodes with/without injection of 6-hydroxydopamine (6-OHDA) after 5 days. Scale bar: 200 μm ( 3 independent experiments). **j,** The level of intracellular NE in CD8⁺ T cells were detected at 5 days after inguinal lymph nodes injected with 6-OHDA (n = 4 per group, 3 independent experiments). **k-m,** KPC1199 tumor cells were subcutaneously implanted into mice. at the same time, sympathetic ablation was performed on the inguinal lymph nodes. Tumor volume size was monitored every 5 days (Ctrl: control, Ex: Running, Ex+6-OHDA: Running + 6-OHDA) (**k**), Percentage of CD8⁺ T (**l**) cells and intracellular GZMB of CD8⁺ T cells (**m**) were detected at 27 days after KPC1199 implantation (n = 4 per group, 3 independent experiments). **n,** LC–MS/MS analysis the absolute concentration of NE in tumor tissue, spleen, and various lymph nodes (n = 6 per group, 3 independent experiments). Statistical significance was determined by two-tailed unpaired Welcht’s *t*-test in a, b, c, d, e, f, g, h, j, k, l and m, Error bars: SEM; unpaired *t*-tests (two-tailed) in n. \**P* < 0.05, \*\**P* < 0.01, \*\*\**P* < 0.001; *N.S.*, not significant.

We next determined whether tumor control reflected changes in CD8⁺ T cell abundance versus effector readiness. Exercise did not alter the proportion of CD8⁺ T cells across the tumor, tdLNs, spleen, axillary lymph nodes, or peritoneal lymph nodes (Fig. 1e and f), arguing against increased expansion or infiltration as the primary driver. Instead, exercise selectively increased granzyme B (GZMB) protein levels in CD8⁺ T cells within the tumor and tdLNs, with no appreciable change in distal lymphoid sites (Fig. 1g and h). TNF-α and IFN-γ remained unchanged (Extended Data Fig.1d-g). Thus, exercise preferentially increases the available cytotoxic effector pool, rather than CD8⁺ T cell number or inflammatory cytokine outpu, in a compartment-restricted manner.

The spatial restriction of GZMB elevation to the tumor and its tdLNs suggested a tdLNs-local input. Because sympathetic activation is a rapid physiological consequence of exercise, we asked whether tdLNs-local sympathetic innervation is required for the increase in CD8⁺ T cell GZMB reserves. Regional chemical sympathectomy^29^ of the tdLNs with 6-hydroxydopamine (6-OHDA) (Fig. 1i and j) largely eliminated exercise-mediated tumor suppression (Fig. 1k and Extended Data Fig. 1h). Importantly, sympathectomy did not change tumor-infiltrating CD8⁺ T cell numbers (Fig. 1l) but significantly reduced GZMB protein levels in CD8⁺ T cells from both the tumor and tdLNs (Fig. 1m), while no significant effects were observed on the levels of TNF-α and IFN-γ (Extended Data Fig. 1i and j). These data indicate that tdLNs-local sympathetic input is necessary for elevating CD8⁺ T cell cytotoxic effector reserves.

To define a molecular correlation of this neural input, we quantified norepinephrine (NE) levels, the principal sympathetic neurotransmitter. Exercise increased NE levels across multiple tissues, with the strongest elevation occurring in the tdLNs (Fig. 1n), consistent with preferential tdLNs enrichment. Together, these results support a model in which tdLNs-local sympathetic NE signaling licenses CD8⁺ T cells to accumulate cytotoxic effector reserves.

### Norepinephrine selectively boosts CD8⁺ T cell GZMB reserves and cytotoxicity

Having implicated tdLNs-local NE as a candidate instructive signal, we next asked whether NE can directly enhance CD8⁺ T cell cytotoxic reserves. CD8⁺ T cells were activated *in vitro* under αCD3/αCD28 stimulation and IL-2 in the presence of NE. NE did not alter proliferation (Extended Data Fig. 2a) but markedly increased intracellular GZMB protein levels in CD8⁺ T cells, with 5 μM NE producing the strongest effect (Fig. 2a). In contrast, IFN-γ and TNF-α were not significantly changed (Extended Data Fig. 2b and c). Thus, NE preferentially increases the cytotoxic effector pool without broadly amplifying cytokine programs.

**Fig. 2.**
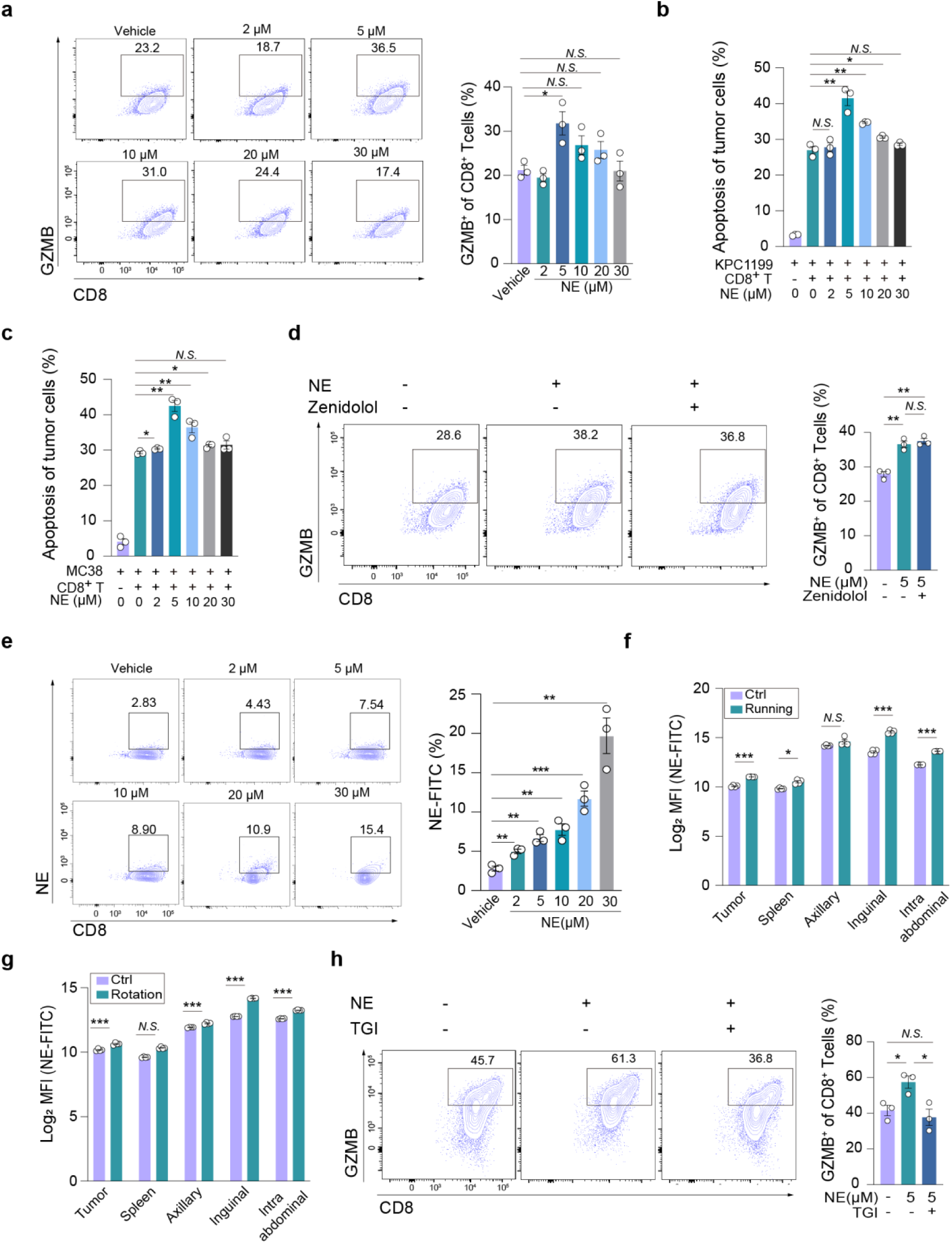
NE potentiates the anti-tumor activity of CD8^+^ T cells through the upregulation of GZMB level. **a,** CD8^+^ T cells were stimulated with IL-2 and αCD3/αCD28 for 72 hours, in the absence (Vehicle) or presence of 2, 5, 10, 20 or 30 μM norepinephrine (NE). Intracellular GZMB protein level was determined by flow cytometry. Left panels: Representative FACS dot plot images. Right panels: Bar graphs show the quantification of GZMB expression (n = 3 per group, 3 independent experiments). **b-c**, MACS-isolated spleen CD8^+^ T cells were stimulated with IL-2 and αCD3/αCD28, in the absence or presence of 2, 5, 10, 20 or 30 μM NE. After 72 hours activation, CD8^+^ T cells were co-cultured with KPC1199 or MC38 tumor cells for 24 hours, the apoptosis of KPC1199 tumor cells (**b**) and MC38 tumor cells (**c**) were measure by flow cytometry. (CD8^+^ T cells: tumor cells = 5:1; n = 3 per group; 3 independent experiments). **d**, CD8^+^ T cells were stimulated with IL-2 and αCD3/αCD28, pretreated with (+) or without (-) 10 μM β2 adrenergic receptor inhibitor Zenidolol. After 1 hour, 5 μM NE was added to continue stimulation for 72 hours, followed by flow cytometry to measure intracellular GZMB protein level. Left panel: Representative FACS dot plot images. Right panel: Bar graph showed the quantification of GZMB expression (n = 3 per group, 3 independent experiments). **e**, CD8^+^ T cells were stimulated with IL-2 and αCD3/αCD28 for 72 hours, in the presence or absence of 0, 2, 5, 10, 20 or 30 μM NE. Intracellular NE concentration were determined by flow cytometry. Left panels: Representative FACS dot plot images. Right panel: Bar graph showed the quantification of NE (n = 3 per group, 3 independent experiments). **f-g,** Flow cytometric analysis of NE levels in CD8⁺ T cells isolated from tumor, spleen, and lymph nodes. Results are shown for running (**e**) and voluntary wheel running (**f**) (n = 4 per group, 3 independent experiments). **h**, CD8^+^ T cells were stimulated with IL-2 and αCD3/αCD28, pretreated with (+) or without (-) 25 μM TGM2 inhibitor TGI. After 1 hour, 5 µM NE was added to continue stimulation for 72 hours, followed by flow cytometry to measure intracellular GZMB protein level. (n = 3 per group, 3 independent experiments). Statistical significance was determined by two-tailed unpaired Welcht’s *t*-test (a, b, c, d, e, f, g and h), Error bars: SEM. \**P* < 0.05, \*\**P* < 0.01, \*\*\**P* < 0.001; *N.S.*, not significant.

We next tested whether increased effector reserves translate into functional gain. NE treatment (5–20 μM) enhanced CD8⁺ T cell cytotoxicity against both KPC1199 and MC38 targets, with 5 μM showing the most robust improvement (Fig. 2b and c, and Extended Data Fig. 3a and b).

Mechanistically, NE can signal through cell-surface adrenergic receptors or through intracellular routes linked to protein post-translational modification. Given that β₂-adrenergic receptor (β₂-AR) is the predominant adrenergic receptor reported in CD8⁺ T cells^20^, we first evaluated β₂-AR dependence. The β₂-AR antagonist Zenidolol did not diminish NE-induced GZMB accumulation (Fig. 2d), confirming that canonical β₂-AR signaling is dispensable in this setting. We therefore examined NE uptake. CD8⁺ T cells showed a dose-dependent uptake of exogenous NE (Fig. 2e), and CD8⁺ T cells isolated from exercised mice across tumors and immune tissues displayed elevated intracellular NE levels (Fig. 2f and g).

Given that intracellular NE can act as a substrate for transglutaminase 2 (TGM2), a post-translational modification enzyme that catalyzes norepinephrinylation of protein substrates^23^, we next investigated the involvement of TGM2. Importantly, inhibition of TGM2 activity attenuated NE-induced GZMB accumulation (Fig. 2h). Together, these results link NE exposure to a receptor-independent, norepinephrinylation-associated route that increase CD8^+^ T cell GZMB reserves.

### GZMB norepinephrinylation as a post-translational mechanism stabilizing cytotoxic effector reserves in CD8^+^ T cells

Because NE enhances GZMB accumulation in CD8⁺ T cells, we next asked whether NE directly modifies GZMB via a post-translational mechanism. To systematically identify norepinephrinylation, we applied an alkyne-functionalized NE analog (PE) coupled with click chemistry (Fig. 3a), and confirmed that CD8^+^ T cells can take up exogenous PE via flow cytometry (Fig. 3b). LC–MS/MS following click conjugation and enrichment identified multiple PE-modified proteins in CD8⁺ T cells (Table S1), including GZMB, while β-actin, a known norepinephrinylation substrate^24^ was detected as a positive control. Western blotting further confirmed a PE-specific modification signal on GZMB (Fig. 3c).

**Fig. 3.**
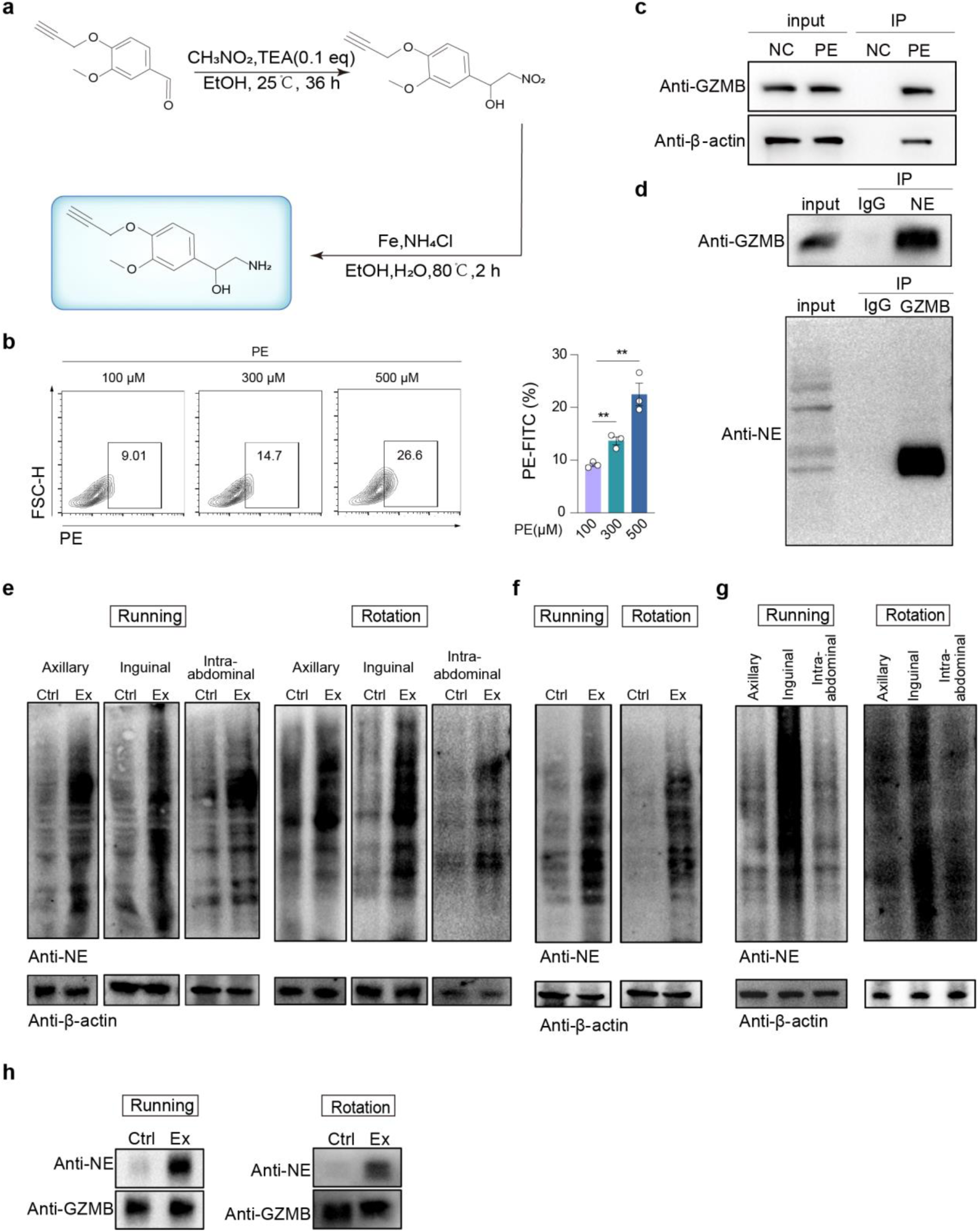
GZMB is norepinephrinylation protein. **a,** A chemical schematic diagram illustrating the synthesis of alkyne-modified NE. **b**, CD8^+^ T cells were stimulated with IL-2 and αCD3/αCD28 for 6 hours, in the presence of 100, 300, 500 μmol/L acetylenated NE (PE). Intracellular PE concentrations were determined by flow cytometry (n = 3 per group, 3 independent experiments). **c**, CD8^+^ T cells were stimulated with IL-2 and αCD3/αCD28 for 72 hours, then were cultured in the presence of 500 μM PE for 6 hours. Biotin-bonded PE was immunoprecipitated using click chemistry, GZMB and β-actin were detected with western blotting (3 independent experiments). IP, immunoprecipitation. **d**, CD8^+^ T cells were stimulated with IL-2 and αCD3/αCD28 for 72 hours. Anti-NE antibody or anti-GZMB antibody were immunoprecipitated, respectively. GZMB protein and GZMB norepinephrinylation were detected by western blot (3 independent experiments). IP, immunoprecipitation. **e-f**, Total proteins were extracted from immune cells of tumor tissues and lymph nodes under exercise and control conditions. Western blot analysis was performed to detect the norepinephrinylated protein in total protein from lymph node tissues (**e**), and from immune cells of tumor tissues (**f**). **g**, Total protein was isolated from axillary, inguinal, and mesenteric lymph nodes of exercised mice. Western blot was used to assess the norepinephrinylated protein levels across different lymphatic regions. **h**, Total proteins were extracted from lymph nodes under exercise and control conditions. GZMB protein was immunoprecipitated. The level of GZMB and GZMB norepinephrinylation were detected by western blot (3 independent experiments). Running (Left panels), voluntary wheel running (Right panels). IP, immunoprecipitation. Statistical significance was determined by two-tailed unpaired Welcht’s *t*-test (b), Error bars: SEM. \*\**P* < 0.01.

To establish GZMB as a bona fide target of norepinephrinylation, we performed reciprocal co-immunoprecipitation assays. NE immunoprecipitation followed by GZMB immunoblotting and the reciprocal GZMB immunoprecipitation followed by NE immunoblotting, together with targeted LC-MS/MS validation, robustly confirmed NE-associated modification on GZMB (Fig. 3d and Table S2).

We next asked whether norepinephrinylation is dynamically regulated *in vivo*. Exercise increased global norepinephrinylation in immune cells from lymph nodes and tumors (Fig. 3e and f), with the most pronounced enhancement in the tdLNs (Fig. 3g). Importantly, norepinephrinylation on GZMB itself was also significantly elevated under exercise conditions (Fig. 3h). Together, these *in vitro* and *in vivo* data support a model in which tdLNs-enriched NE promotes GZMB norepinephrinylation, favoring GZMB accumulation within CD8⁺ T cells.

### Attenuation of norepinephrinylation compromises GZMB stability and CD8⁺ T cell function

To genetically attenuate NE-associated covalent modification in CD8⁺ T cells, we generated CD8⁺ T cell–specific *Tgm2* conditional knockout mice (*Tgm2*^fl/fl^; CD8-Cre)^30^. In both KPC1199 and MC38 tumor models, *Tgm2*^fl/fl^-CD8-Cre mice exhibited accelerated tumor growth compared with littermate controls (Fig. 4a-d). Consistent with impaired anti-tumor immunity, CD8⁺ T cells isolated from these mice displayed reduced cytotoxic activity in *ex vivo* killing assays (Fig. 4e and Extended Data Fig. 3c), indicating that disrupting norepinephrinylation compromises CD8⁺ T cell effector function.

**Fig. 4.**
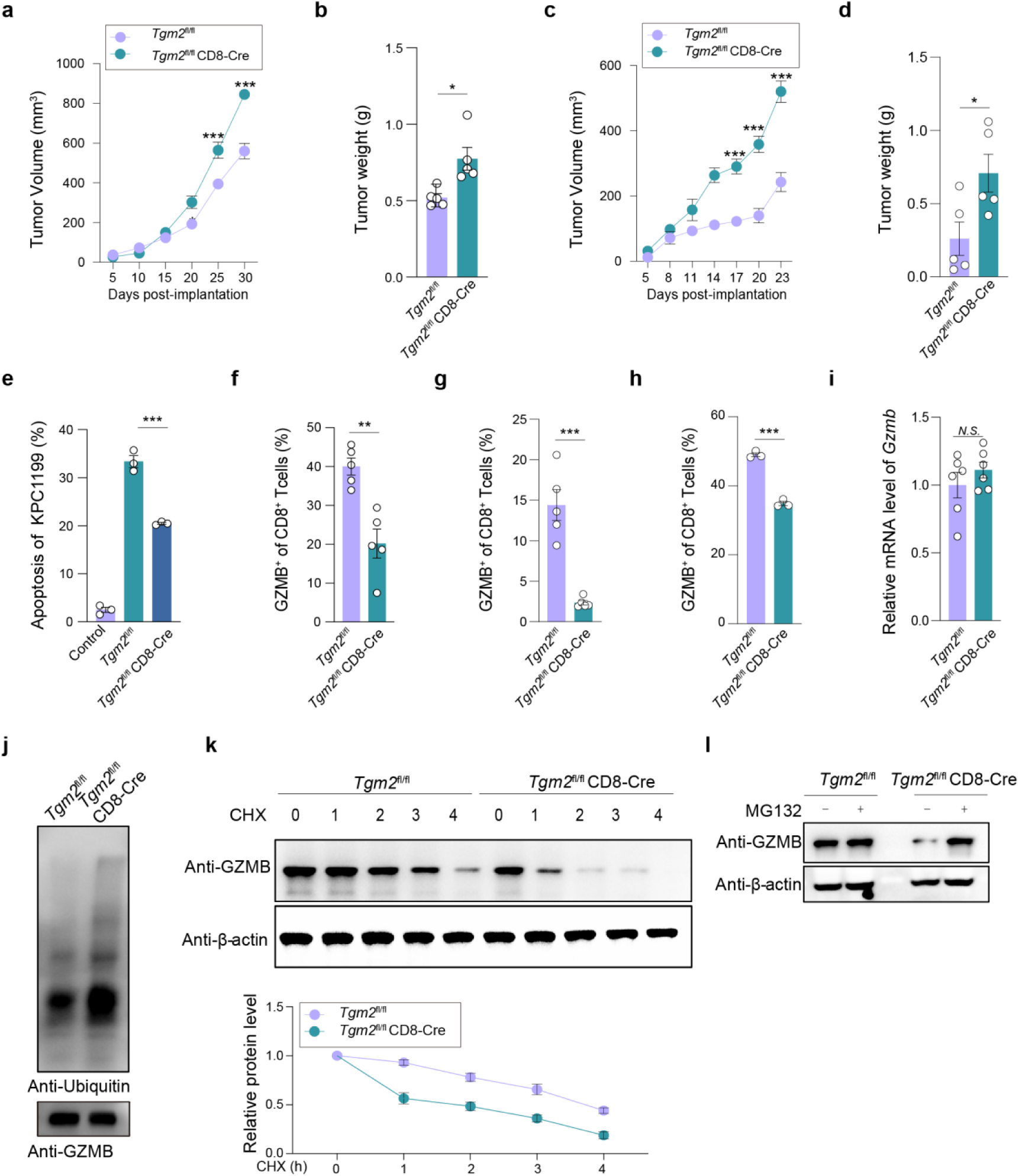
Attenuation of norepinephrinylation compromises GZMB stability and CD8⁺ T-cell function. **a-b**, KPC1199 tumor cells were subcutaneously implanted into syngeneic *Tgm2*^fl/fl^ or *Tgm2*^fl/fl^CD8-Cre mice. Tumor volume (**a**) was monitored every 5 days, and tumor weight (**b**) was measured at 30 days after implantation (n = 5 per group, 3 independent experiments). **c-d,** MC38 tumor cells were subcutaneously implanted into syngeneic *Tgm2*^fl/fl^ or *Tgm2*^fl/fl^CD8-Cre mice. Tumor volume (**c**) was monitored every 3 days, and tumor weight (**d**) was detected at the 23 days after implantation (n = 5 per group, 3 independent experiments). **e**, CD8^+^ T cells from *Tgm2*^fl/fl^ or *Tgm2*^fl/fl^CD8-Cre mice were stimulated with IL-2 and αCD3/αCD28 for 72 hours. After 72 hours, CD8^+^ T cells were co-cultured with KPC1199 tumor cells for 24 hours, and then flow cytometry was used to assess the apoptosis in KPC1199 tumor cells (n = 3 per group, 3 independent experiments). **f-g,** Frequency of intracellular GZMB in CD8^+^ T cells was detected from the tumor tissues collected at 30 or 23 days after implantation with KPC1199 cells (**f**) and MC38 cells (**g**) (n = 5 per group, 3 independent experiments). **h**, *Tgm2*^fl/fl^ or *Tgm2*^fl/fl^CD8-Cre CD8^+^ T cells were stimulated with IL-2 and αCD3/αCD28 for 72 hours. Intracellular GZMB were determined by flow cytometry (n = 3 per group, 3 independent experiments). **i,** *Tgm2*^fl/fl^ or *Tgm2*^fl/fl^CD8-Cre CD8^+^ T cells were stimulated with IL-2 and αCD3/αCD28 for 72 hours. The mRNA levels of GZMB were analyzed by quantitative real-time PCR (qRT-PCR) (n = 6 per group, 3 independent experiments). **j,** *Tgm2*^fl/fl^ or *Tgm2*^fl/fl^CD8-Cre CD8^+^ T cells were stimulated with IL-2 and αCD3/αCD28 for 72 hours. GZMB protein was immunoprecipitated, and the ubiquitination levels of GZMB was analyzed by western blotting. GZMB was set as control (3 independent experiments). **k,** *Tgm2*^fl/fl^ or *Tgm2*^fl/fl^CD8-Cre CD8^+^ T cells were stimulated with IL-2 and αCD3/αCD28 for 72 hours. After 72 hours cells were treated with inhibitor cycloheximide (CHX), and the GZMB protein levels were detected by western blot at 0, 1, 2, 3 and 4 hours after CHX treatment, respectively. Representative western blot images were shown, β-actin was set as control and the graph shown the relative intensity (3 independent experiments). **l.** *Tgm2*^fl/fl^ or *Tgm2*^fl/fl^CD8-Cre CD8^+^ T cells were stimulated with IL-2 and αCD3/αCD28 for 72 hours with/without MG-132 treatment. After 72 hours, GZMB protein levels in *Tgm2*^fl/fl^ or *Tgm2*^fl/fl^CD8-Cre CD8^+^ T cells with/without MG-132 treatment were detected by western blot. β-actin was set as control (3 independent experiments). Statistical significance was determined by two-tailed unpaired Welcht’s *t*-test (a, b, c, d, e, f, g, h and i), Error bars: SEM. \**P* < 0.05, \*\**P* < 0.01, \*\*\**P* < 0.001; *N.S.*, not significant.

We next investigated the molecular basis of this defect. GZMB protein levels were reduced in tumor-derived CD8⁺ T cells and were similarly decreased in spleen-derived CD8⁺ T cells isolated from *Tgm2*^fl/fl^ -CD8-Cre mice after in vitro reactivation (Fig. 4f-h and Extended Data Fig. 3d). In contrast, *Gzmb* mRNA abundance was unchanged (Fig. 4i), suggesting a post-transcriptional mechanism.

We therefore focused on GZMB stability. GZMB must be rapidly packaged into secretory granules; unincorporated cytosolic GZMB is ubiquitinated and cleared via the proteasome. In line with this model, attenuation of norepinephrinylation increased GZMB ubiquitination (Fig. 4j) and accelerated GZMB decay in CHX chase assays (Fig. 4k). Proteasome inhibition with MG132 fully restored GZMB protein levels (Fig. 4l), indicating that loss of norepinephrinylation renders GZMB more susceptible to ubiquitin–proteasome–mediated degradation. Together, these data demonstrate that norepinephrinylation preserves GZMB protein stability, thereby sustaining cytotoxic effector reserves and tumor control.

### GZMB Gln43 as the NE-responsive determinant of UHRF1-dependent ubiquitination and protein stability

To define the modification site(s), we reconstituted an *in vitro* norepinephrinylation reaction and mapped modified residues on GZMB peptides by LC–MS/MS using either MDC (a broadly reactive amine donor) or NE as substrate. MDC modified multiple glutamine residues (Extended Data Fig. 4a-e), whereas NE showed striking selectivity and modified only Gln43 under identical conditions (Fig. 5a), identifying Gln43 as a dominant site for norepinephrinylation.

**Fig. 5.**
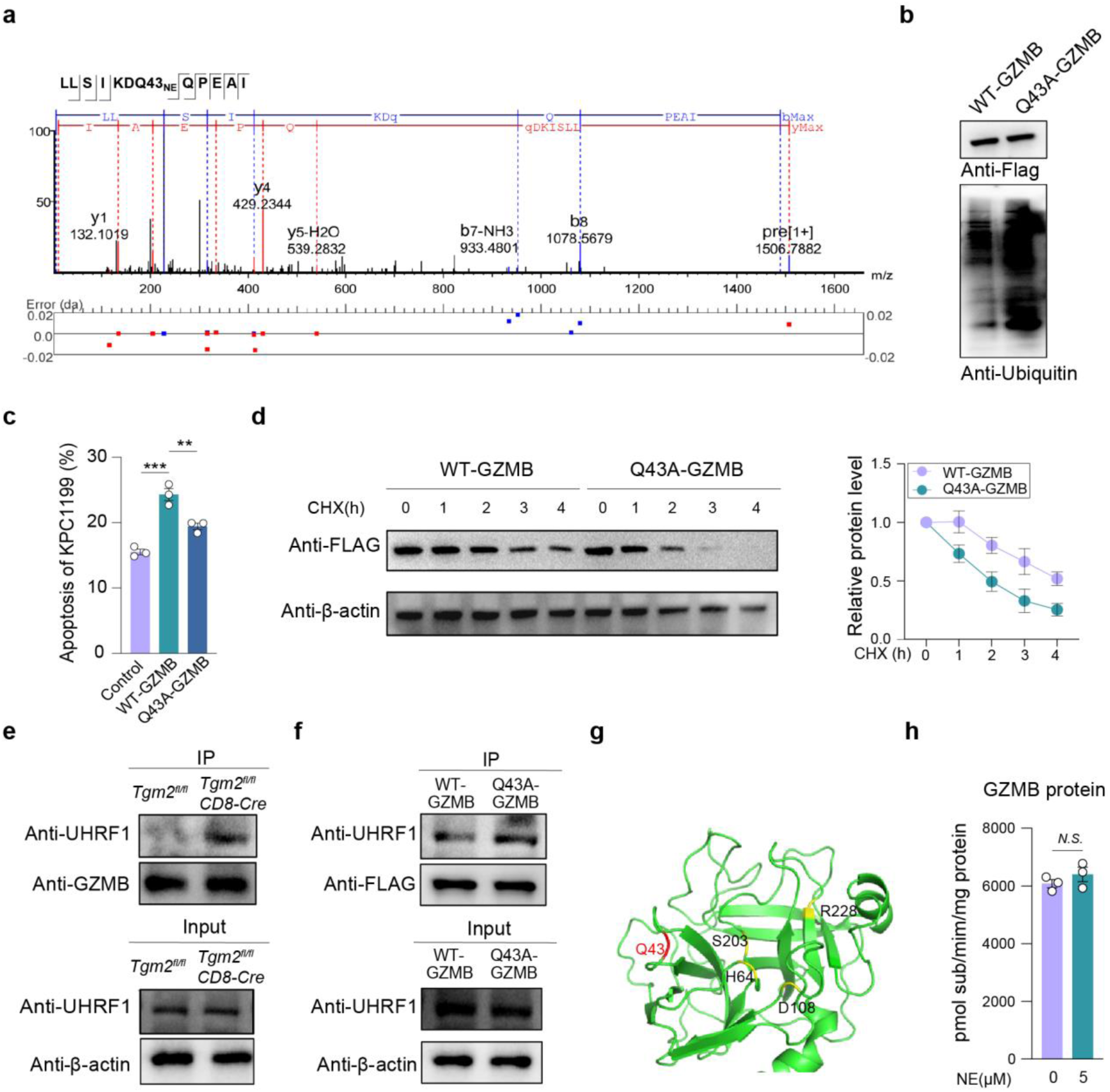
The glutamine residue Gln43 of GZMB is norepinephrinylated and regulates UHRF1 to recognize GZMB. **a,** Targeted liquid chromatography-tandem mass spectrometry (LC-MS/MS) analysis of a TGM2-transamidated NE to GZMB peptide 37–48. **b,** EL4 cells were transfected with Flag-fused lentiviruses expressing either WT-GZMB or Gln43 to Ala (Q43A) mutated GZMB (Q43A-GZMB). Cells were stimulated with IL-2 and αCD3/αCD28 for 72 hours, Flag-beads were used to pull down Flag-GZMB, and the ubiquitination level of GZMB protein were detected in EL4 cells with overexpression of WT-GZMB or Q43A-GZMB (3 independent experiments). **c,** EL4 cells were stimulated with IL-2 and αCD3/αCD28 for 72 hours. After 72 hours, EL4 cells were co-cultured with KPC1199 tumor cells for 24 hours, and then flow cytometry was used to assess the levels of apoptosis in KPC1199 tumor cells. As a control, tumor cells incubation with wide-type EL4 cells were subjected to FACS analysis. Bar graph showed the quantification of apoptosis (EL4 cells: tumor cells = 5:1; n=3 per group; 3 independent experiments). **d,** EL4 cells were stimulated with IL-2 and αCD3/αCD28 for 72 hours. After 72 hours, cells were treated with inhibitor CHX at 0, 1, 2, 3 and 4 hours, and WT-GZMB or Q43A-GZMB protein levels were detected by western blot respectively. Left panels: Representative western blot images, β-actin was set as control. Right panels: Graph showed the relative intensity (3 independent experiments). **e,** Upper panels: *Tgm2*^fl/fl^ or *Tgm2*^fl/fl^CD8-Cre CD8^+^ T cells were stimulated with IL-2 and αCD3/αCD28 for 72 hours, GZMB was immunoprecipitated and UHRF1 protein binding to GZMB was detected by western blot. GZMB was set as control. Lower panels: UHRF1 protein levels in *Tgm2*^fl/fl^ or *Tgm2*^fl/fl^CD8-Cre CD8^+^ T cells were detected by western blot. β-actin was set as control, IP, immunoprecipitation (3 independent experiments). **f,** Upper panels: EL4 cells were stimulated with IL-2 and αCD3/αCD28 for 72 hours, Flag-GZMB was immunoprecipitated and UHRF1 protein binding to GZMB were detected by western blot. GZMB was set as control. Lower panels: UHRF1 protein levels in WT-GZMB (EL4) or Q43A-GZMB (EL4) cells were detected by western blot. β-actin was set as control, IP, immunoprecipitation (3 independent experiments). **g,** A ribbon trace of GZMB with labeled surface loops. The locations are shown for the catalytic residues, H64, D108, S203, and R228 (in yellow), and site of norepinephrinylation amino acids Q43 (in red). This model was from Uniprot (AF-P04187-F1-V4). **h,** TGM2 monoaminylation assay with GZMB protein in the absence (Vehicle) or presence of 5 μM NE. Activity of norepinephrinylated GZMB was analyzed. The immediately fluorescenceintensity at Ex/Em = 380/500 nm was measured in a microplate reader in kinetic mode for 30 min at 37°C protected from light (n = 3 per group, 3 independent experiments). Statistical significance was determined by two-tailed unpaired Welcht’s *t*-test (c, h), Error bars: SEM. \*\**P* < 0.01; \*\*\**P* < 0.001; *N.S.*, not significant.

To test functional necessity, we expressed wild-type GZMB (WT-GZMB) or a Gln43-Ala mutant (Q43A-GZMB) in EL4 T cells. Upon αCD3/αCD28 stimulation, Q43A-GZMB showed increased ubiquitination (Fig. 5b), reduced killing of KPC1199 cells (Fig. 5c and Extended Data Fig. 4f), and accelerated degradation in CHX chase assays (Fig. 5d), establishing Gln43 as a key determinant of GZMB stability and cytotoxic fitness.

Given the ubiquitination phenotype, we next sought candidate E3 ligases. LC–MS/MS of GZMB immunoprecipitants identified multiple candidates (Table S3 and Extended Data Fig. 5g), with UHRF1 being the most abundant. Co-immunoprecipitation confirmed increased UHRF1 association with GZMB in *Tgm2*-deficient CD8⁺ T cells (Fig. 5e) and stronger binding of UHRF1 to Q43A-GZMB than WT-GZMB (Fig. 5f), suggesting that the Gln43 state controls UHRF1 recognition, thereby tuning ubiquitination and stability. Furthermore, Gln43 lies outside the catalytic H64, D108, S203, and R228 core^31^ (Fig. 5g), and NE treatment did not alter GZMB enzymatic activity in vitro (Fig. 5h). Collectively, these results indicate that norepinephrinylation at Gln43 stabilizes GZMB by limiting UHRF1-mediated ubiquitination and proteasomal degradation, without affecting catalytic function.

### *Ex vivo* norepinephrine conditioning as a minimal strategy to enhance adoptive CD8⁺ T cell tumor control

Mechanistically defining the NE-GZMB stability axis suggested a reductionist translational strategy: whether a brief *ex vivo* NE exposure during activation could pre-load CD8⁺ T cells with stabilized GZMB reserves and thereby improve adoptive tumor control. CD8⁺ T cells from CD45.1⁺ donors were activated for 72 h with αCD3/αCD28 antibodies and IL-2 in the presence of 5 μM NE, and then transferred into CD45.2⁺ recipients bearing established KPC1199 tumors (day 7; Fig. 6a). NE-conditioned CD8⁺ T cells significantly suppressed tumor growth, reducing both tumor volume and weight (Fig. 6b and c), demonstrating that a minimal *ex vivo* conditioning step is sufficient to enhance *in vivo* efficacy.

**Fig. 6.**
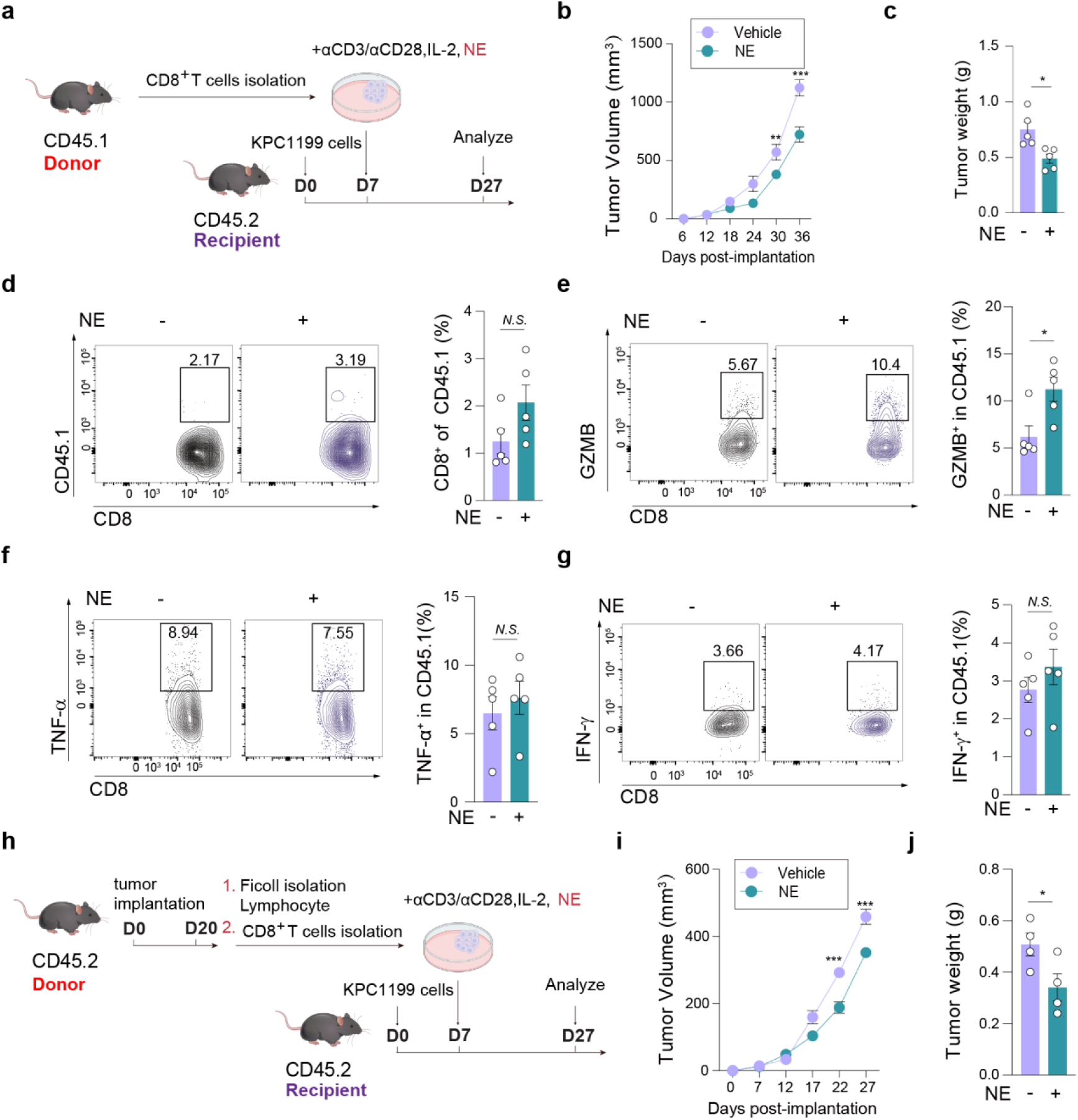
NE enhances the anti-tumor ability of CD8^+^ T cells in adoptive therapy. **a,** Experimental set up: CD8^+^ T cells from the spleen of CD45.1 mice were stimulated with IL-2, IL-2 and αCD3/αCD28 for 72 hours, in the absence (Vehicle) or presence of 5 μM NE. After 72 hours, CD45.1^+^ CD8^+^ T cells (donor cells) were injected into CD45.2 recipient mice, which were subjected to subcutaneous injection of KPC1199 cells 7 days prior. **b-c**, Tumor volume (**b**) was monitored every 5 days, and tumor weight (**c**) was measured at 27 days after implantation (n = 5 per group, 3 independent experiments). **d,** The frequency CD8^+^ T cells in donor CD45.1^+^ cells were detected at 27 days after implantation (n = 5 per group, 3 independent experiments). **e,**The intracellular GZMB in CD45.1^+^CD8^+^ T cells were detected at 27 days after implantation (n = 5 per group, 3 independent experiments). **f-g,** The intracellular TNF-α (**f**), and IFN-γ (**g**) in CD45.1^+^CD8^+^ T cells were detected at 27 days after implantation (n = 5 per group, 3 independent experiments). **h,** Experimental set up: KPC1199 tumor cells were subcutaneously implanted into C57BL/6 mice. CD8^+^ T cells from the tumor tissues at 20 days after implantation were purified, and the cells were stimulated with IL-2 and αCD3/αCD28 in vitro for 72 hours, in the absence (Vehicle) or presence of 5 μM NE. After 72 hours, cells were injected into C57BL/6 mice, which were subjected to subcutaneous injection of KPC1199 cells 7 days prior. **i-j**, Tumor volume (**i**) was monitored every 5 days, and tumor weight (**j**) was measured at 27 days after implantation (n = 4 per group, 3 independent experiments). Statistical significance was determined by two-tailed unpaired Welcht’s *t*-test ( b, c, d, f, g, i and j), Error bars: SEM. \**P* < 0.05; \*\**P* < 0.01. *N.S.*, not significant.

We next asked whether this benefit reflected altered accumulation versus altered effector reserves in situ. NE conditioning did not change the intratumoral frequency of donor-derived (CD45.1⁺) or recipient-derived (CD45.2⁺) CD8⁺ T cells (Fig. 6d and Extended Data Fig. 5a). Instead, NE conditioning selectively increased GZMB expression in donor-derived CD8⁺ T cells (Fig. 6e) without altering TNF-α or IFN-γ (Fig. 6f and g), and recipient-derived CD8⁺ T cells remained unchanged (Extended Data Fig. 6b-d). Thus, NE conditioning preferentially strengthens the cytotoxic effector module rather than inducing non-specific expansion or inflammatory rewiring.

Finally, we tested generalizability across T cell states. Tumor-infiltrating CD8⁺ T cells harvested from day-20 tumor-bearing mice, when subjected to the same *ex vivo* NE conditioning and transferred, likewise mediated significant tumor suppression (Fig. 6h-j). Together, these results demonstrate that once the decisive NE-GZMB stability mechanism is defined, both peripheral and tumor-experienced CD8⁺ T cells can be potentiated through a simple, controllable NE-based *ex vivo* intervention.

## Discussion

High intracellular stores of granzyme B (GZMB) enable cytotoxic T lymphocytes (CTLs) to deliver rapid, high-amplitude effector output immediately upon target recognition, a kinetic advantage that is essential for controlling fast-replicating viruses and eliminating nascent malignant cells^32,33^. At the same time, excessive or misdirected cytotoxic effector release can damage healthy tissues and have been implicated in immunopathology^34^, including autoimmune diseases such as type 1 diabetes^35^, multiple sclerosis^36^, and graft-versus-host disease^37^. These considerations suggest that CTLs must operate within a tightly regulated window that maximizes pathogen/tumor clearance while minimizing collateral injury. Here, we uncover a physiological mechanism that tunes this balance by controlling the size of the cytotoxic effector reserve, rather than simply altering T cell number or inflammatory cytokine programs. A central finding of this study is that tdLNs-local sympathetic neurotransmission instructs CD8⁺ T cell cytotoxic programming by regulating GZMB protein homeostasis during early priming. While canonical models emphasize antigen load, TCR signal strength, costimulation, and cytokines as primary determinants of effector differentiation^38,39^, these signals do not readily explain why effector T cells can display substantial heterogeneity in cytotoxic effector reserves even under similar antigenic conditions. Our data argue that an additional layer of control operates at the level of effector protein fate: within the critical time-and-space window of the dLNs, sympathetic input dynamically adjusts the amount of cytotoxic machinery that CD8⁺ T cells carry forward into peripheral tissues. Conceptually, this reframes “effector programming” not only as transcriptional specification, but also as post-translational allocation and stabilization of effector proteins that define the functional ceiling of cytotoxicity.

Mechanistically, we identify a non-canonical NE pathway that extends beyond adrenergic receptor signaling. We show that NE can enter CD8⁺ T cells and promote a covalent post-translational modification of GZMB, norepinephrinylation at Gln43, catalyzed by transglutaminase 2 (TGM2). This modification protects GZMB from UHRF1-dependent ubiquitination and proteasomal degradation, thereby extending GZMB protein half-life and enabling its accumulation as an effector reserve. Importantly, this pathway regulates cytotoxic capacity without broadly increasing pro-inflammatory cytokine output, indicating that sympathetic NE can selectively potentiate the cytotoxic module rather than globally amplifying T cell activation. By linking a tdLNs-enriched neural cue to a defined post-translational mechanism that stabilizes a core effector protein, our findings revise the prevailing view of how the nervous system shapes adaptive immunity, highlighting effector protein homeostasis as a neuro-immune checkpoint.

Our results also illuminate an important physiological principle: NE acts within a concentration window. NE enhanced GZMB accumulation and cytotoxicity at defined doses, whereas higher levels attenuated these effects. This dose dependence suggests an intrinsic safeguard that couples enhanced immune efficacy to self-protection, and consistent with the broader notion that moderate physiological stimulation can be beneficial whereas excessive stimulation may be counterproductive. In the context of sympathetic immunoregulation, such a “Goldilocks” regime could help explain why physiological states (e.g., exercise versus severe stress) can have divergent immunological outcomes despite sharing catecholaminergic features.

Beyond conceptual advances, our study provides a reductionist translational strategy. Because the NE-GZMB stability axis operates at the level of effector reserves, it can be harnessed *ex vivo* without reproducing the full complexity of the *in vivo* neuro-immune environment. Brief NE exposure during activation was sufficient to increase GZMB reserves and improve tumor control by adoptively transferred CD8⁺ T cells, including both splenic CD8⁺ T cells and tumor-infiltrating T cells. These findings suggest that controlled NE conditioning may represent a practical approach to enhance adoptive cell therapy (ACT) by reinforcing cytotoxic effector readiness. More broadly, they point to neuro-immune modulation of effector protein stability as an actionable lever to improve antitumor immunity, potentially complementing strategies aimed at reversing exhaustion or improving trafficking.

Several questions remain. First, our *in vivo* work relies primarily on subcutaneous tumor models, where neuroanatomical organization may differ from that of orthotopic or metastatic tumors. Testing the generality of the tdLNs-local NE-GZMB axis across additional tumor types, anatomical sites, and human samples will be important. Second, catecholamine biology is highly context- and dose-dependent; how norepinephrinylation integrates with adrenergic receptor signaling, metabolic state, and inflammatory cues across diverse physiological and pathological settings requires systematic investigation. Third, the structural basis by which GZMB Gln43 norepinephrinylation limits UHRF1 recognition remains to be resolved, whether through steric effects, conformational shifts, or interplay with other post-translational modifications. Finally, lymph nodes are spatially compartmentalized, and sympathetic innervation is heterogeneous; defining how regional neural patterns within distinct nodal niches shape CD8⁺ T cell fate decisions, and whether analogous mechanisms operate in other lymphoid organs or during responses to different classes of antigens, will further refine this model.

In summary, our work uncovers a cross-system regulatory pathway linking tdLNs-local sympathetic NE to CTL effector quality through post-translational stabilization of GZMB. By establishing the NE-GZMB axis as a determinant of cytotoxic effector reserves, we expand current paradigms of effector programming and provide a mechanistically grounded, minimally invasive strategy to potentiate CD8⁺ T cell mediated tumor control.

## Materials and methods

### Cell lines

KPC1199 Pancreatic cancer cell line was generously provided by Prof. Jing Xue (Ren Ji Hospital, School of Medicine, Shanghai Jiao Tong University). MC38 colorectal cancer cell line was purchased from Cell bank of Chinese Academy of Sciences. EL4 cells were purchased from ATCC. KPC1199 and MC38 cells were cultured in Dulbecco’s modified Eagle’s medium (DMEM) and EL4 cells were cultured in RPMI 1640, supplemented with 10% fetal bovine serum (FBS) and penicillin and streptomycin (100 U/ml each) at 37°C and 5% CO_2_.

### Mouse strains

*Tgm2*^fl/fl^ mice were purchased from Cyagen Biosciences. *Cd8a-*cre, CD45.1 and wild type (WT) C57BL/6 mice were purchased from GemPharmatech Co., Ltd. Mice were bred and housed according to protocols approved by the Institutional Animal Care and Use Committee (IACUC) of Shanghai Jiao Tong University (the Animal Protocol number is A2023083), and were used between the ages of 6-8 weeks. All animal experiments were conducted in accordance with the National Institutes of Health Guide for the Care and Use of Laboratory Animals.

### Animal studies

#### Subcutaneous tumor model

KPC1199 cells (1 × 10^6^) or MC38 cells (0.5 × 10^6^) were suspended in 100 μL PBS and subcutaneously injected into the right flank of syngeneic *Tgm2*^fl/fl^ and *Tgm2*^fl/fl^CD8-Cre mice. Tumor growth was monitored with an electronic calliper by measuring tumor size every 5 days (KPC1199) or 3 days (MC38). Tumor volumes were calculated as 1/2 × Length × Width^2^.

#### Treadmill and Voluntary wheel running protocols

C57BL/6J WT male mice were used in all experiments. Mice were randomly divided into two groups: Control (ctrl) and Exercise (Ex). The treadmill test was performed starting with 7-week-old mice, exercised mice were involuntarily placed on a Rodent 6-lane treadmill (Shanghai XinRuan Information Technology Co., Ltd, Cat No: XR-PT-10B), for 30 min per day at 15 cm/sec^40^, for a minimum of 5 days/week. Rotated exercises were provided by running wheels (diameter 12.7 cm) (Lafayette Instrument Company, Lafayette, USA, Cat No: 80822S). The mice were inoculated subcutaneously with 1 × 10^6^ KPC1199 tumor cells. Exercise interventions (treadmill and voluntary wheel running) commenced 5 days following tumor cell implantation.

#### Identification of Draining Lymph Nodes

Mice were randomly divided into a control group and Evans blue group. Mice were subcutaneously injected into the subcutaneous tumor with a 1% (w/v) solution of Evans blue prepared in sterile PBS. Mice were euthanized by CO_2_ and cervical dislocation after 15 min, to allow for lymphatic trafficking of the dye.

#### Sympathetic denervation with 6-OHDA

For targeted sympathetic denervation of the mouse inguinal lymph nodes, 20 μL of 100 mg/kg 6-hydroxydopamine (6-OHDA) (MCE) or PBS was administered every 7 days. Following anesthesia, the skin was incised to expose the inguinal lymph nodes, and the injections were delivered into the adipose tissue adjacent to the lymph nodes. The initial injection coincided with tumor cell inoculation. 6-OHDA administration was performed until the experimental endpoint.

#### Adoptive transfer experiments

Magnetically activated cell sorting (MACS)-isolated CD8^+^ T cells from C57/BL6 (CD45.1) mouse spleen or KPC1199 solid tumors were stimulated with IL-2, αCD3/αCD28 for 72 hours, in the presence or absence of 5 μmol/L NE. C57/BL6 (CD45.2) mice were implanted with KPC1199 tumor cells. After 7 days, mice were intravenously injected with 3 × 10^5^ CD8^+^ T cells that had been activated *in vitro*. Tumor growth was monitored with an electronic calliper by measuring tumor size every 5 days. Tumor volumes were calculated as 1/2 × Length × Width^2^.

#### Mice tissues sampling and biochemical analysis

Mice were euthanized by carbon dioxide inhalation, and their tumor tissues, axillary lymph nodes, inguinal lymph nodes, intra-abdominal lymph nodes (including mesenteric and renal hilar/para-aortic lymph nodes), serum, and spleen samples were immediately frozen in liquid nitrogen and stored at -80℃. The levels of monoamine neurotransmitters were measured by Metabo-Profile BiotechnologyCo., Ltd (Shanghai, China).

Briefly, a 10 μL aliquot of serum was transferred to a centrifuge tube, followed by the addition of 70 μL of methanol containing internal standards. The mixture was vortexed at 10°C and 1400 rpm for 20 min. After centrifugation at 18,000 ×g and 4°C for 20 min, 55 μL of the supernatant was collected into a 96-well plate and vacuum-dried. Then, 50 μL of Phenylisothiocyanate (PITC) derivatization solution was added, and the reaction was carried out at 30°C for 30 min. The solution was dried under nitrogen flow, reconstituted with 200 μL of methanol containing 5 mmol/L ammonium acetate, and vortexed for 30 min. Following centrifugation for 10 min, 50 μL of the supernatant was transferred to a new 96-well plate, mixed with 50 μL of deionized water, and prepared for injection.

Exactly 10 mg of tissue sample was weighed into a centrifuge tube, to which 30 μL of deionized water and 10 grinding beads were added, followed by homogenization for 2 min. Then, 150 μL of methanol containing internal standards was added, and homogenization continued for another 3 min. The mixture was centrifuged at 18,000 ×g and 4°C for 20 min. A 20 μL aliquot of the supernatant was transferred to a 96-well plate and lyophilized. Subsequently, 50 μL of PITC derivatization solution was added, and the derivatization proceeded at 30°C for 30 min. The sample was dried under nitrogen, reconstituted with 200 μL of methanol containing 5 mmol/L ammonium acetate, and vortexed for 30 min. After centrifugation for 10 min, 50 μL of the supernatant was collected into a new 96-well plate, mixed with 50 μL of deionized water, and prepared for injection.

Analysis was performed using an ultra-performance liquid chromatography-tandem mass spectrometry (UPLC-MS/MS) system (ACQUITY UPLC-Xevo TQ-S, Waters, USA) under the following conditions: a C18 analytical column (2.1 × 100 mm, 1.7 μm) maintained at 45°C; mobile phase A: aqueous solution with 0.1% formic acid, mobile phase B: acetonitrile/methanol solution; flow rate: 0.4 mL/min; capillary voltage: 3 kV, following the manufacturer’s recommended protocol.

#### *In vivo* cell isolation and flow cytometry analyses

Mice were euthanized and fresh tissues were collected. Spleens were mechanically disrupted through disrupted through 70 μm Nylon mesh to release single cells. Tumor tissue was cut into small pieces and digested with 1mg/ml collagenase A (Sigma) and 1× DNase I (Sigma) for 20 min. Digestion was terminated with 3-5 mL medium containing 5% FBS. The liquid with fragments was filtered with 70 μm nylon filter and washed with PBS, and then lymphocytes were isolated with Ficoll gradient.

For staining molecules expressed on cell surface, single cell suspensions from spleens, was blocked with anti-mouse CD16/CD32 antibodies (BD Biosciencs) for 10 min and labeled with fluorophore-conjugated antibodies: PerCP/Cyanine5.5 anti-mouse CD45.1 (Biolegend, 1:200), PE anti-mouse CD45.2 (Biolegend, 1:200), and PE-Cy7 anti-mouse CD8 (Biolegend, 1:200).

To enrich immunocytes in tumors, cells were centrifuged through a 37.5% ficoll (cytiva) at 600 × g for 35 min. For intracellular staining, single cell suspensions from tumor tissues were stimulated with leukocyte activation cocktail (BD Biosciencs) for 4 hours, and labeled with PerCP/Cyanine5.5 anti-mouse CD45.1 (Biolegend, 1:200), PE anti-mouse CD45.2 (Biolegend, 1:200), and PE-Cy7 anti-mouse CD8 (Biolegend, 1:200). After fixation and permeabilization using the Fixation/Permeablization Kit (BD Biosciencs) following the manufacturer’s instructions, cells were labeled with APC anti-human/mouse GZMB, BV421 anti-mouse IFNγ (Biolegend, 1:200), and BV605 anti-mouse TNFα (Biolegend, 1:200). Flow cytometry was performed in BD FACSCanto II.

#### *In vitro* CD8^+^ T cells activation and flow cytometry analyses

CD8^+^ T cells were isolated from mouse spleen by CD8^+^ magnetic bead selection (Miltenyi Biotec). IL-2 (Biolegend, 10 ng/mL), αCD3 (Biolegend, 5 μg/mL) and αCD28 (Biolegend, 2 μg/mL) antibodies were used to activate the naïve CD8^+^ T cells or EL4 cells in RPMI 1640, supplemented with 10% fetal bovine serum (FBS) (Dcell biologics) at 37°C and 5% CO_2_. Cells were treated with NE (MCE) at indicated concentrations in the presence or absence of TGMs inhibitor TGI (LDN-27219 (MCE)), DBH inhibitor Fusaric acid (MCE), and β2AR inhibitor Zenidolol (MCE, 10 μmol/mL). For proliferation studies, CD8^+^ T cells were labelled with 1 μmol/L carboxyfluorescein succinimidyl ester (CFSE, Invitrogen) and then activated by IL-2 and αCD3/αCD28. The proliferation of CD8^+^ T cells was detected with flow cytometry and analyzed by CFSE dilution. For intracellular staining, cells were treated with leukocyte activation cocktail (BD Biosciencs) for 30 min and then fixed and permeabilized using the Fixation/Permeablization Kit (BD Biosciencs) following the manufacturer’s instructions. APC anti-mouse GZMB (BioLegend, 1:200), and anti-NE antibody (LSbio, 1:200) followed by secondary antibody (Servicebio, 1:500) was performed for intracellular staining.

#### Co-culture of CD8^+^ T cells and tumor cells for tumor cell apoptosis analysis

CD8^+^ T cells were labelled with 1 μmol/L cell Trace Violet (Invitrogen) and then activated by IL-2 and αCD3/αCD28. Approximately 1.5 × 10^4^ KPC1199 cells/well were cultured in a 24-well plate for 24 hours. CD8^+^ T cells (effector cells) were cocultured with target cells (KPC1199) at certain effector-target (E: T=5:1) ratios as indicated for 24 hours. After 24 hours, the cells were washed twice by PBS. In order to dissociate the adhered cells, 0.05% trypsin–EDTA was used. After removing trypsin–EDTA by centrifugation, the binding buffer was mixed with the cell pellet. 5 μL of both annexin V-FITC and PI (Share-bio) were added into the cell solution. After 15 min of incubation, stained cell solution was diluted with 400 μL of the binding buffer to the final concentration of approximately 1 × 10^4^ cells/well and fed to the FACS system.

#### Stable transfection of EL4 cells

EGFP-FLAG-fused lentiviruses expressing WT-GZMB or Gln to Ala (Q43A) mutated GZMB were constructed by GenePharma. EL4 cells were transfected at MOI 50. 6 μg/mL puromycin was added to the culture medium for 48 hours for selection. After 10 days of culture, the cell expressing EGFP were sorted using a BD FACSAria instrument.

#### Quantitative real-time PCR (qRT-PCR)

RNA was isolated from each cell by using the CellAmp^TM^ Direct RNA Prep Kit for RT-PCR (TaKaRa). First-strand cDNA was synthesized using PrimeScript^TM^ RT Master Mix (TaKaRa). The reaction was performed in a 20 μL reaction mixture. cDNAs were stored at −20°C. For qRT-PCR, the reaction mixture was prepared with SYBR Premix Ex Taq (Servicebio) according to the manufacturer’s instructions and PCR was performed with a Prism 7000 Sequence Detection System (Applied Biosystems). The sequences of primers were as follows: *GZMB*-Forward: 5’-3’CCACTCTCGACCCTACATGG; *GZMB*-Reverse: 5’-3’ GGCCCCCAAAGTGACATTTATT; *GAPDH*-Forward: 5’-3 AGGTCGGTGTGAACGGATTTG; *GAPDH*-Reverse: 5’-3’ TGTAGACCATGTAGTTGAGGTCA. PCR was performed under the following conditions: 95°C for 30 s, 40 cycles of 95°C for 5 s, and 60°C for 30 s followed by a standard dissociation run to obtain melt curve profiles of the amplicons. Using the 2 ^-ΔΔCt^ method, relative internal mRNA expression of target genes was normalized to *GAPDH*.

#### PE immunoprecipitations and LC-MS/MS analyses

With the help of The ChemBio Company, PE was synthetized. For a report on the synthesis process, as shown in Fig. 3a. CD8^+^ T cells were stimulated with IL-2 and αCD3/αCD28 for 48 hours, and then in the presence or absence of 100 μmol/L PE, after 6 hours to collect the cells. Cells were sonicated in 500 μL of PBS containing protease inhibitor cocktails (NCM Biotech) in 4℃. After centrifugation at 21,500 × g for 30 min at 4°C to remove debris. To label norepinehrinylated proteins, PE was conjugated to a biotin–azide molecule (Thermo Fisher) using the copper-click chemistry following by a freshly prepared click mixture (1 mmol/L CuSO_4_, 1 mmol/L TCEP, 100 µmol/L TBTA, 100 µmol/L axide-biotin)^41^. The mixture was incubated by rotating at room temperature for 1 hour. The resulting click-labeled lysates were centrifuged at 8000 × g for 5 min at 4°C and washed twice with 1 mL cold methanol. The proteins were resuspended in 1 mL PBS containing 0.2% SDS. Samples were quantified and 5% of the lysates were retained as inputs. 100 μL streptavidin beads (Thermo Fisher Scientific) were washed for three times with 1 mL PBS, and resuspended in 100 μL PBS, which was added to the protein solution. The resulting solution was incubated for 4 hours at 30°C, followed by washing with 1 mL PBS for three times.

For LC-MS/MS analyses, proteins were isolated in 10% SDS-PAGE gels and stained with coomassie brilliant blue. Differentially expressed protein bands were excised from SDS-PAGE gels and digested with Trypsin (Promega) for 20 hours at 37°C. The digested peptides were desalted using C18 StageTip. Peptides were taken from each sample for chromatographic separation using an Easy nLC 1200 chromatographic system (Thermo Fisher) with a nanoliter flow rate. Buffer: A solution was 0.1% formic acid in water, and B solution was 0.1% formic acid, acetonitrile and water (wherein acetonitrile was 95%). The column is equilibrated with 100% liquid A. The sample is injected into Trap Column and then passed through the chromatographic analysis column for gradient separation, with a flow rate of 300 nL/min. The liquid phase separation gradient was as follows: 0 min-2 min, linear gradient of buffer B from 2 % to 8 %; 2 min-42 min, linear gradient of buffer B from 8 % to 28 %; 42 min-50 min, linear gradient of buffer B from 28 % to 40 %; 50 min-51 min, buffer B linear gradient from 40% to 100%; 51 min-60 min, buffer B maintained at 100%. Peptide separation was followed by Data Dependent Acquisition (DDA) mass spectrometry analysis using a Q-Exactive HF-X mass spectrometer (Thermo Fisher). The MS detection mode was positive and precursor ion MS scanning range was set as 300-1500 m/z. The full MS scan was resolution 60,000 at m/z 200, AGC target 3e6, and maximum IT 50 ms. The peptide MS/MS were collected according to the 20 highest-intensity precursor ions after each full scan. The MS2 parameters were set as following: Resolution 15,000 at m/z 200, AGC target 1e5, Maximum IT 50 ms, MS2 activation type HCD, isolation window 1.6 m/z, Normalized collision energy 28. The mass spectrometry database search software used was MaxQuant 1.6.1.0. The protein database used was the Uniprot Protein Database, and the species was Mus musculus.

#### Enzymatic norepinephrinylation assays

MDC (5 mmol/L) or NE (1 mmol/L) was transaminated to GZMB peptide (10 μg) by 0.25 μg functional TGM2 protein (Novus), in a final volume of 25 μL enzymatic buffer (25 mmol/L Tris-Cl, pH 8 and 5 mmol/L CaCl_2_, plus protease inhibitors).

Peptides of GZMB were synthesized by Jiangsu JiTai Peptide Industry Science and Technology Co, Ltd. Transamidation of MDC, NE or PE to GZMB peptides (10 μg) was performed with the same procedures as described above for protein transamination, followed by LC-MS/MS analyses of norepinehrinylations.

#### Western blotting

CD8^+^ T cells or EL4 cells were activated with IL-2 and αCD3/αCD28 for 72 hours. After 72 hours cells were suspended in 500 μL of lysis buffer supplemented with protease inhibitor cocktails on ice for 30 min. Mouse spleen and lymph node tissues were homogenized at a controlled frequency of 50 Hz for a duration of 60 seconds. After centrifugation at 21,500 × g for 10 min at 4°C, 25 µg of these samples were suspended in an SDS sample buffer. After the samples were boiled, they were subjected to 12% SDS-PAGE. After SDS-PAGE, the proteins were transferred to nitrocellulose membranes (PALL). The membranes were blocked in 5% milk (Sangon Biotech) in TBST buffer (10 mmol/L Tris-HCl, pH 7.5, 150 mmol/L NaCl, and 0.05% (v/v) Tween-20) and probed with the appropriate antibodies diluted in QuickBlockTM Primary Antibody Dilution Buffer (Beyotime) for 1 hour at room temperature or overnight at 4°C. After washing with TBST three times, the membranes were incubated with HRP-conjugated secondary antibodies diluted in 5% milk in TBST buffer for 1 hour at room temperature and washed three times in TBST buffer. The following primary antibodies were used: rabbit anti-GZMB (proetintech, 1: 1000); mouse anti-GZMB (Biolegend, 1: 1000); rabbit anti-NE (LSbio, 1: 1000); rabbit anti-UHRF1 (PTMBIO, 1:1000); mouse anti-β-actin (Share-bio, 1:3000). Primary antibodies were detected using the secondary anti-mouse IgG HRP-linked (Cell signaling technology), or anti-rabbit IgG HRP-linked (Cell signaling technology, 1:1000). Signals were detected with ECL Substrate (Bio-Rad). Images were captured using a Bio-Rad system.

#### Co-immunoprecipitation

CD8^+^ T cells, EL4 cells expressing or not expressing Flag-WT-GZMB, Flag-Q43A-GZMB were lysed according to the Western Blot method previously described, and protein quantification was performed using the BCA method. For immunoprecipitation of GZMB or norepinehrinylated proteins, freshly prepared samples containing 600 µg of total protein were pre-cleaned via incubation with protein A/G magnetic beads for 40 min at 4°C. The supernatant was incubated with 2 µg of anti-GZMB or anti-NE antibody at 4°C overnight. Immune complexes were precipitated by protein A/G magnetic beads at 4°C for 2 hours. The immunoprecipitates were washed 5 × with lysis buffer on ice for three times. For immunoprecipitation of Flag-WT-GZMB or Flag-Q43A-GZMB, immunoprecipitation was carried out with Anti-Flag beads by rotating continuously at 4°C for 12 hours. The beads were then thoroughly washed with PBS containing 0.1% Tween, followed by the addition of SDS sample buffer. The proteins were dissociated from the beads by heating at 100°C for 10 min, and the supernatant was collected for WB analysis.

#### Protein stability assay

To evaluate the effect of GZMB protein stability, CD8^+^ T or EL4 cells were treated with the protein synthesis inhibitor cycloheximide (CHX). Cells were harvested at 0, 1, 2, 3 and 4 hours after CHX (10 mmol/L) treatment, and the protein was extracted. The GZMB protein levels were detected by western blot.

#### Ubiquitination level detection assay

CD8^+^ T and EL4 cells were treated with proteasome inhibitor MG-132 (10 mmol/L). After 6 hours, the protein in the cells was extracted. 20 µg protein was detected by western blot. Another 0.5 mg protein was taken out and incubated with the reference antibody overnight at 4℃ with inversion, and then incubated with 50 μL beads for 1 hours at 4℃ with inversion. Finally, the ubiquitin level of GZMB protein was detected by Co-IP using anti-Ubiquitin antibody.

#### 3D imaging of lymph nodes

##### Lymph Nodes clearing

The lymph nodes clearing were carried out as previously described^16^. Briefly, the samples were washed with 1× PBST (3 × 1 hour) to remove formaldehyde, followed by decolorization in clearing buffer at 37°C for 2 days (buffer changed daily). After PBS washes (3 × 1 hour), tissue clearing was performed through a graded THF series (50%, 70%, 80%, 95%, 30 min each) at 4°C, two treatments in 100% dichloromethane (2 × 1 hour) at 4°C, and a reverse THF gradient. Finally, samples were washed with PBS and incubated in Easy Index (37°C, overnight; then RT) for refractive index matching before imaging.

##### Sample staining

Immunostaining was performed as previously described^16^. Briefly, a 5% (w/v) blocking solution was prepared using Bovine Serum Albumin in sterile 1× PBS and used immediately. Cleared samples were blocked overnight in BSA solution at 4°C with shaking. After three 1-hour washes with 1× PBS at room temperature, samples were incubated with primary antibody against Tyrosine Hydroxylase (Abcam, 1:200) in 1× PBST at 37°C for 7 days with shaking. Following another three 1-hour PBS washes, samples were incubated with secondary antibody (ShareBio, 1:200) at 37°C for 7 days with shaking.

##### Light sheet fluorescence microscopy (LSFM) imaging

Prepare a 1-2% (w/v) solution of low-melting agarose (BBI) in Easy Index buffer. Heat the solution in a microwave until the agarose is completely dissolved, and remove any bubbles by gentle mixing or brief centrifugation. Subsequently, maintain the molten agarose at 65°C in a constant-temperature oven to prevent premature gelling. Thoroughly clean glass slides with absolute ethanol and ensure they are free of dust or residues. Pipette the molten agarose evenly onto the pre-cleaned slides, taking care to avoid bubble formation. Using fine-tipped forceps, carefully position the samples onto the agarose-coated slides. Image the LN samples in the final step using a 9 × objective (voxel size: 0.72 μm) on a MegaSPIM light-sheet microscope (LifeCanvas Technologies). Maintain consistent laser intensity settings across all samples to ensure reproducible imaging conditions.

##### Image processing and quantization

Utilize Imaris 10.1 (Oxford Instruments) to perform 3D reconstruction of the imported imaging data. For each sample channel, apply the Surface module to segment regions exhibiting positive target signals. Subsequently, employ the Filament module to connect the segmented surfaces and reconstruct continuous structures. Quantitative data, including Filament Segment Length (sum) and Filament Number of Segment Branch Points, were then exported for further statistical analysis.

#### GZMB activity assay

GZMB activity was analyzed with assay kits (abcam) following the manufacturer’s instructions. Human GZMB protein is positive control, 3 μL GZMB protein was used to do TGM2 monoaminylation assay reaction in the presence or absence of 5 μmol/L NE. The reaction buffer was incubated at 37℃ for 3 hours in the dark, then following the manufacturer’s instructions and measure immediately fluorescence at Ex/Em = 380/500 nm in a microplate reader in kinetic mode for 30 min at 37°C protected from light.

#### Quantification and statistical analysis

Statistical analysis of the data was performed using GraphPad Prism 10 software (GraphPad Software). Statistical significance was determined with a two-tailed unpaired Welcht’s *t*-test or tow-tailed unpaired *t*-test. Detailed statistical methods are described in the corresponding Figure legends. *P* < 0.05 was considered statistically significant. Each experiment was repeated with at least three independent biological replicates.

## Ethics declaration

This study involved animal subjects and was approved by the Laboratory Animal Center of Shanghai Jiao Tong University (ID: A2023083).

## Acknowledgments

This study was supported by the National Natural Science Foundation of China (82350123, 82230087, 82203228), the Shanghai Municipal Education Commission-Gaofeng Clinical Medicine Grant Support (20181708), Innovative research team of high-level local universities in Shanghai (SHSMU-ZDCX20210802), Shanghai Eastern Talent Plan (QNWS2024104), Shanghai Pilot Program for Basic Research - Shanghai Jiao Tong University (21TQ00225), 111 project (no. B21024), Shanghai Science and Technology Commission Sailing Project (ID 22YF1445600), Key Areas Research and Development Programs of Guangdong Province (ID 2023B1111050009).

## Author contributions

Z.-G. Z. and X.-L Z. conceived the project. Z.-G. Z., D.-X. L., and X.X. supervised the project. Y. Y., X.-L Z., A. T. and S.-Y. X. designed, performed, and analyzed most experiments. Y.-Z. Q., J.-M. L., G.-H. S., R. L., and J.-J. W. helped with cell culture studies. X.-Q. L., H.-T. M. helped with animal studies. W.-T. S. and Y.-X. H helped with immunoprecipitation and western blotting analyses. J.-L. H. helped with mass-spectrometry analysis. L. Q., L.-P. H., and J. J., helped with scientific writing. All authors discussed the results and revised the manuscript.

---

**Extended Data Fig. 1.**
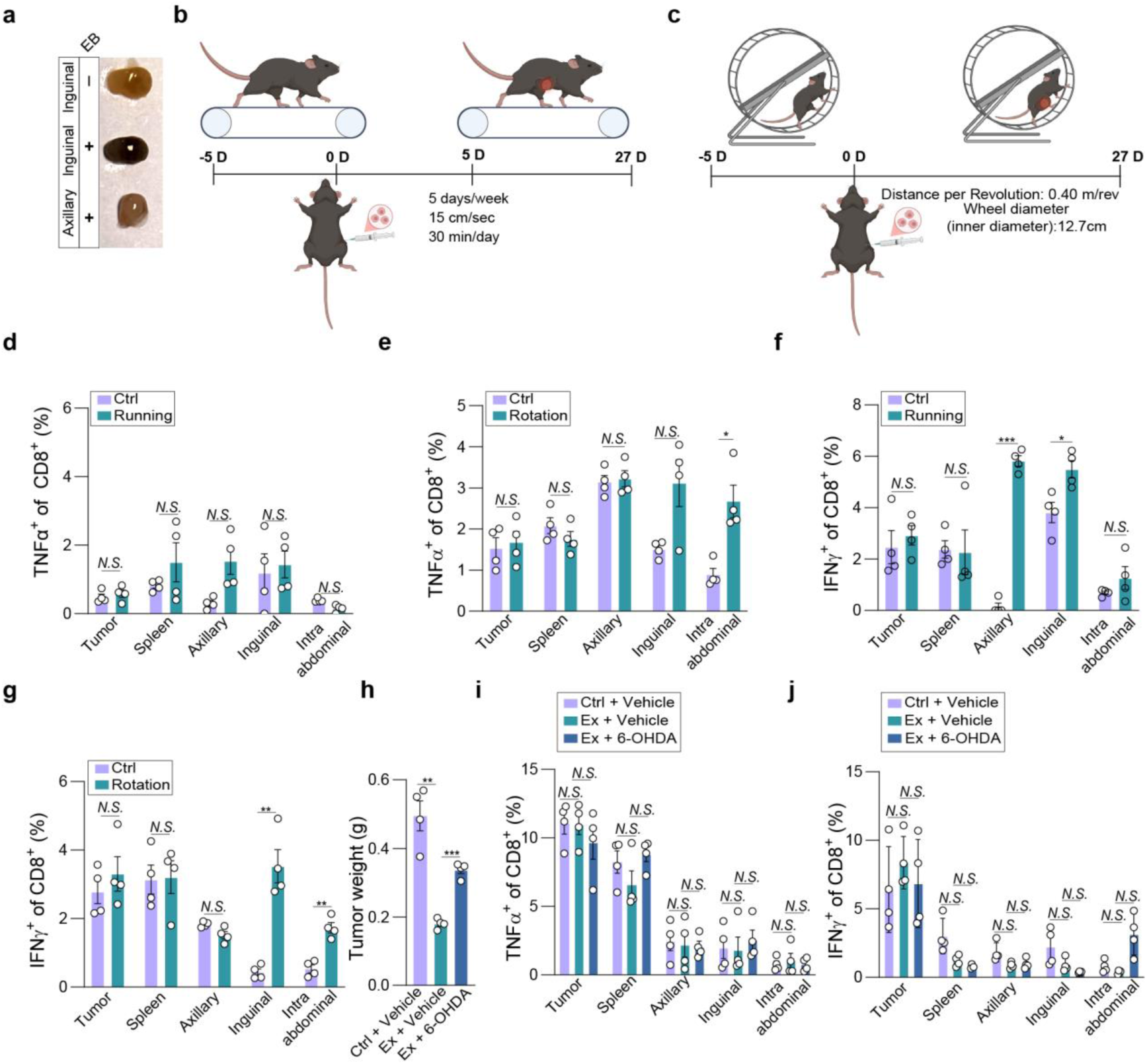
Exercise-mediated tumor suppression requires norepinephrine release from sympathetic nerves mainly in draining lymph node. **a,** Gross appearance views of the lymph nodes within (+) Evan blue (EB) or without (-) Evan blue. **b-c**, Schematic illustrations of the two exercise paradigms: The forced treadmill running model (**b**) and voluntary wheel running model (**c**). **d-e**, Frequency and intracellular TNF-α of CD8⁺ T cells were detected at 27 days after KPC1199 implantation. Running model (**d**), and voluntary wheel running model (**e**) (n = 4 per group, 3 independent experiments). **f-g,** Frequency and intracellular IFN-γ of CD8⁺ T cells were detected at 27 days after KPC1199 implantation. Runnig modle (**f**), and voluntary wheel running modle (**g**) (n = 4 per group, 3 independent experiments). **h**, KPC1199 tumor cells were subcutaneously implanted into mice. At the same time, sympathetic ablation was performed on the inguinal lymph nodes. Tumor weight was measured at 27 days after implantation. **i-j**, Frequency and intracellular TNF-α or IFN-γ of CD8⁺ T cells were detected at 27 days after KPC1199 implantation. TNF-α (**i**), and IFN-γ (**j**) (n = 4 per group, 3 independent experiments). Statistical significance was determined by two-tailed unpaired Welcht’s *t*-test in **d**, **e**, **f**, **g**, **h**, **i**, and **j**, Error bars: SEM. \**P* < 0.05, \*\**P* < 0.01, \*\*\**P* < 0.001; *N.S.*, not significant.

**Extended Data Fig. 2.**
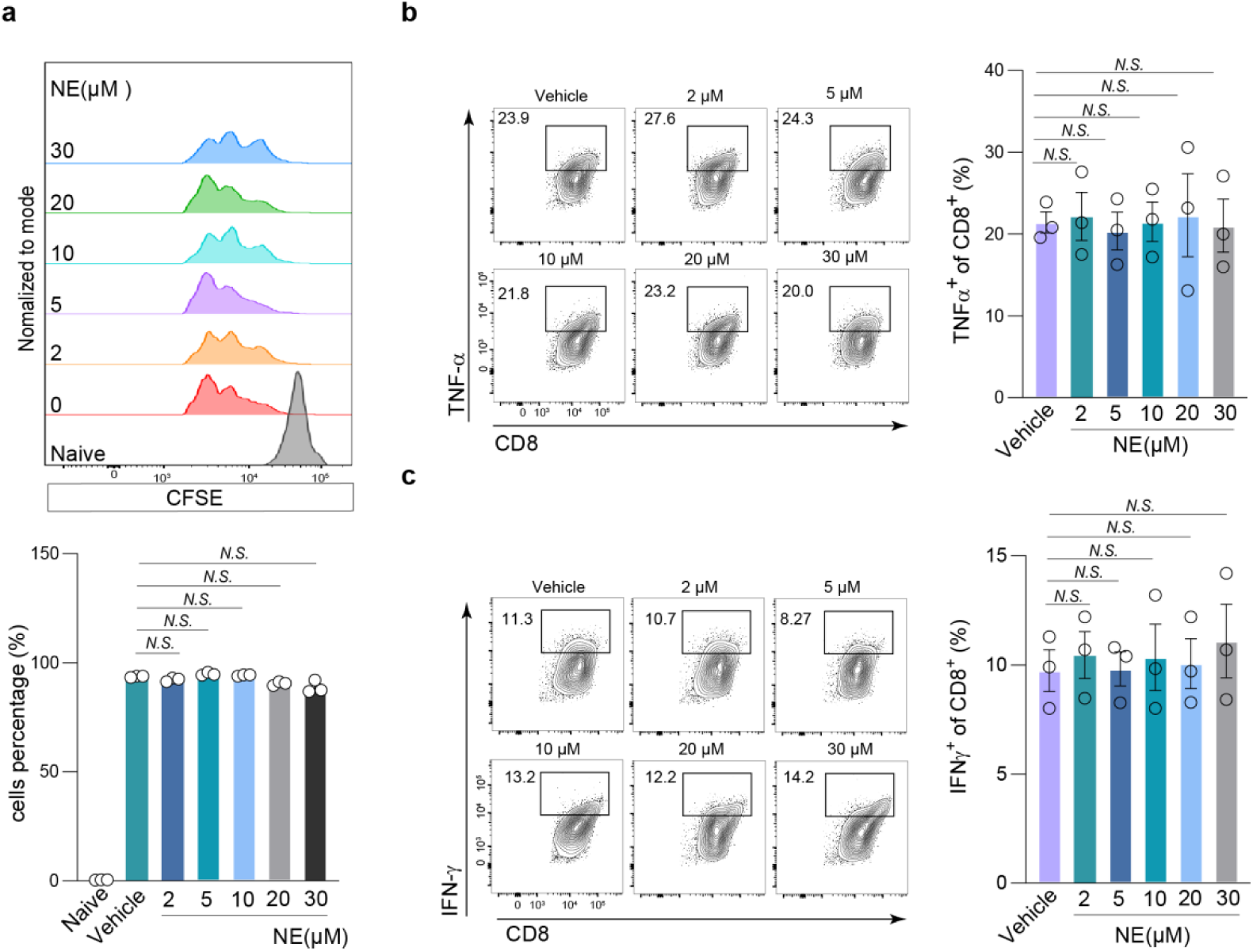
Norepinephrine selectively enhances GZMB reserves in CD8 ⁺T cells. **a**, CFSE-labelled CD8+ T cells were stimulated with IL-2 and αCD3/αCD28 for 72 hours, in the presence or absence of 0, 2, 5 or 10 μM NE. Cell proliferation were determined by flow cytometry (n = 3 per group, 3 independent experiments). **b-c,** CD8^+^ T cells were stimulated with αCD3/αCD28 for 72 hours, in the absence (Vehicle) or presence of 2, 5, 10, 20 or 30 μM norepinephrine (NE). Intracellular TNF-α protein level (**b**) and IFN-γ protein level (**c**) was determined by flow cytometry. Left panels: Representative FACS dot plot images. Right panels: Bar graphs show the quantification of protein expression (n = 3 per group, 3 independent experiments). Statistical significance was determined by two-tailed unpaired Welcht’s *t*-test in **a**, **b** and **c**, Error bars: SEM. *N.S.*, not significant.

**Extended Data Fig. 3.**
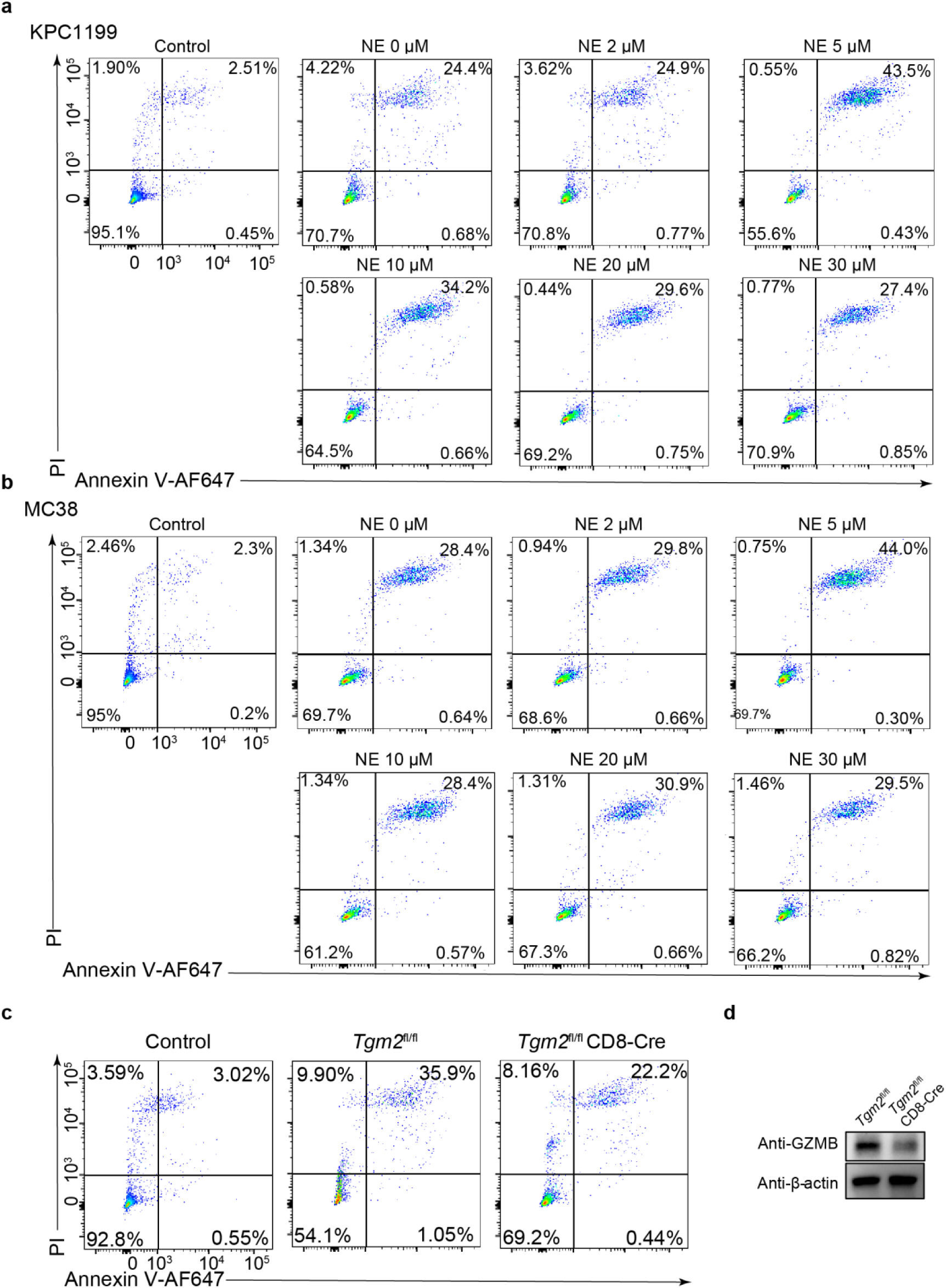
NE enhances the anti-tumor ability of CD8^+^ T cells. **a-b**, Representative FACS dot plot images of apoptosis in KPC1199 tumor cells (**a**) and MC38 tumor cells (**b**). As a control, tumor cells without incubation with CD8^+^ T were subjected to FACS analysis. **c**, Representative FACS dot plots showing the apoptosis of KPC1199 tumor cells with/without co-culturing with *Tgm2*^fl/fl^ or *Tgm2*^fl/fl^CD8-Cre CD8^+^ T cells for 24 hours. **d**, *Tgm2*^fl/fl^ or *Tgm2*^fl/fl^CD8-Cre CD8^+^ T cells were stimulated with IL-2 and αCD3/αCD28 for 72 hours. GZMB was analyzed by western blotting. β-actin was set as control (3 independent experiments).

**Extended Data Fig. 4.**
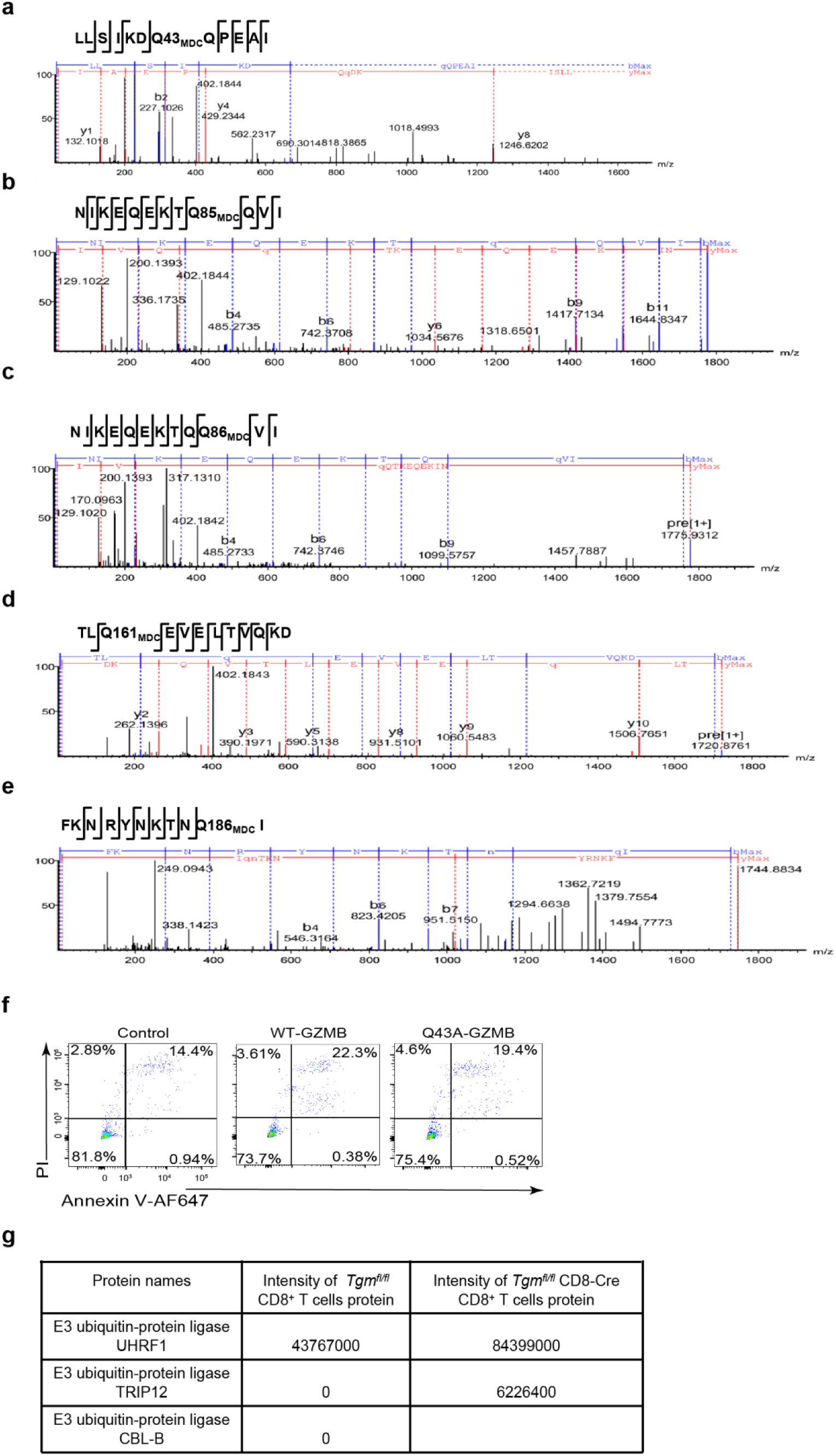
Identification of the sites for norepinephrinylation of GZMB. **a-e**, Targeted liquid chromatography–tandem mass spectrometry (LC-MS/MS) analysis of a TGM2-transamidated monodansylcadaverine (MDC) to GZMB peptide 37-48 (**a**), peptide 77-89 (**b** and **c**), peptide 159-170 (**d**), peptide 177-187 (**e**). **f**. Representative FACS dot plots showing the apoptosis of KPC1199 tumor cells with/without co-culturing with EL4 cells. **g**, Mass spectrometry analysis was conducted on E3 ligase proteins that interacted with GZMB, the samples purified using GZMB antibody in Co-IP experiments.

**Extended Data Fig. 5.**
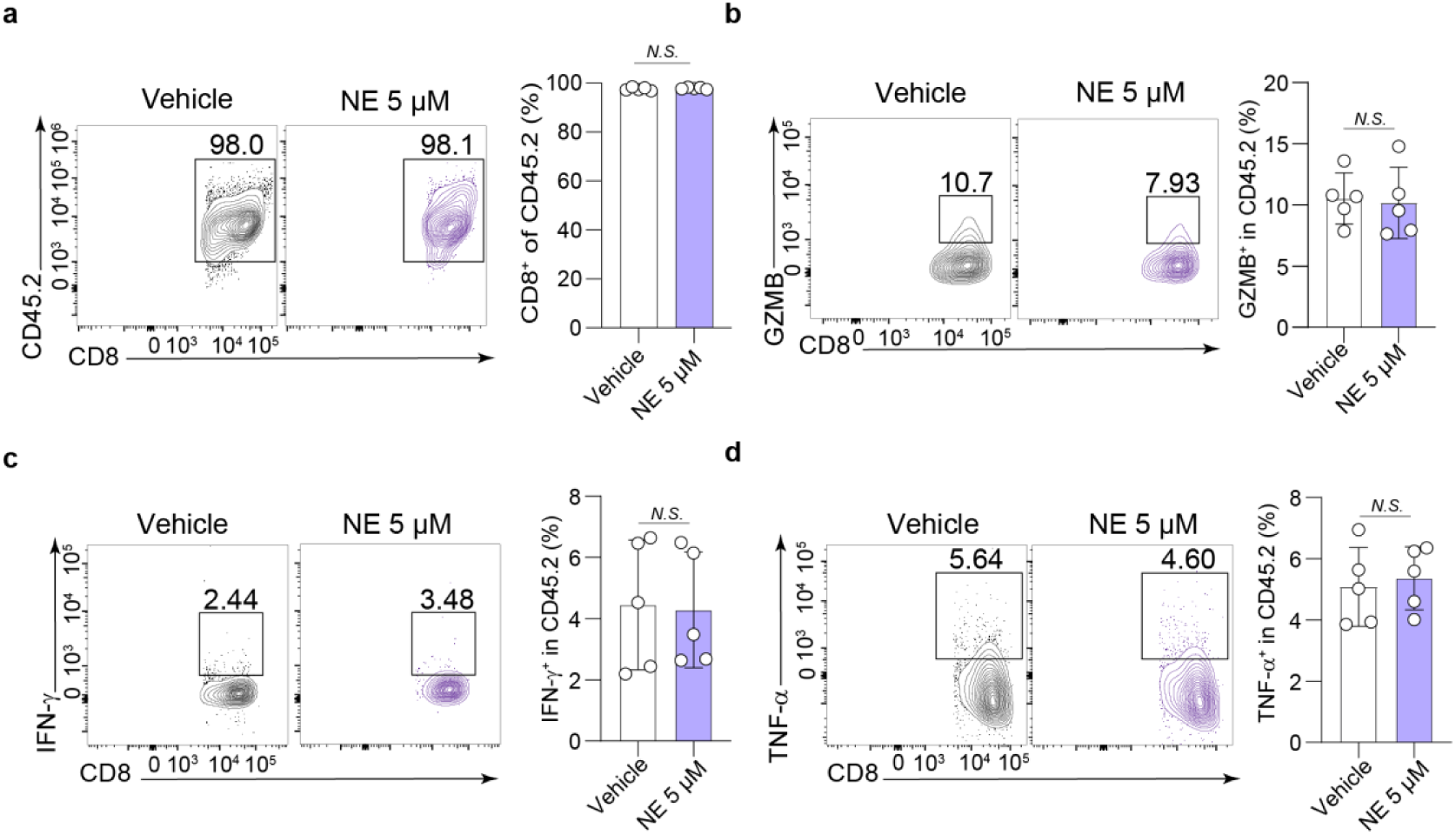
NE stimulation increases the amount of GZMB protein in CD8^+^ T cells. **a**, The frequency CD8^+^ T cells of CD45.2 and intracellular GZMB of CD8^+^ T cells (CD45.2) were detected at 30 days after implantation (n = 5 per group, 3 independent experiments). **b-c,** Intracellular GZMB of CD8^+^ T cells (CD45.2) (**b**), IFN-γ of CD8^+^ T cells (CD45.2) (**c**) and TNF-α of CD8^+^ T cells (CD45.1) (n = 5 per group, 3 independent experiments). **d,** were detected at 27 days after implantation (n = 5 per group, 3 independent experiments). GZMB was analyzed by western blotting. β-actin was set as control (3 independent experiments). Statistical significance was determined by two-tailed unpaired Welcht’s *t*-test in **a**, **b**, **c**, and **d**. Error bars: SEM. *N.S.*, not significant.

## Reference

1. Zhang, N. & Bevan, M. J. CD8+ T Cells: Foot Soldiers of the Immune System. Immunity 35, 161–168 (2011).

2. Mahanti, K. & Bhattacharyya, S. Rough neighborhood: Intricacies of cancer stem cells and infiltrating immune cell interaction in tumor microenvironment and potential in therapeutic targeting. Translational Research 265, 51–70 (2024).

3. Kaech, S. M. & Cui, W. Transcriptional control of effector and memory CD8+ T cell differentiation. Nat Rev Immunol 12, 749–761 (2012).

4. McLane, L. M., Abdel-Hakeem, M. S. & Wherry, E. J. CD8 T Cell Exhaustion During Chronic Viral Infection and Cancer. Annu. Rev. Immunol. 37, 457–495 (2019).

5. Pace, L. et al. The epigenetic control of stemness in CD8^+^ T cell fate commitment. Science 359, 177–186 (2018).

6. Philip, M. & Schietinger, A. CD8+ T cell differentiation and dysfunction in cancer. Nat Rev Immunol 22, 209–223 (2022).

7. Klein Geltink, R. I., Kyle, R. L. & Pearce, E. L. Unraveling the Complex Interplay Between T Cell Metabolism and Function. Annu. Rev. Immunol. 36, 461–488 (2018).

8. Melo-Silva, C. R. & Sigal, L. J. Innate and adaptive immune responses that control lymph-borne viruses in the draining lymph node. Cell Mol Immunol 21, 999–1007 (2024).

9. Henrickson, S. E. et al. Antigen Availability Determines CD8+ T Cell-Dendritic Cell Interaction Kinetics and Memory Fate Decisions. Immunity 39, 496–507 (2013).

10. Kastenmüller, W., Torabi-Parizi, P., Subramanian, N., Lämmermann, T. & Germain, R.N. A Spatially-Organized Multicellular Innate Immune Response in Lymph Nodes Limits Systemic Pathogen Spread. Cell 150, 1235–1248 (2012).

11. Prokhnevska, N. et al. CD8+ T cell activation in cancer comprises an initial activation phase in lymph nodes followed by effector differentiation within the tumor. Immunity 56, 107–124.e5 (2023).

12. Connolly, K. A., et al. A reservoir of stem-like CD8^+^ T cells in the tumor-draining lymph node preserves the ongoing antitumor immune response. Sci. Immunol. 6, eabg7836 (2021).

13. Lieberman, J. The ABCs of granule-mediated cytotoxicity: new weapons in the arsenal. Nat Rev Immunol 3, 361–370 (2003).

14. Martínez-Lostao, L., Anel, A. & Pardo, J. How Do Cytotoxic Lymphocytes Kill Cancer Cells? Clinical Cancer Research 21, 5047–5056 (2015).

15. Griffiths, G. M. & Isaaz, S. Granzymes A and B are targeted to the lytic granules of lymphocytes by the mannose-6-phosphate receptor. The Journal of cell biology 120, 885–896 (1993).

16. Elenkov, I. J., Wilder, R. L., Chrousos, G. P. & Vizi, E. S. The Sympathetic Nerve—An Integrative Interface between Two Supersystems: The Brain and the Immune System. Pharmacological Reviews 52, 595–638 (2000).

17. Zhang, Z.-G., et al. Fibroblastic reticular cell–derived HGF orchestrates sympathetic nerves in tumor-induced lymph node remodeling and metastasis. Preprint at 10.21203/rs.3.rs-6607085/v1 (2025).

18. Zouhal, H., Jacob, C., Delamarche, P. & Gratas-Delamarche, A. Catecholamines and the Effects of Exercise, Training and Gender: Sports Medicine 38, 401–423 (2008).

19. Moriyama, S. et al. β_2_ -adrenergic receptor–mediated negative regulation of group 2 innate lymphoid cell responses. Science 359, 1056–1061 (2018).

20. Sanders, V. M. & Straub, R. H. Norepinephrine, the β-Adrenergic Receptor, and Immunity. Brain, Behavior, and Immunity 16, 290–332 (2002).

21. Bottaro, D., Shepro, D., Peterson, S. & Hechtman, H. B. Serotonin, norepinephrine, and histamine mediation of endothelial cell barrier function in vitro. Journal Cellular Physiology 128, 189–194 (1986).

22. Walther, D. J., Stahlberg, S. & Vowinckel, J. Novel roles for biogenic monoamines: from monoamines in transglutaminase-mediated post-translational protein modification to monoaminylation deregulation diseases. The FEBS Journal 278, 4740–4755 (2011).

23. Hummerich, R., Thumfart, J.-O., Findeisen, P., Bartsch, D. & Schloss, P. Transglutaminase-mediated transamidation of serotonin, dopamine and noradrenaline to fibronectin: Evidence for a general mechanism of monoaminylation. FEBS Letters 586, 3421–3428 (2012).

24. Johnson, K. Modification of proteins by norepinephrine is important for vascular contraction. Front. Physio. 1, (2010).

25. Wan, H. et al. Regional cerebral perfusion and sympathetic activation during exercise in hypoxia and hypercapnia: preliminary insight into ‘Cushing’s mechanism’. The Journal of Physiology 603, 5103–5119 (2025).

26. Wennerberg, E. et al. Exercise reduces immune suppression and breast cancer progression in a preclinical model. Oncotarget 11, 452–461 (2020).

27. Ashcraft, K. A., Peace, R. M., Betof, A. S., Dewhirst, M. W. & Jones, L. W. Efficacy and Mechanisms of Aerobic Exercise on Cancer Initiation, Progression, and Metastasis: A Critical Systematic Review of *In Vivo* Preclinical Data. Cancer Research 76, 4032–4050 (2016).

28. Harrell, M. I., Iritani, B. M. & Ruddell, A. Lymph node mapping in the mouse. Journal of Immunological Methods 332, 170–174 (2008).

29. Malmfors, T. & Sachs, C. Degeneration of adrenergic nerves produced by 6-hydroxydopamine. European Journal of Pharmacology 3, 89–92 (1968).

30. Wang, X. et al. A GAPDH serotonylation system couples CD8+ T cell glycolytic metabolism to antitumor immunity. Molecular Cell 84, 760–775.e7 (2024).

31. Waugh, S. M., Harris, J. L., Fletterick, R. & Craik, C. S. The structure of the pro-apoptotic protease granzyme B reveals the molecular determinants of its specificity.

32. Lord, S. J., Rajotte, R. V., Korbutt, G. S. & Bleackley, R. C. Granzyme B: a natural born killer. Immunological Reviews 193, 31–38 (2003).

33. Li, J. et al. Real-Time Detection of CTL Function Reveals Distinct Patterns of Caspase Activation Mediated by Fas versus Granzyme B. The Journal of Immunology 193, 519–528 (2014).

34. Voskoboinik, I., Whisstock, J. C. & Trapani, J. A. Perforin and granzymes: function, dysfunction and human pathology. Nat Rev Immunol 15, 388–400 (2015).

35. Kägi, D. et al. Reduced Incidence and Delayed Onset of Diabetes in Perforin-deficient Nonobese Diabetic Mice. The Journal of Experimental Medicine 186, 989–997 (1997).

36. Prokhnevska, N. et al. CD8+ T cell activation in cancer comprises an initial activation phase in lymph nodes followed by effector differentiation within the tumor. Immunity 56, 107–124.e5 (2023).

37. Drokov, M. et al. High expression of granzyme B in conventional CD4+ T cells is associated with increased relapses after allogeneic stem cells transplantation in patients with hematological malignancies. Transplant Immunology 65, 101295 (2021).

38. Kagoya, Y. et al. A novel chimeric antigen receptor containing a JAK–STAT signaling domain mediates superior antitumor effects. Nat Med 24, 352–359 (2018).

39. Omar, I., Alakhras, A., Mutwali, S. & Bakhiet, M. Molecular insights into T cell development, activation and signal transduction (Review). Biomed Rep 22, 1–8 (2025).

40. Kurz, E. et al. Exercise-induced engagement of the IL-15/IL-15Rα axis promotes anti-tumor immunity in pancreatic cancer. Cancer Cell 40, 720–737.e5 (2022).

41. Qin, W. et al. Chemoproteomic Profiling of Itaconation by Bioorthogonal Probes in Inflammatory Macrophages. J. Am. Chem. Soc. 142, 10894–10898 (2020).

