## Supplemental Table for "A Neural Arming Niche in Tumor-Draining Lymph Nodes Programs CD8⁺T Cell Cytotoxicity via GZMB Norepinephrinylation"

**Table S1** Analysis of Mass Spectrometry Data.

| Protein IDs | Gene names | Peptides | Sequence coverage [%] | Mol. weight [kDa] | Score | Intensity |
|---|---|---|---|---|---|---|
| P60710 | Actb | 20 | 52.3 | 41.736 | 323.31 | 83838000000 |
| P11499 | Hsp90ab1 | 37 | 39.5 | 83.28 | 323.31 | 27075000000 |
| Q5U405 | Tmprss13 | 1 | 1.5 | 59.805 | 6.6346 | 23547000000 |
| P10126 | Eef1a1 | 11 | 23.6 | 50.113 | 79.835 | 20240000000 |
| P58252 | Eef2 | 29 | 31 | 95.313 | 258.51 | 18038000000 |
| Q8VDD5 | Myh9 | 49 | 23.7 | 226.37 | 323.31 | 13200000000 |
| Q61003 | Cd6 | 2 | 3 | 72.255 | 11.41 | 11786000000 |
| P68372 | Tubb4b;Tubb4a | 12 | 27 | 49.83 | 107.02 | 11111000000 |
| P16858 | Gapdh;Gapdhrt; | 10 | 33 | 35.81 | 66.465 | 10724000000 |
| P63017 | Hspa8 | 26 | 40.1 | 70.87 | 257.74 | 9787200000 |
| P52480 | Pkm | 19 | 36.9 | 57.844 | 145.88 | 9531600000 |
| P06151 | Ldha | 13 | 33.1 | 36.498 | 156.39 | 8024100000 |
| P05064 | Aldoa;Aldoart1 | 16 | 43.1 | 39.355 | 140.5 | 7897400000 |
| P60843 | Eif4a1 | 17 | 36.9 | 46.153 | 233.7 | 7177600000 |
| P68373 | Tuba1c;Tuba1a; | 11 | 25.2 | 49.909 | 107.88 | 6907900000 |
| P17182 | Eno1 | 13 | 27.9 | 47.14 | 115.8 | 6671800000 |
| Q7SIG6 | Asap2 | 1 | 0.8 | 106.8 | 7.5921 | 6607600000 |
| P09405 | Ncl | 18 | 23.5 | 76.722 | 152.64 | 6053300000 |
| Q4VBD9 | Gzf1 | 1 | 1.4 | 79.532 | 5.9475 | 4540700000 |

| | | | | | | |
|---|---|---|---|---|---|---|
| P63101 | Ywhaz | 10 | 37.1 | 27.771 | 97.211 | 4109100000 |
| P42932 | Cct8 | 21 | 38.1 | 59.555 | 151.48 | 3994000000 |
| P09411 | Pgk1 | 14 | 35 | 44.55 | 105.76 | 3851000000 |
| Q9CZN7 | Shmt2 | 13 | 25.8 | 55.758 | 106.65 | 3714000000 |
| Q61656 | Ddx5 | 16 | 21.3 | 69.289 | 129 | 3680500000 |
| Q01853 | Vcp | 22 | 28.8 | 89.321 | 150.79 | 3667900000 |
| P47911 | Rpl6 | 10 | 25.3 | 33.509 | 82.509 | 3638100000 |
| Q9D8E6 | Rpl4 | 12 | 26.3 | 47.153 | 85.538 | 3516700000 |
| O89053 | Coro1a | 12 | 22.8 | 50.989 | 147.98 | 3508300000 |
| Q61753 | Phgdh | 13 | 22 | 56.585 | 101.25 | 3477200000 |
| P27659 | Rpl3 | 13 | 29.8 | 46.109 | 124.3 | 3400300000 |
| P40142 | Tkt | 16 | 28.7 | 67.63 | 151.58 | 3389100000 |
| Q8BG05 | Hnrnpa3 | 12 | 28.2 | 39.652 | 101.78 | 3365300000 |
| O88569 | Hnrnpa2b1 | 11 | 40.5 | 37.402 | 117.46 | 3241500000 |
| Q61937 | Npm1 | 5 | 16.8 | 32.56 | 42.276 | 3192100000 |
| P50580 | Pa2g4 | 12 | 34.5 | 43.698 | 82.117 | 3189300000 |
| P62918 | Rpl8 | 4 | 16.7 | 28.024 | 50.027 | 3139900000 |
| P35980 | Rpl18 | 9 | 42 | 21.644 | 84.142 | 3136400000 |
| P62754 | Rps6 | 7 | 26.9 | 28.68 | 52.908 | 3054800000 |
| Q7TPV4 | Mybbp1a | 17 | 16.1 | 152.04 | 251.28 | 2956100000 |
| P07901 | Hsp90aa1 | 24 | 28.2 | 84.787 | 103.91 | 2901800000 |
| P14148 | Rpl7 | 14 | 34.4 | 31.419 | 109.84 | 2889600000 |
| P63038 | Hspd1 | 15 | 32.3 | 60.955 | 117.37 | 2833700000 |
| P38647 | Hspa9 | 19 | 33.1 | 73.46 | 174.98 | 2813000000 |
| P49312 | Hnrnpa1 | 8 | 21.6 | 34.196 | 55.706 | 2783000000 |
| Q8R081 | Hnrnpl | 11 | 21.2 | 63.963 | 129.48 | 2774200000 |

| | | | | | | |
|---|---|---|---|---|---|---|
| P80318 | Cct3 | 23 | 40.6 | 60.629 | 157.73 | 2744100000 |
| P62908 | Rps3 | 9 | 40.7 | 26.674 | 85.361 | 2710300000 |
| Q9D8N0 | Eef1g | 11 | 27.2 | 50.06 | 86.666 | 2696800000 |
| P97351 | Rps3a | 9 | 28.4 | 29.885 | 62.096 | 2543100000 |
| Q03265 | Atp5f1a | 17 | 30.6 | 59.752 | 125.26 | 2410000000 |
| P62264 | Rps14 | 5 | 31.1 | 16.273 | 55.32 | 2399300000 |
| P47962 | Rpl5 | 10 | 26.3 | 34.4 | 78.265 | 2379000000 |
| Q80TE0 | Rpap1 | 1 | 0.7 | 155.27 | 6.2764 | 2375400000 |
| Q6ZWN5 | Rps9 | 7 | 26.3 | 22.591 | 44.557 | 2352800000 |
| P62259 | Ywhae | 12 | 45.1 | 29.174 | 99.192 | 2319900000 |
| P11983 | Tcp1 | 13 | 26.6 | 60.448 | 82.785 | 2313500000 |
| Q9CY58 | Serbp1 | 13 | 36.4 | 44.714 | 199.2 | 2305000000 |
| P62962 | Pfn1 | 8 | 60.7 | 14.957 | 67.94 | 2304200000 |
| P18760 | Cfl1 | 5 | 29.5 | 18.559 | 38.645 | 2273000000 |
| P62827 | Ran;1700009N14Rik; | 7 | 24.5 | 24.423 | 45.195 | 2256100000 |
| Q61233 | Lcp1 | 20 | 33.7 | 70.148 | 184.09 | 2209400000 |
| P29341 | Pabpc1;Pabpc6 | 14 | 21.2 | 70.67 | 106.4 | 2206300000 |
| P80315 | Cct4 | 14 | 26.7 | 58.066 | 85.452 | 2194300000 |
| Q9JIK5 | Ddx21 | 18 | 24.6 | 93.55 | 193.72 | 2177500000 |
| Q02053 | Uba1 | 16 | 18.1 | 117.81 | 147.78 | 2167300000 |
| Q61316 | Hspa4 | 17 | 26.3 | 94.132 | 175.93 | 2167000000 |
| A0A2R8VHP3 | Gm5478 | 10 | 13.6 | 57.919 | 13.155 | 2132500000 |
| P27773 | Pdia3 | 14 | 30.7 | 56.678 | 93.218 | 2118700000 |
| P19253 | Rpl13a | 6 | 24.6 | 23.464 | 36.835 | 2081000000 |

| | | | | | | |
|---|---|---|---|---|---|---|
| Q8CGP1 | H2bc12;H2bc15; | 7 | 54 | 13.92 | 79.236 | 2053000000 |
| P80317 | Cct6a | 11 | 17.5 | 58.004 | 68.268 | 2052500000 |
| E9Q557 | Dsp | 37 | 11.8 | 332.91 | 299.73 | 2044400000 |
| P12970 | Rpl7a | 11 | 35.7 | 29.976 | 72.925 | 2037400000 |
| P17742 | Ppia | 4 | 22 | 17.971 | 30.838 | 2032000000 |
| P17751 | Tpi1 | 5 | 26.1 | 26.712 | 53.674 | 2024800000 |
| P80314 | Cct2 | 12 | 22.8 | 57.477 | 108.42 | 1979600000 |
| Q01768 | Nme2 | 5 | 34.9 | 17.363 | 54.574 | 1926500000 |
| P25444 | Rps2 | 6 | 20.8 | 31.231 | 41.537 | 1895600000 |
| P62806 | H4c16 | 7 | 52.4 | 11.367 | 52.981 | 1894000000 |
| P57780 | Actn4 | 20 | 26.9 | 104.98 | 161.31 | 1852300000 |
| P62702 | Rps4x | 8 | 24.7 | 29.597 | 56.15 | 1830000000 |
| P04187 | Gzmb | 6 | 25.1 | 27.47 | 38.309 | 1805440000 |
| Q922D8 | Mthfd1 | 17 | 18.3 | 101.2 | 188.86 | 1768200000 |
| Q6ZWV3 | Rpl10;Rpl10l | 5 | 18.7 | 24.604 | 46.044 | 1747000000 |
| P47963 | Rpl13 | 8 | 35.1 | 24.305 | 90.558 | 1718300000 |
| P80313 | Cct7 | 12 | 22.8 | 59.652 | 84.123 | 1685200000 |
| P19096 | Fasn | 29 | 13.5 | 272.43 | 203.1 | 1670400000 |
| Q8BTM8 | Flna | 26 | 12.9 | 281.22 | 182.38 | 1668900000 |
| P61979 | Hnrnpk | 9 | 16.4 | 50.976 | 78.295 | 1645000000 |
| P08113 | Hsp90b1 | 13 | 20.3 | 92.475 | 102.11 | 1629100000 |
| P26041 | Msn | 14 | 18.4 | 67.766 | 103.76 | 1624300000 |
| Q9WTL4 | Insrr | 2 | 0.6 | 144.87 | 11.791 | 1596600000 |
| P15532 | Nme1 | 5 | 32.2 | 17.208 | 11.643 | 1580000000 |
| Q99K48 | Nono | 9 | 18.4 | 54.54 | 70.242 | 1526000000 |

| | | | | | | |
|---|---|---|---|---|---|---|
| Q91VC3 | Eif4a3;Eif4a3l2; | 14 | 29.2 | 46.839 | 120.63 | 1517600000 |
| P17918 | Pcna | 9 | 39.5 | 28.785 | 94.063 | 1513500000 |
| P80316 | Cct5 | 11 | 15.2 | 59.623 | 69.732 | 1486700000 |
| P99024 | Tubb5;Tubb3 | 11 | 23.2 | 49.67 | 18.126 | 1469000000 |
| Q99020 | Hnrnpab | 4 | 16.8 | 30.831 | 46.356 | 1458600000 |

**Table S2** Analysis of Mass Spectrometry Data.

| Protein IDs | Gene names | Peptides | Sequence coverage [%] | Mol. weight [kDa] | Score | Intensity |
|---|---|---|---|---|---|---|
| E9PXX4 | Pde8b | 1 | 0.8 | 82.639 | 2.8634 | 1005300000 |
| Q91YZ8 | Pabpc4 | 14 | 29.4 | 67.852 | 24.607 | 284580000 |
| P47911 | Rpl6 | 5 | 15.2 | 33.509 | 62.588 | 251350000 |
| D3YXL8 | Rab3il1 | 2 | 16.3 | 10.174 | 1.1762 | 204690000 |
| Q9ESX5 | Dkc1 | 6 | 19.1 | 57.401 | 155.49 | 204200000 |
| E9QN08 | Eef1d | 6 | 32.7 | 27.217 | 68.034 | 185290000 |
| P30681 | Hmgb2 | 6 | 20.5 | 24.162 | 11.549 | 179440000 |
| P17225 | Ptbp1 | 5 | 12.3 | 59.321 | 81.615 | 146010000 |
| Q8K0E8 | Fgb | 3 | 5.4 | 54.752 | 2.9057 | 141610000 |
| P04187 | Gzmb | 4 | 17 | 27.47 | 5.4843 | 141300000 |
| I7HLV2 | Rpl10 | 3 | 17.9 | 23.072 | 5.8823 | 140450000 |
| G3UZP7 | H2-D1 | 6 | 20.3 | 42.78 | 8.6511 | 140060000 |
| E9PW20 | Chtop | 4 | 25.6 | 22.106 | 8.6006 | 138020000 |
| P10852 | Slc3a2 | 4 | 9.9 | 58.336 | 23.901 | 130340000 |
| A0A0U1RQA1 | Septin1 | 1 | 3.4 | 37.372 | -2 | 127200000 |
| Q61599 | Arhgdib | 4 | 38.5 | 22.851 | 61.827 | 123290000 |
| Q9CR57 | Rpl14 | 3 | 17.1 | 23.564 | 27.268 | 122910000 |
| P26041 | Msn | 9 | 17.3 | 67.766 | 11.817 | 115930000 |
| Q99K94 | Stat1 | 5 | 7.6 | 83.106 | 10.544 | 111750000 |
| Q9DCH4 | Eif3f | 5 | 16.6 | 37.984 | 6.545 | 110700000 |
| P08113 | Hsp90b1 | 7 | 10.3 | 92.475 | 10.26 | 107850000 |
| P80315 | Cct4 | 8 | 21.5 | 58.066 | 31.412 | 107700000 |
| B8JK33 | Hnrnpm | 9 | 20.2 | 68.161 | 25.254 | 105320000 |
| Q61753 | Phgdh | 4 | 8.4 | 56.585 | 11.699 | 105120000 |

| | | | | | | |
|---|---|---|---|---|---|---|
| Q8VDW0 | Ddx39a | 3 | 8.4 | 49.067 | 17.174 | 103190000 |
| P17918 | Pcna | 5 | 23 | 28.785 | 8.1434 | 92084000 |
| E9Q133 | Cct3 | 10 | 20.7 | 56.526 | 21.914 | 87464000 |
| P25444 | Rps2 | 3 | 13 | 31.231 | 6.9923 | 87354000 |
| P31362 | Pou2f3 | 1 | 2.6 | 46.959 | -2 | 81399000 |
| A0A0J9YUZ4 | Hmgb1 | 5 | 19.9 | 24.233 | 5.2489 | 79211000 |
| Q8BH05 | Znf750 | 1 | 1 | 76.645 | -2 | 78968000 |
| Q99PT1 | Arhgdia | 4 | 23 | 23.407 | 39.384 | 74192000 |
| B1AU76 | Nasp | 4 | 8.9 | 48.765 | 3.9386 | 74010000 |
| Q80WS3 | Fbll1 | 4 | 16.2 | 33.338 | 1.4206 | 73552000 |
| P47962 | Rpl5 | 3 | 12.8 | 34.4 | 4.6354 | 71106000 |
| Q60932 | Vdac1 | 3 | 13.9 | 32.351 | 4.085 | 68978000 |
| P31041 | Cd28 | 2 | 10.1 | 25.243 | 2.9656 | 68317000 |
| Q52KG9 | Cct6a | 4 | 7.2 | 58.076 | 4.0716 | 68045000 |
| O70456 | Sfn | 2 | 7.3 | 27.706 | -2 | 63693000 |
| Q99KP6 | Prpf19 | 2 | 4.8 | 55.238 | 1.7241 | 60491000 |
| P70375 | F7 | 1 | 2.7 | 50.276 | -2 | 60237000 |
| Q3U2G2 | Hspa4 | 3 | 4.9 | 94.208 | 9.6384 | 59154000 |
| P62082 | Rps7 | 2 | 21.6 | 22.127 | 8.0892 | 58831000 |
| E9PYJ7 | Pitpnm2 | 1 | 1.2 | 147.62 | 0.96612 | 57410000 |
| P20801 | Tnnc2 | 2 | 18.1 | 18.11 | 1.8409 | 57253000 |
| Q5SQG5 | Phb;Phb1 | 3 | 15.5 | 22.999 | 10.729 | 55893000 |
| Q3TXX3 | Zfyve27 | 1 | 1.4 | 46.202 | 5.0837 | 55017000 |
| E0CZA1 | Cct5 | 3 | 16.6 | 21.525 | 15.517 | 54686000 |
| P35564 | Canx | 3 | 6.3 | 67.277 | 7.5531 | 53472000 |
| D3Z3N4 | Hnrnph3 | 4 | 16.2 | 36.869 | 57.14 | 52414000 |

| | | | | | | |
|---|---|---|---|---|---|---|
| P80313 | Cct7 | 5 | 11 | 59.652 | 4.2534 | 52353000 |
| P97855 | G3bp1 | 3 | 8.4 | 51.828 | 4.6958 | 50100000 |
| Q9Z1Q5 | Clic1 | 3 | 17.4 | 27.013 | 14.57 | 48627000 |
| F6UFG6 | Anp32a | 2 | 11.6 | 15.306 | 2.6131 | 48351000 |
| Q8JZQ9 | Eif3b | 4 | 5.5 | 91.369 | 28.249 | 47655000 |
| P60122 | Ruvbl1 | 3 | 8.6 | 50.213 | 5.0979 | 45466000 |
| Q9Z0E6 | Gbp2 | 2 | 4.8 | 66.739 | 34.763 | 45120000 |
| Q9EPU0 | Upf1 | 5 | 5.6 | 123.97 | 6.8553 | 43348000 |
| A0A1B0GS70 | Psma1 | 1 | 4.6 | 26.496 | 1.1979 | 42500000 |
| P30416 | Fkbp4 | 3 | 8.3 | 51.572 | 7.0266 | 42232000 |
| Q8QZY6 | Tspan14 | 1 | 3 | 30.673 | 1.2053 | 40655000 |
| Q9Z0N1 | Eif2s3x;Eif2s3y | 2 | 8.3 | 51.065 | 1.0382 | 39872000 |
| P24547 | Impdh2 | 3 | 7.6 | 55.814 | 3.2592 | 38936000 |
| P14211 | Calr | 1 | 2.2 | 47.994 | 1.8146 | 37109000 |
| A0A0A0MQ97 | Nccrp1 | 2 | 7.6 | 33.027 | 9.9539 | 37023000 |
| Q99K85 | Psat1 | 3 | 9.2 | 40.472 | 3.5829 | 36904000 |
| E9QA05 | Sp100 | 4 | 7.7 | 56.101 | 6.5366 | 36536000 |
| A0A0A0MQ76 | Nop58 | 4 | 12.4 | 50.063 | 9.2582 | 36450000 |
| A0A1L1SST0 | Ppia | 1 | 5.8 | 17.047 | 0.96336 | 36370000 |
| Q8VC69 | Slc22a6 | 1 | 2.2 | 60.013 | -2 | 36033000 |
| P62192 | Psmc1 | 2 | 7.5 | 49.184 | 29.342 | 35601000 |
| Q3TH01 | H2-K1 | 4 | 12.2 | 40.285 | 9.6066 | 33843000 |
| Q80X68 | Csl;Cs | 2 | 5.8 | 52.325 | 2.9846 | 33503000 |
| P61161 | Actr2 | 2 | 6.9 | 44.76 | 2.5438 | 33103000 |
| D3Z0Y2 | Prdx6 | 1 | 5.5 | 22.494 | 2.8804 | 32701000 |
| Q3TU25 | Ddx17 | 5 | 13.4 | 47.385 | 5.3385 | 32438000 |

| | | | | | | |
|---|---|---|---|---|---|---|
| Q9Z1Q9 | Vars1;Vars | 3 | 3.4 | 140.21 | 6.7613 | 32108000 |
| P70372 | Elavl1 | 3 | 10.7 | 36.169 | 7.0481 | 32092000 |
| S4R1M2 | Safb | 2 | 1.4 | 105.32 | 1.0697 | 31541000 |
| P25206 | Mcm3 | 2 | 3.2 | 91.545 | 9.4729 | 30870000 |
| Q62318 | Trim28 | 2 | 2.4 | 88.846 | 2.459 | 30458000 |
| Q921F24 | Tardbp | 2 | 7 | 44.547 | 26.841 | 30355000 |
| Q9D8S5 | Srsf5 | 2 | 9.6 | 30.978 | 3.5282 | 30083000 |
| Q3KQM4 | U2af2 | 2 | 9.4 | 33.901 | 1.5952 | 28905000 |
| P70168 | Kpnb1 | 2 | 2.6 | 97.183 | 3.7771 | 28807000 |
| P40124 | Cap1 | 2 | 6.3 | 51.564 | 3.3097 | 28503000 |
| Q6PDM2 | Srsf1 | 3 | 13.7 | 27.744 | 6.6082 | 28271000 |
| Q8C318 | Uhrf1bp1l;Bltp3b | 1 | 2.5 | 66.798 | -2 | 27496000 |
| Q9DCL9 | Paics | 2 | 4.9 | 47.006 | 2.3078 | 27284000 |
| P50247 | Ahcy | 4 | 10.6 | 47.688 | 10.581 | 27009000 |
| P16627 | Hspa1l | 7 | 11.9 | 70.636 | 2.9213 | 26667000 |
| Q8BFR5 | Tufm | 2 | 5.8 | 49.508 | 1.241 | 26589000 |
| Q8R2P8 | Kars;Kars1 | 2 | 4.2 | 71.29 | 8.8251 | 26533000 |
| H3BJ30 | Cpsf6 | 2 | 5.1 | 59.308 | 5.6435 | 25746000 |
| D3YZ06 | Hspb1 | 2 | 12.9 | 16.818 | 1.1842 | 25586000 |
| Q3TML0 | Pdia6 | 2 | 6.3 | 48.689 | 4.005 | 25561000 |
| K3W4S6 | Gyg;Gyg1 | 1 | 2.7 | 41.926 | 1.1756 | 25392000 |
| Q8C879 | Zfp202 | 1 | 1.6 | 73.38 | -2 | 25202000 |
| Q9CZN7 | Shmt2 | 3 | 8.1 | 55.758 | 7.7005 | 25174000 |

**Table S3** Analysis of Mass Spectrometry Data.

| Protein IDs | Gene names | Mol. weight [kDa] | Score | Peptides *Tgm2*$^{fl/fl}$ CD8 cells protein | Peptides *Tgm2*$^{fl/fl}$ CD8-Cre CD8 cells protein | iBAQ *Tgm2*$^{fl/fl}$ CD8 cells protein | iBAQ *Tgm2*$^{fl/fl}$ CD8-CreCD8 cells protein |
|---|---|---|---|---|---|---|---|
| P04187 | Gzmb | 27.47 | 63.826 | 10 | 13 | 305490000 | 343330000 |
| Q9JKR6 | Hyou1 | 111.18 | 63.572 | 8 | 9 | 2027900 | 3305300 |
| P42225 | Stat1 | 87.196 | 62.342 | 17 | 17 | 10531000 | 9966000 |
| P06800 | Ptprc | 144.84 | 60.549 | 9 | 11 | 7156100 | 7137700 |
| Q91WK2 | Eif3h | 39.832 | 58.452 | 6 | 7 | 2460600 | 3233000 |
| P63325 | Rps10 | 18.916 | 56.788 | 6 | 6 | 18198000 | 28654000 |
| Q9D0E1 | Hnrnpm | 77.648 | 55.459 | 15 | 17 | 10013000 | 12457000 |
| Q99JR1 | Sfxn1 | 35.649 | 54.52 | 3 | 3 | 1490200 | 1052800 |
| P01837 | Igkc | 11.934 | 54.3 | 5 | 7 | 199040000 | 320630000 |
| Q99K48 | Nono | 54.54 | 53.517 | 12 | 11 | 11483000 | 7775300 |
| P01869 | Ighg1 | 43.386 | 53.192 | 9 | 9 | 652200000 | 792130000 |
| A0A0A6YW6 7 | Gm8797;Uba52; | 8.7279 | 52.857 | 3 | 5 | 101630000 | 284240000 |
| Q9DCE9 | Igtp | 48.493 | 52.433 | 6 | 4 | 2823700 | 3750500 |
| B2RQC6 | Cad | 243.24 | 51.795 | 13 | 18 | 915460 | 1474900 |
| P51150 | Rab7a | 23.489 | 50.127 | 4 | 6 | 5231300 | 15711000 |
| Q8VEK3 | Hnrnpu | 87.917 | 50.014 | 14 | 16 | 28141000 | 44972000 |
| P97351 | Rps3a | 29.885 | 47.712 | 13 | 14 | 40743000 | 82785000 |
| Q61937 | Npm1 | 32.56 | 47.684 | 9 | 12 | 44866000 | 97448000 |
| P80316 | Cct5 | 59.623 | 47.466 | 10 | 13 | 8992200 | 16058000 |
| Q7TPR4 | Actn1 | 103.07 | 46.576 | 8 | 9 | 725400 | 739870 |
| Q9Z0N1 | Eif2s3x;Eif2s3y | 51.065 | 46.335 | 7 | 9 | 5309600 | 14676000 |

| | | | | | | | |
|---|---|---|---|---|---|---|---|
| P19096 | Fasn | 272.43 | 46.194 | 14 | 17 | 1774100 | 2522100 |
| P62192 | Psmc1 | 49.184 | 45.994 | 6 | 9 | 2224200 | 5594800 |
| Q99JY3 | Gimap4 | 38.044 | 45.895 | 7 | 5 | 5582700 | 2258400 |
| P15702 | Spn | 40.038 | 44.855 | 1 | 1 | 1911500 | 0 |
| Q6ZWX6 | Eif2s1 | 36.108 | 44.083 | 6 | 7 | 9363100 | 14907000 |
| P57776 | Eef1d | 31.293 | 42.146 | 4 | 5 | 2149500 | 6870100 |
| P56959 | Fus | 52.673 | 41.905 | 10 | 9 | 22769000 | 15452000 |
| P11983 | Tcp1 | 60.448 | 41.74 | 10 | 13 | 11178000 | 17516000 |
| Q9D1G1 | Rab1b | 22.187 | 41.728 | 5 | 5 | 12521000 | 13860000 |
| P97461 | Rps5 | 22.876 | 41.72 | 3 | 5 | 10312000 | 18962000 |
| Q9JIK5 | Ddx21 | 93.55 | 41.581 | 14 | 16 | 8067300 | 16014000 |
| P62242 | Rps8 | 24.205 | 41.385 | 8 | 9 | 78639000 | 115540000 |
| O70133 | Dhx9 | 149.47 | 41.373 | 19 | 19 | 6314000 | 8104400 |
| P54823 | Ddx6 | 54.191 | 41.175 | 6 | 6 | 2117600 | 3966600 |
| Q9QZQ8 | Macroh2a1 | 39.289 | 40.517 | 3 | 4 | 3055700 | 7742300 |
| Q9CS42 | Prps2;Prps1l3; | 34.786 | 40.5 | 2 | 4 | 336220 | 2677200 |
| Q9D6Z1 | Nop56 | 64.464 | 40.186 | 12 | 14 | 5192000 | 11465000 |
| P62900 | Rpl31 | 14.463 | 39.797 | 4 | 5 | 30940000 | 56349000 |
| Q1HFZ0 | Nsun2 | 85.451 | 39.739 | 8 | 10 | 1941200 | 4713000 |
| Q61233 | Lcp1 | 70.148 | 39.119 | 17 | 17 | 14811000 | 20867000 |
| P63276 | Rps17 | 15.524 | 38.74 | 6 | 5 | 55128000 | 70715000 |
| P97350 | Pkp1 | 80.895 | 38.635 | 3 | 6 | 1084300 | 3913800 |
| P14869 | Rplp0 | 34.216 | 38.499 | 9 | 10 | 26556000 | 49130000 |
| Q61635 | Ifi47 | 46.784 | 38.456 | 11 | 12 | 7697200 | 10083000 |
| Q9Z130 | Hnrnpdl | 33.558 | 38.297 | 7 | 8 | 14355000 | 16903000 |

| | | | | | | | |
|---|---|---|---|---|---|---|---|
| P48024 | Eif1;Eif1b | 12.746 | 38.014 | 2 | 2 | 5083900 | 16564000 |
| Q62318 | Trim28 | 88.846 | 37.951 | 7 | 10 | 2622800 | 4394000 |
| P47963 | Rpl13 | 24.305 | 37.732 | 4 | 8 | 25344000 | 50645000 |
| P80318 | Cct3 | 60.629 | 37.192 | 22 | 21 | 20208000 | 26245000 |
| Q9Z0E6 | Gbp2 | 66.739 | 36.714 | 5 | 5 | 2772900 | 1353900 |
| Q6P4T2 | Snrnp200 | 244.54 | 36.452 | 8 | 14 | 485500 | 965400 |
| Q8BGB5 | Limd2 | 14.237 | 36.257 | 1 | 1 | 5475000 | 7037900 |
| Q8QZT1 | Acat1 | 44.816 | 35.864 | 4 | 4 | 2720600 | 4717700 |
| Q9WVA4 | Tagln2 | 22.395 | 35.395 | 9 | 8 | 16092000 | 24624000 |
| Q9Z2X1 | Hnrnpf | 45.729 | 34.946 | 9 | 7 | 24560000 | 37010000 |
| A0A8I4SYN6 | H3f4;H3-3b;H3c15; | 15.429 | 34.28 | 2 | 3 | 3955200 | 34188000 |
| P29341 | Pabpc1 | 70.67 | 33.338 | 20 | 19 | 16494000 | 19737000 |
| P53026 | Rpl10a | 24.916 | 33.198 | 7 | 8 | 42508000 | 81786000 |
| Q8C1B7 | Septin11 | 49.694 | 32.551 | 6 | 3 | 3853000 | 2448000 |
| Q8VHM5 | Hnrnpr | 70.887 | 32.149 | 9 | 9 | 2094600 | 4216500 |
| Q99JX4 | Eif3m | 42.516 | 31.976 | 2 | 3 | 810780 | 1170100 |
| Q8BWT1 | Acaa2 | 41.829 | 31.835 | 7 | 8 | 6926700 | 11207000 |
| Q8BP47 | NARS1 | 64.279 | 31.735 | 4 | 6 | 1924800 | 2593300 |
| Q99020 | Hnrnpab | 30.831 | 31.215 | 12 | 11 | 51221000 | 64842000 |
| Q9Z1D1 | Eif3g | 35.638 | 31.12 | 5 | 5 | 3286400 | 2260700 |
| Q8CGC7 | Eprs1 | 170.08 | 30.949 | 12 | 13 | 1147800 | 2068800 |
| P57780 | Actn4 | 104.98 | 30.705 | 13 | 16 | 1975900 | 3749600 |
| P11725 | Otc | 39.764 | 30.444 | 1 | 1 | 783460 | 557010 |
| Q8BMS1 | Hadha | 82.669 | 30.289 | 11 | 12 | 10157000 | 11638000 |
| P25444 | Rps2 | 31.231 | 30.281 | 6 | 8 | 29289000 | 47616000 |

| | | | | | | | |
|---|---|---|---|---|---|---|---|
| Q9D8N0 | Eef1g | 50.06 | 30.199 | 7 | 7 | 10651000 | 25395000 |
| P62317 | Snrpd2 | 13.527 | 29.985 | 4 | 5 | 15023000 | 25704000 |
| P70168 | Kpnb1 | 97.183 | 29.784 | 13 | 13 | 4584800 | 8079300 |
| Q64737 | Gart | 107.5 | 29.701 | 8 | 10 | 973710 | 1773600 |
| P62270 | Rps18;Rps18-ps6; | 17.718 | 29.697 | 11 | 12 | 41755000 | 67210000 |
| Q9CZ13 | Uqcrc1 | 52.851 | 29.438 | 3 | 4 | 1875600 | 3484800 |
| Q9DBG6 | Rpn2 | 69.062 | 29.264 | 5 | 4 | 1232000 | 1005700 |
| P62830 | Rpl23 | 14.865 | 29.153 | 3 | 4 | 2897200 | 7600900 |
| Q3TXS7 | Psmd1 | 105.73 | 29.018 | 5 | 6 | 687070 | 904970 |
| P97311 | Mcm6 | 92.866 | 28.903 | 11 | 13 | 2704000 | 5488800 |
| P42932 | Cct8 | 59.555 | 28.838 | 20 | 20 | 18459000 | 28062000 |
| P50580 | Pa2g4 | 43.698 | 28.484 | 7 | 7 | 9272000 | 12219000 |
| Q8BVY0 | Rsl1d1 | 50.421 | 28.47 | 7 | 9 | 8635200 | 21764000 |
| Q68FD5 | Cltc | 191.55 | 28.463 | 14 | 17 | 1252700 | 3367200 |
| P26516 | Psmd7 | 36.539 | 28.278 | 3 | 4 | 3408000 | 7018400 |
| P60335 | Pcbp1 | 37.497 | 28.247 | 6 | 5 | 8515400 | 14040000 |
| P62264 | Rps14 | 16.273 | 28.238 | 5 | 5 | 67855000 | 99616000 |
| P50518 | Atp6v1e1 | 26.157 | 28.068 | 0 | 1 | 0 | 261440 |
| Q8BMF4 | Dlat | 67.941 | 27.922 | 3 | 5 | 657210 | 1505000 |
| Q9Z1X4 | Ilf3 | 96.02 | 27.899 | 7 | 6 | 1135000 | 1899000 |
| Q9Z1R9 | Prss1 | 26.134 | 27.703 | 1 | 1 | 4136800000 | 3870100000 |
| Q9WU78 | Pdcd6ip | 96.023 | 27.695 | 4 | 5 | 836580 | 1580200 |
| Q9JJI8 | Rpl38 | 8.2038 | 27.669 | 1 | 2 | 7264300 | 21412000 |
| Q61990 | Pcbp2 | 38.221 | 27.296 | 7 | 5 | 14840000 | 19864000 |
| Q99JY0 | Hadhb | 51.386 | 27.093 | 9 | 8 | 4963400 | 7096800 |

| | | | | | | | |
|---|---|---|---|---|---|---|---|
| Q64012 | Raly | 33.188 | 26.723 | 5 | 5 | 4204000 | 6712300 |
| O08756 | Hsd17b10 | 27.418 | 26.409 | 3 | 2 | 816710 | 2341700 |
| P18760 | Cfl1 | 18.559 | 26.294 | 3 | 4 | 8383400 | 43077000 |
| Q99L45 | Eif2s2 | 38.092 | 26.199 | 4 | 4 | 1664200 | 4310300 |
| P62754 | Rps6 | 28.68 | 26.069 | 5 | 7 | 26537000 | 35068000 |
| Q01853 | Vcp | 89.321 | 26.02 | 9 | 13 | 3781400 | 5826600 |
| Q8BTM8 | Flna | 281.22 | 25.963 | 16 | 18 | 803800 | 1205600 |
| Q62167 | Ddx3x;D1Pas1 | 73.101 | 25.945 | 15 | 16 | 16529000 | 15567000 |
| Q91V41 | Rab14 | 23.897 | 25.668 | 7 | 6 | 2625700 | 3104900 |
| Q8QZY1 | Eif3l | 66.612 | 25.595 | 7 | 9 | 1790200 | 4142600 |
| Q9Z1Q9 | Vars1 | 140.21 | 25.537 | 10 | 11 | 2794300 | 4283000 |
| O88685 | Psmc3 | 49.548 | 25.284 | 7 | 9 | 2033900 | 6634400 |
| P07742 | Rrm1 | 90.209 | 25.273 | 9 | 13 | 3042000 | 5218100 |
| P17182 | Eno1 | 47.14 | 25.153 | 6 | 7 | 6876400 | 6534700 |
| Q921M3 | Sf3b3 | 135.55 | 25.07 | 8 | 9 | 1524500 | 2728300 |
| Q6ZWN5 | Rps9 | 22.591 | 25.053 | 12 | 13 | 66710000 | 100050000 |
| Q5SUF2 | Luc7l3 | 51.45 | 24.871 | 2 | 3 | 950370 | 1116400 |
| P49718 | Mcm5 | 82.406 | 24.604 | 7 | 11 | 2471400 | 5210200 |
| Q9EQU5 | Set | 33.377 | 24.389 | 3 | 3 | 6135100 | 7573400 |
| P62702 | Rps4x | 29.597 | 24.312 | 11 | 11 | 52817000 | 92881000 |
| Q8C196 | Cps1 | 164.62 | 24.09 | 3 | 2 | 370940 | 241570 |
| P07724 | Alb | 68.692 | 23.777 | 4 | 4 | 16526000 | 22174000 |
| P63101 | Ywhaz | 27.771 | 23.767 | 8 | 8 | 11294000 | 22895000 |
| Q07813 | Bax | 21.394 | 23.579 | 4 | 4 | 4868000 | 7558900 |
| P21958 | Tap1 | 78.863 | 23.578 | 2 | 3 | 791720 | 934950 |

| | | | | | | | |
|---|---|---|---|---|---|---|---|
| Q9WTM5 | Ruvbl2 | 51.112 | 23.551 | 8 | 13 | 3126700 | 7877900 |
| Q8BK67 | Rcc2 | 55.983 | 23.244 | 5 | 6 | 723330 | 2300600 |
| P20152 | Vim | 53.687 | 22.839 | 11 | 13 | 4868100 | 6021700 |
| Q64152 | Btf3 | 22.031 | 22.821 | 2 | 2 | 744060 | 1278300 |
| Q8VDN2 | Atp1a1 | 112.98 | 22.776 | 8 | 10 | 2554300 | 3824100 |
| O89053 | Coro1a | 50.989 | 22.759 | 9 | 12 | 13914000 | 20923000 |
| Q6PE01 | Snrnp40 | 39.275 | 22.71 | 1 | 2 | 1463000 | 967950 |
| Q8BKC5 | Ipo5 | 123.59 | 22.615 | 10 | 9 | 2266600 | 3047900 |
| Q9DB77 | Uqcrc2 | 48.234 | 22.44 | 7 | 7 | 4841900 | 9483800 |
| Q9CYG7 | Tomm34 | 34.278 | 21.832 | 5 | 5 | 2671300 | 3211800 |
| Q8CIN4 | Pak2 | 57.93 | 21.714 | 5 | 6 | 2304800 | 1948200 |
| Q8BJW6 | Eif2a | 64.403 | 21.574 | 4 | 6 | 1914200 | 4961100 |
| P08113 | Hsp90b1 | 92.475 | 21.549 | 13 | 13 | 2652300 | 6314400 |
| Q9WUA3 | Pfkp | 85.454 | 21.064 | 9 | 12 | 4921000 | 8735400 |
| P28656 | Nap1l1 | 45.345 | 20.83 | 6 | 6 | 20762000 | 38071000 |
| Q8BFR5 | Tufm | 49.508 | 20.725 | 11 | 10 | 6078600 | 7365200 |
| Q62293 | Tgtp1;Tgtp2 | 47.12 | 20.181 | 7 | 6 | 5589200 | 4762200 |
| Q3U9G9 | Lbr | 71.439 | 20.024 | 4 | 5 | 1849200 | 3387600 |
| Q64674 | Srm | 33.995 | 19.888 | 2 | 3 | 604370 | 843970 |
| Q9CPQ3 | Tomm22 | 15.537 | 19.805 | 1 | 2 | 366160 | 4790700 |
| P40124 | Cap1 | 51.564 | 19.677 | 6 | 7 | 3684900 | 4832900 |
| P62962 | Pfn1 | 14.957 | 19.613 | 5 | 8 | 12787000 | 17954000 |
| Q9CZM2 | Rpl15 | 24.146 | 19.572 | 0 | 1 | 0 | 68916 |
| Q922B2 | Dars1 | 57.147 | 19.516 | 6 | 10 | 1112700 | 5408300 |
| P84099 | Rpl19 | 23.466 | 19.453 | 6 | 7 | 33298000 | 29522000 |

| | | | | | | | |
|---|---|---|---|---|---|---|---|
| Q6ZWY9 | H2bc8;Hist2 h2bb; | 13.906 | 19.377 | 5 | 6 | 139860000 | 302480000 |
| Q91VR5 | Ddx1 | 82.499 | 19.34 | 6 | 7 | 1124700 | 2295300 |
| P26039 | Tln1 | 269.82 | 19.27 | 16 | 14 | 1245700 | 1330900 |
| Q9CZN7 | Shmt2 | 55.758 | 18.991 | 5 | 8 | 2754700 | 8454600 |
| Q60865 | Caprin1 | 78.168 | 18.922 | 4 | 3 | 4348100 | 3931500 |
| P12970 | Rpl7a | 29.976 | 18.779 | 10 | 10 | 24827000 | 60552000 |
| Q68FL6 | Mars1 | 101.43 | 18.609 | 6 | 9 | 1432300 | 2460500 |
| P09411 | Pgk1 | 44.55 | 18.548 | 5 | 4 | 990560 | 884490 |
| Q91YQ5 | Rpn1 | 68.527 | 18.506 | 5 | 8 | 2243000 | 3556200 |
| O35286 | Dhx15 | 91.006 | 18.472 | 8 | 7 | 1695500 | 2167200 |
| P62137 | Ppp1ca | 37.54 | 18.33 | 4 | 5 | 761580 | 3531500 |
| P25206 | Mcm3 | 91.545 | 18.265 | 13 | 17 | 5161900 | 8014200 |
| O55131 | Septin7 | 50.549 | 18.153 | 7 | 8 | 4831600 | 5840000 |
| A8DUK4 | Hbb-bs;Hbb-b1 | 15.748 | 18.138 | 5 | 5 | 5038400 | 12154000 |
| Q3U0V1 | Khsrp | 76.775 | 18.11 | 8 | 7 | 2134500 | 2516800 |
| Q9D880 | Timm50 | 39.776 | 18.068 | 3 | 4 | 2382400 | 4869400 |
| Q9D6R2 | Idh3a | 39.638 | 17.931 | 5 | 5 | 2419000 | 4234800 |
| P97855 | G3bp1 | 51.828 | 17.756 | 9 | 9 | 6802800 | 8542100 |
| Q9CXW4 | Rpl11 | 20.252 | 17.685 | 3 | 4 | 30949000 | 51843000 |
| Q922K7 | Nop2 | 86.751 | 17.493 | 6 | 8 | 1268100 | 2758300 |
| Q8CAQ8 | Immt | 83.899 | 17.462 | 9 | 7 | 470450 | 953250 |
| P15864 | H1-2 | 21.266 | 17.346 | 7 | 11 | 28193000 | 95587000 |
| Q8R081 | Hnrnpl | 63.963 | 17.319 | 8 | 8 | 5219800 | 9061600 |
| P14131 | Rps16 | 16.445 | 17.31 | 9 | 10 | 36901000 | 55823000 |

| | | | | | | | |
|---|---|---|---|---|---|---|---|
| Q99JI4 | Psmd6 | 45.536 | 17.218 | 6 | 8 | 1189200 | 2050300 |
| Q91YE6 | Ipo9 | 116.05 | 17.212 | 0 | 1 | 0 | 371990 |
| Q9DBR1 | Xrn2 | 108.69 | 17.13 | 7 | 5 | 1283200 | 1244700 |
| P54775 | Psmc4 | 47.408 | 17.116 | 6 | 8 | 438920 | 1711900 |
| P35564 | Canx | 67.277 | 16.931 | 6 | 5 | 3848300 | 1241700 |
| P49717 | Mcm4 | 96.735 | 16.912 | 10 | 15 | 1370200 | 4165200 |
| P62814 | Atp6v1b2 | 56.55 | 16.861 | 4 | 3 | 410450 | 332140 |
| Q99MN1 | Kars1 | 67.839 | 16.738 | 9 | 11 | 4115400 | 8971000 |
| Q62351 | Tfrc | 85.73 | 16.724 | 8 | 7 | 1423600 | 2559600 |
| Q8BWY3 | Etf1 | 49.03 | 16.635 | 5 | 6 | 2782800 | 5454600 |
| Q9CR57 | Rpl14 | 23.564 | 16.633 | 4 | 4 | 17088000 | 26678000 |
| Q9DCH4 | Eif3f | 37.984 | 16.491 | 6 | 5 | 9981200 | 13108000 |
| Q08288 | Lyar | 43.735 | 16.33 | 5 | 6 | 2415800 | 4385500 |
| Q8BQ46 | Taf15 | 58.6 | 16.005 | 4 | 4 | 264620 | 123760 |
| P35922 | Fmr1 | 68.988 | 15.969 | 1 | 0 | 0 | 0 |
| P97310 | Mcm2 | 102.08 | 15.852 | 10 | 10 | 2132900 | 3818700 |
| P32848 | Pvalb | 11.93 | 15.609 | 1 | 3 | 841410 | 988910 |
| P62821 | Rab1A | 22.677 | 15.509 | 3 | 3 | 4064600 | 5962600 |
| Q61316 | Hspa4 | 94.132 | 15.323 | 4 | 5 | 736630 | 1449600 |
| Q6GSS7 | Hist2h2aa2; H2ac20; | 14.095 | 15.246 | 4 | 6 | 115410000 | 346190000 |
| O54825 | Bysl | 49.783 | 15.197 | 2 | 2 | 0 | 951200 |
| P51174 | Acadl | 47.907 | 15.105 | 3 | 3 | 1306200 | 581010 |
| P61089 | Ube2n | 17.138 | 15.031 | 2 | 3 | 4489100 | 5389100 |
| Q8BHD7 | Ptbp3 | 56.7 | 15.026 | 2 | 3 | 470780 | 551320 |
| P60867 | Rps20 | 13.373 | 15.008 | 4 | 4 | 36846000 | 65166000 |

| | | | | | | | |
|---|---|---|---|---|---|---|---|
| O08583 | Alyref;Alyref2 | 26.94 | 14.862 | 4 | 4 | 9112300 | 13078000 |
| Q9CPN7 | 1810009J06 Rik;Gm2663 | 26.503 | 14.79 | 1 | 1 | 5360600 | 10219000 |
| Q9JKP5 | Mbnl1 | 36.975 | 14.736 | 3 | 4 | 1526700 | 2952300 |
| Q8K1B8 | Fermt3 | 75.634 | 14.729 | 7 | 6 | 2583800 | 2426000 |
| Q6ZWU9 | Rps27 | 9.461 | 14.685 | 4 | 3 | 23086000 | 31262000 |
| Q8BG32 | Psmd11 | 47.436 | 14.581 | 10 | 10 | 3448400 | 6896400 |
| Q8BMJ2 | Lars1 | 134.19 | 14.472 | 9 | 12 | 1698100 | 3025000 |
| P13020 | Gsn | 85.941 | 14.41 | 0 | 1 | 0 | 0 |
| P36371 | Tap2 | 77.444 | 14.399 | 3 | 3 | 409150 | 471780 |
| P23116 | Eif3a | 161.93 | 14.382 | 16 | 15 | 2641900 | 3729300 |
| P07356 | Anxa2 | 38.676 | 14.369 | 9 | 12 | 3468500 | 12787000 |
| P67778 | Phb1 | 29.82 | 14.245 | 7 | 7 | 6693100 | 11002000 |
| Q8BU30 | Iars1 | 144.27 | 14.137 | 4 | 4 | 483000 | 810150 |
| O70194 | Eif3d | 63.988 | 14.136 | 4 | 5 | 2457400 | 4174500 |
| Q8BY71 | Hat1 | 49.278 | 14.044 | 3 | 3 | 1680900 | 3318500 |
| Q9Z1Q5 | Clic1 | 27.013 | 14.029 | 3 | 6 | 2500900 | 4410500 |
| P62334 | Psmc6 | 44.172 | 13.964 | 8 | 9 | 3870900 | 7124900 |
| Q9CQE8 | RTRAF | 28.152 | 13.885 | 2 | 2 | 2163500 | 2484900 |
| P99027 | Rplp2;Rplp2-ps1 | 11.651 | 13.871 | 5 | 6 | 19945000 | 38915000 |
| Q8JZQ9 | Eif3b | 91.369 | 13.775 | 6 | 9 | 2784900 | 5648900 |
| P80313 | Cct7 | 59.652 | 13.565 | 10 | 12 | 9831300 | 16784000 |
| P97371 | Psme1 | 28.673 | 13.5 | 7 | 7 | 12255000 | 12550000 |
| O35737 | Hnrnph1 | 49.199 | 13.393 | 9 | 7 | 12756000 | 11681000 |
| P14148 | Rpl7 | 31.419 | 13.383 | 10 | 11 | 19962000 | 50900000 |

| | | | | | | | |
|---|---|---|---|---|---|---|---|
| Q9CX86 | Hnrnpa0 | 30.53 | 13.331 | 6 | 11 | 7531200 | 14344000 |
| Q60737 | Csnk2a1 | 45.133 | 13.293 | 2 | 4 | 774970 | 1628000 |
| Q9JIX8 | Acin1 | 150.72 | 13.007 | 1 | 3 | 0 | 0 |
| Q8BPU7 | Elmo1 | 83.935 | 12.708 | 4 | 4 | 548600 | 756710 |
| Q9Z110 | Aldh18a1 | 87.265 | 12.686 | 9 | 9 | 2368600 | 4166400 |
| Q9JKB3 | Ybx3 | 38.813 | 12.683 | 4 | 2 | 2257000 | 1394500 |
| Q3THK3 | Gtf2f1 | 57.241 | 12.554 | 1 | 2 | 0 | 538490 |
| P70372 | Elavl1 | 36.169 | 12.484 | 6 | 7 | 7744400 | 13273000 |
| Q569Z6 | Thrap3 | 108.18 | 12.433 | 11 | 8 | 5591400 | 3591700 |
| Q921F2 | Tardbp | 44.547 | 12.354 | 5 | 5 | 11284000 | 15503000 |
| Q6ZWV3 | Rpl10;Rpl10l | 24.604 | 12.336 | 5 | 7 | 15403000 | 34446000 |
| Q9ERK4 | Cse1l | 110.45 | 12.307 | 8 | 6 | 12061000 | 1543500 |
| Q9QZD9 | Eif3i | 36.46 | 12.275 | 6 | 4 | 1478900 | 2128300 |
| Q99MD9 | Nasp | 83.953 | 12.266 | 0 | 1 | 0 | 99365 |
| Q922R8 | Pdia6 | 48.1 | 12.128 | 5 | 7 | 3600800 | 7700900 |
| Q62093 | Srsf2 | 25.476 | 11.997 | 3 | 2 | 4538300 | 5743900 |
| Q8VDM4 | Psmd2 | 100.2 | 11.965 | 9 | 12 | 1559600 | 3573100 |
| Q9CPS7 | Pno1 | 27.453 | 11.894 | 1 | 2 | 1578300 | 3069700 |
| Q921M7 | Cyrib | 36.776 | 11.852 | 2 | 5 | 1348000 | 3350800 |
| O09167 | Rpl21 | 18.579 | 11.839 | 4 | 4 | 12566000 | 34357000 |
| Q99LF4 | Rtcb | 55.249 | 11.786 | 8 | 9 | 3289700 | 5669200 |
| P62267 | Rps23 | 15.807 | 11.777 | 3 | 3 | 27118000 | 37311000 |
| P61164 | Actr1a | 42.613 | 11.774 | 1 | 2 | 1409200 | 2024000 |
| P61222 | Abce1 | 67.314 | 11.752 | 7 | 5 | 1847800 | 2601900 |
| P35278 | Rab5c;Rab5a; | 23.412 | 11.714 | 4 | 4 | 1994900 | 3964500 |

| | | | | | | | |
|---|---|---|---|---|---|---|---|
| P32067 | Ssb | 47.756 | 11.684 | 3 | 3 | 1497900 | 2034200 |
| Q9CQ43 | Dut | 17.384 | 11.676 | 0 | 2 | 0 | 0 |
| Q9D903 | Ebna1bp2 | 34.702 | 11.665 | 3 | 6 | 449700 | 3152800 |
| A0A140T8M0 | Igkv1-117;Igkv1-110; | 13.117 | 11.651 | 3 | 3 | 267940000 | 151920000 |
| Q9R0N0 | Galk1 | 42.295 | 11.622 | 2 | 2 | 2181000 | 4195900 |
| P63073 | Eif4e | 25.053 | 11.609 | 2 | 3 | 1108500 | 1758900 |
| Q9CZX8 | Rps19 | 16.085 | 11.549 | 7 | 8 | 23662000 | 46649000 |
| P13439 | Umps | 52.292 | 11.497 | 4 | 4 | 1346600 | 2446900 |
| Q60972 | Rbbp4 | 47.655 | 11.425 | 2 | 3 | 2119000 | 2939400 |
| O70456 | Sfn | 27.706 | 11.228 | 4 | 10 | 4135000 | 21093000 |
| D3Z7P3 | Gls | 73.963 | 11.175 | 2 | 2 | 625640 | 1173000 |
| Q99ME9 | Gtpbp4 | 74.112 | 11.101 | 5 | 9 | 801000 | 3341900 |
| P60710 | Actb | 41.736 | 11.081 | 21 | 25 | 4852400 | 10340000 |
| P30416 | Fkbp4 | 51.572 | 11.08 | 4 | 6 | 503240 | 1067300 |
| Q501J6 | Ddx17 | 72.399 | 11.078 | 5 | 8 | 188230 | 847310 |
| P62082 | Rps7 | 22.127 | 11.072 | 5 | 7 | 10119000 | 17888000 |
| Q8K310 | Matr3 | 94.629 | 11.027 | 8 | 9 | 2100700 | 3739000 |
| Q6ZQ58 | Larp1 | 121.12 | 11.024 | 7 | 6 | 2702900 | 2356700 |
| Q8VH51 | Rbm39 | 59.406 | 10.946 | 4 | 4 | 3422600 | 4746700 |
| P27773 | Pdia3 | 56.678 | 10.862 | 12 | 13 | 3943500 | 7814600 |
| Q9DCL9 | Paics | 47.006 | 10.806 | 7 | 7 | 4670200 | 6421100 |
| Q3THE2 | Myl12b;Myl9 | 19.779 | 10.794 | 6 | 8 | 16741000 | 25315000 |
| Q6NZJ6 | Eif4g1 | 176.07 | 10.79 | 13 | 14 | 1237700 | 1995000 |
| Q61753 | Phgdh | 56.585 | 10.783 | 7 | 9 | 20600000 | 27599000 |

| | | | | | | | |
|---|---|---|---|---|---|---|---|
| Q61599 | Arhgdib | 22.851 | 10.752 | 4 | 3 | 3902300 | 4738300 |
| Q9D2G2 | Dlst | 48.994 | 10.688 | 3 | 2 | 678460 | 1681600 |
| Q6DFW4 | Nop58 | 60.342 | 10.634 | 8 | 8 | 3835900 | 8722700 |
| Q9EQ61 | Pes1 | 67.795 | 10.499 | 4 | 4 | 1045900 | 2508100 |
| P06151 | Ldha | 36.498 | 10.496 | 7 | 8 | 8401700 | 10946000 |
| Q60749 | Khdrbs1 | 48.37 | 10.447 | 5 | 5 | 12293000 | 11205000 |
| Q8C2Q3 | Rbm14 | 69.448 | 10.433 | 11 | 10 | 7710300 | 3683100 |
| Q9D051 | Pdhb | 38.937 | 10.399 | 5 | 5 | 2243600 | 5116700 |
| P40142 | Tkt | 67.63 | 10.375 | 8 | 8 | 4369700 | 3951000 |
| P60122 | Ruvbl1 | 50.213 | 10.338 | 7 | 8 | 5940800 | 11855000 |
| Q99MR6 | Srrt | 100.45 | 10.182 | 8 | 6 | 3061000 | 3486500 |
| P47962 | Rpl5 | 34.4 | 10.168 | 7 | 10 | 13740000 | 26578000 |
| Q9DB20 | Atp5po | 23.363 | 10.104 | 5 | 6 | 6420800 | 13719000 |
| P24547 | Impdh2 | 55.814 | 10.099 | 7 | 8 | 2461800 | 5510900 |
| Q8R1B4 | Eif3c | 105.53 | 9.9791 | 5 | 8 | 2104300 | 5797600 |
| P42209 | Septin1 | 42.019 | 9.9547 | 5 | 4 | 15659000 | 30871000 |
| Q91VA7 | Idh3b | 42.194 | 9.9378 | 9 | 12 | 5272300 | 9883700 |
| Q60932 | Vdac1 | 32.351 | 9.8745 | 4 | 4 | 1126900 | 4135200 |
| P05064 | Aldoa;Aldoart1 | 39.355 | 9.632 | 4 | 6 | 2568700 | 5417200 |
| O08553 | Dpysl2 | 62.277 | 9.6018 | 5 | 5 | 1653100 | 1855000 |
| P62849 | Rps24 | 15.423 | 9.5686 | 5 | 5 | 41771000 | 49215000 |
| P62301 | Rps13 | 17.222 | 9.5537 | 7 | 8 | 35007000 | 57512000 |
| Q6P5F9 | Xpo1 | 123.09 | 9.499 | 3 | 4 | 290160 | 727380 |
| P46061 | Rangap1 | 63.53 | 9.4776 | 1 | 4 | 449450 | 491810 |
| Q64324 | Stxbp2 | 66.357 | 9.4733 | 1 | 1 | 0 | 0 |

| | | | | | | | |
|---|---|---|---|---|---|---|---|
| P47964 | Rpl36 | 12.254 | 9.3949 | 2 | 3 | 10493000 | 17095000 |
| Q6PDQ2 | Chd4 | 217.75 | 9.3756 | 4 | 5 | 392380 | 672720 |
| Q76MZ3 | Ppp2r1a | 65.322 | 9.2705 | 5 | 5 | 668820 | 2615300 |
| Q5SUA5 | Myo1g | 117.23 | 9.2454 | 4 | 6 | 126390 | 490310 |
| P32921 | Wars1 | 54.357 | 9.2378 | 3 | 3 | 715260 | 1182300 |
| Q60973 | Rbbp7 | 47.789 | 9.0828 | 3 | 5 | 1764800 | 5383200 |
| O35350 | Capn1 | 82.105 | 9.0783 | 7 | 6 | 2766900 | 1947500 |
| P35550 | Fbl | 34.306 | 8.8778 | 6 | 11 | 3221400 | 8882400 |
| Q9EQH3 | Vps35 | 91.712 | 8.7188 | 4 | 3 | 457910 | 929840 |
| P35979 | Rpl12 | 17.804 | 8.6782 | 3 | 3 | 14464000 | 29149000 |
| Q3U7R1 | Esyt1 | 121.55 | 8.6655 | 5 | 6 | 677580 | 865470 |
| Q3V3R1 | Mthfd1l | 105.73 | 8.6598 | 5 | 10 | 1480500 | 3092300 |
| Q91YK2 | Rrp1b | 80.581 | 8.6445 | 0 | 2 | 0 | 147850 |
| Q8VDW0 | Ddx39a | 49.067 | 8.587 | 9 | 8 | 11323000 | 20812000 |
| P47856 | Gfpt1 | 78.538 | 8.5254 | 1 | 2 | 166480 | 323690 |
| Q60605 | Myl6 | 16.93 | 8.467 | 4 | 4 | 9317000 | 16707000 |
| Q61545 | Ewsr1 | 68.461 | 8.4235 | 5 | 4 | 5871700 | 4032200 |
| Q8CI11 | Gnl3 | 60.786 | 8.3699 | 2 | 5 | 782750 | 2059500 |
| Q9WVK4 | Ehd1 | 60.602 | 8.3037 | 5 | 5 | 1338100 | 1643000 |
| Q920B9 | Supt16h | 119.82 | 8.267 | 5 | 8 | 772960 | 2619000 |
| Q9CXW3 | Cacybp | 26.51 | 8.2662 | 4 | 5 | 2216900 | 3202300 |
| Q99M31 | Hspa14 | 54.65 | 8.2217 | 2 | 4 | 529590 | 1150100 |
| P09103 | P4hb | 57.058 | 8.1883 | 4 | 3 | 1598700 | 2534600 |
| Q99KP6 | Prpf19 | 55.238 | 8.178 | 7 | 6 | 4200800 | 5185200 |
| P46471 | Psmc2 | 48.647 | 8.1503 | 8 | 10 | 1955300 | 5062300 |
| O08992 | Sdcbp | 32.379 | 8.1493 | 1 | 0 | 0 | 0 |

| | | | | | | | |
|---|---|---|---|---|---|---|---|
| Q9CZD3 | Gars1 | 81.877 | 8.0925 | 5 | 5 | 1558400 | 2340400 |
| P42669 | Pura | 34.883 | 8.0696 | 2 | 2 | 324960 | 585990 |
| P63242 | Eif5a;Eif5a2 | 16.832 | 8.0566 | 2 | 2 | 8645300 | 19570000 |
| P97372 | Psme2 | 27.057 | 8.0431 | 5 | 5 | 6435800 | 12650000 |
| Q9Z204 | Hnrnpc | 34.384 | 8.0182 | 7 | 8 | 8842800 | 15038000 |
| Q8R3C7 | Ddx19b;Ddx 19a | 53.893 | 7.9127 | 3 | 3 | 1571200 | 1675400 |
| Q792Z1 | Try10 | 26.221 | 7.8798 | 1 | 1 | 378620000 | 425320000 |
| O08810 | Eftud2 | 109.36 | 7.7977 | 11 | 14 | 1808900 | 2678300 |
| O35129 | Phb2 | 33.296 | 7.797 | 6 | 5 | 4811900 | 10149000 |
| Q01320 | Top2a | 172.79 | 7.7562 | 5 | 7 | 402800 | 1268200 |
| P47757 | Capzb | 31.345 | 7.7532 | 2 | 4 | 293800 | 4630700 |
| P01897 | H2-L; | 40.711 | 7.6859 | 6 | 6 | 7673600 | 9305100 |
| Q99J36 | Thumpd1 | 38.884 | 7.6567 | 1 | 2 | 632560 | 1471600 |
| Q8VEE4 | Rpa1 | 69.036 | 7.608 | 4 | 3 | 837430 | 1538700 |
| P14685 | Psmd3 | 60.718 | 7.6072 | 8 | 9 | 1396100 | 3018500 |
| O55029 | Copb2 | 102.45 | 7.4969 | 3 | 4 | 491090 | 992610 |
| P68510 | Ywhah | 28.211 | 7.4889 | 3 | 4 | 845020 | 3452900 |
| O35857 | Timm44 | 51.091 | 7.397 | 3 | 4 | 1352600 | 1794600 |
| Q91Y97 | Aldob | 39.507 | 7.3926 | 1 | 0 | 589550 | 0 |
| Q6ZQ38 | Cand1 | 136.33 | 7.2998 | 5 | 4 | 296360 | 459120 |
| P35980 | Rpl18 | 21.644 | 7.2683 | 5 | 6 | 21372000 | 42394000 |
| P62855 | Rps26 | 13.015 | 7.2647 | 2 | 2 | 31201000 | 48797000 |
| P47915 | Rpl29 | 17.587 | 7.2499 | 1 | 1 | 122640 | 1535200 |
| Q9CQX2 | Cyb5b | 16.318 | 7.1872 | 1 | 1 | 1605900 | 4298100 |
| P61290 | Psme3 | 29.506 | 7.1354 | 5 | 4 | 2746700 | 3288300 |

| | | | | | | | |
|---|---|---|---|---|---|---|---|
| Q9Z2I8 | Suclg2 | 46.839 | 7.1124 | 4 | 4 | 2390700 | 3890500 |
| P52293 | Kpna2 | 57.927 | 6.9476 | 4 | 5 | 3342500 | 9237100 |
| Q8R2M2 | Dnttip2 | 84.276 | 6.924 | 0 | 1 | 0 | 0 |
| P39749 | Fen1 | 42.314 | 6.9106 | 3 | 3 | 653270 | 667780 |
| A0A0A6YVU 8;Q9JKV1 | Adrm1b;Adr m1 | 42.147 | 6.9001 | 2 | 1 | 3156300 | 2513000 |
| P01831 | Thy1 | 18.08 | 6.862 | 4 | 4 | 14385000 | 24393000 |
| P61028;Q9D D03;Q8K386 | Rab8b | 23.603 | 6.8042 | 4 | 4 | 6385500 | 7335900 |
| Q8CGK3 | Lonp1 | 105.84 | 6.8019 | 3 | 3 | 153180 | 717030 |
| Q9EPL8 | Ipo7 | 119.49 | 6.7748 | 5 | 5 | 1117400 | 1497600 |
| Q9WTL4 | Insrr | 144.87 | 6.7594 | 2 | 2 | 5422500 | 8085000 |
| P60229 | Eif3e | 52.22 | 6.7517 | 5 | 8 | 2707200 | 4217200 |
| Q99M28 | Rnps1 | 34.208 | 6.7425 | 1 | 2 | 1211600 | 1194500 |
| P62827;Q14 AA6;Q61820 | Ran;170000 9N14Rik; | 24.423 | 6.6959 | 4 | 6 | 7935100 | 26757000 |
| Q8R326 | Pspc1 | 58.758 | 6.6681 | 0 | 2 | 0 | 974490 |
| P29351 | Ptpn6 | 67.558 | 6.6385 | 10 | 10 | 3130600 | 5070600 |
| Q8C7V3 | Utp15 | 59.374 | 6.5211 | 0 | 3 | 0 | 777660 |
| Q91WJ8 | Fubp1 | 68.539 | 6.4753 | 5 | 5 | 1272200 | 1430400 |
| P51881;Q3V 132 | Slc25a5 | 32.931 | 6.4456 | 9 | 9 | 30418000 | 62981000 |
| Q78ZA7 | Nap1l4 | 42.679 | 6.3953 | 6 | 4 | 1740000 | 2199800 |
| Q99LC2 | Cstf1 | 48.381 | 6.2428 | 1 | 2 | 496940 | 635280 |
| P62245 | Rps15a | 14.839 | 6.2399 | 7 | 7 | 33234000 | 45535000 |
| P68040 | Rack1 | 35.076 | 6.1575 | 5 | 7 | 9684100 | 22754000 |

| | | | | | | | |
|---|---|---|---|---|---|---|---|
| Q921N6 | Ddx27 | 85.938 | 6.1472 | 3 | 6 | 634770 | 1782800 |
| P01786 | | 12.374 | 6.122 | 1 | 1 | 15607000 | 7585800 |
| O89086 | Rbm3 | 16.604 | 6.1045 | 5 | 4 | 10282000 | 24591000 |
| Q8R323 | Rfc3 | 40.526 | 6.046 | 2 | 2 | 803180 | 1210900 |
| Q99PV0 | Prpf8 | 273.61 | 6.0072 | 5 | 8 | 177890 | 340960 |
| P37913 | Lig1 | 102.29 | 5.9784 | 7 | 5 | 356270 | 734340 |
| P43276 | H1-5 | 22.576 | 5.9656 | 4 | 4 | 10623000 | 20279000 |
| Q9CPR4 | Rpl17 | 21.397 | 5.9342 | 5 | 6 | 7019600 | 14102000 |
| P48678 | Lmna | 74.237 | 5.9322 | 0 | 6 | 0 | 2218300 |
| O35654 | Pold2 | 51.354 | 5.9058 | 2 | 2 | 335360 | 879990 |
| P26041 | Msn | 67.766 | 5.8994 | 7 | 7 | 3036600 | 5217200 |
| Q91YU8 | Ppan | 52.755 | 5.8885 | 0 | 2 | 0 | 965570 |
| P08752 | Gnai2 | 40.489 | 5.8723 | 4 | 5 | 5299900 | 6491700 |
| Q91V92 | Acly | 119.73 | 5.8671 | 3 | 5 | 593570 | 1661500 |
| Q3TIX9 | Usp39 | 65.146 | 5.826 | 1 | 2 | 0 | 979090 |
| Q8BP67 | Rpl24 | 17.779 | 5.7884 | 4 | 4 | 28855000 | 68721000 |
| P43275 | H1-1 | 21.785 | 5.773 | 3 | 3 | 3737600 | 9145000 |
| P26369 | U2af2 | 53.516 | 5.7171 | 4 | 3 | 4931900 | 5061500 |
| P11157 | Rrm2 | 45.095 | 5.7013 | 3 | 3 | 2335300 | 3108000 |
| Q8BL97 | Srsf7 | 30.817 | 5.6699 | 1 | 3 | 1264400 | 2251700 |
| P61161 | Actr2 | 44.76 | 5.6383 | 4 | 5 | 3677100 | 5873600 |
| Q91VD9 | Ndufs1 | 79.776 | 5.6355 | 2 | 4 | 527200 | 652760 |
| Q9EPU0 | Upf1 | 123.97 | 5.5826 | 7 | 8 | 397300 | 680900 |
| P62880 | Gnb2;Gnb4 | 37.331 | 5.5722 | 3 | 4 | 4628000 | 9339200 |
| Q9QYJ0 | Dnaja2 | 45.745 | 5.5679 | 2 | 4 | 1059800 | 1389100 |
| Q91VE6 | Nifk | 36.265 | 5.5483 | 2 | 4 | 710860 | 3604500 |

| | | | | | | | |
|---|---|---|---|---|---|---|---|
| P11835 | Itgb2 | 85.025 | 5.5449 | 5 | 5 | 551170 | 1041300 |
| Q99LH1 | Gnl2 | 83.345 | 5.479 | 0 | 2 | 0 | 280300 |
| Q9QUJ7 | Acsl4 | 79.076 | 5.4727 | 1 | 3 | 423550 | 1161100 |
| P42227 | Stat3 | 88.053 | 5.4645 | 4 | 3 | 970560 | 217600 |
| Q61334 | Bcap29 | 27.964 | 5.4442 | 0 | 1 | 0 | 0 |
| P41105 | Rpl28 | 15.733 | 5.4265 | 3 | 5 | 6859500 | 28955000 |
| P26638 | Sars1 | 58.388 | 5.4168 | 3 | 3 | 920980 | 1106600 |
| P31230 | Aimp1 | 33.997 | 5.414 | 3 | 3 | 1608700 | 1597000 |
| P53994 | Rab2a;Rab2b | 23.547 | 5.3897 | 3 | 2 | 2398600 | 3026100 |
| Q99NB9 | Sf3b1 | 145.81 | 5.3085 | 3 | 6 | 99040 | 269810 |
| Q9D0I9 | Rars1 | 75.673 | 5.2482 | 7 | 8 | 1742800 | 3070300 |
| Q9CXY6 | Ilf2 | 43.062 | 5.2312 | 4 | 5 | 2956300 | 3707500 |
| Q9CW03 | Smc3 | 141.55 | 5.199 | 4 | 5 | 352990 | 860540 |
| O35593 | Psmd14 | 34.577 | 5.1463 | 2 | 3 | 2090100 | 4332400 |
| P00405 | Mtco2 | 25.976 | 5.0525 | 2 | 2 | 9158900 | 9499300 |
| Q3TJZ6 | Fam98a | 55.055 | 5.0298 | 2 | 2 | 1055900 | 705670 |
| Q8VDJ3 | Hdlbp | 141.74 | 5.0115 | 7 | 7 | 303310 | 574280 |
| P61924 | Copz1 | 20.198 | 4.9794 | 3 | 4 | 7987200 | 2094600 |
| Q8BML9 | Qars1 | 87.676 | 4.9755 | 6 | 7 | 617960 | 809980 |
| Q9DBE9 | Ftsj3 | 95.531 | 4.9564 | 3 | 3 | 355600 | 534580 |
| Q9CY57 | Chtop | 26.585 | 4.9153 | 3 | 3 | 780360 | 3061300 |
| Q8BK64 | Ahsa1 | 38.117 | 4.9129 | 2 | 3 | 0 | 566360 |
| O08797 | Serpinb9 | 42.259 | 4.8999 | 3 | 5 | 2393800 | 2395800 |
| P06240 | Lck | 57.942 | 4.8841 | 5 | 6 | 2480500 | 2340000 |
| P50516 | Atp6v1a | 68.325 | 4.8782 | 4 | 4 | 474170 | 1072400 |

| | | | | | | | |
|---|---|---|---|---|---|---|---|
| Q8CDN6 | Txnl1 | 32.237 | 4.8487 | 2 | 5 | 1416400 | 3694800 |
| Q8VEM8 | Slc25a3 | 39.632 | 4.8382 | 3 | 3 | 4336900 | 12181000 |
| Q9DBG3 | Ap2b1;Ap1b1 | 104.58 | 4.8169 | 3 | 4 | 812650 | 1192300 |
| P84104 | Srsf3 | 19.329 | 4.8135 | 2 | 2 | 5712800 | 22125000 |
| Q9R0T7 | Try4;Try5 | 26.274 | 4.7705 | 1 | 1 | 2456800 | 1717300 |
| Q6P542 | Abcf1 | 94.944 | 4.7623 | 4 | 4 | 569340 | 1166100 |
| P62852 | Rps25 | 13.742 | 4.7568 | 6 | 6 | 105110000 | 167540000 |
| Q8BKS9 | Pum3 | 72.799 | 4.7521 | 0 | 3 | 0 | 310290 |
| Q64310 | Surf4 | 30.381 | 4.7336 | 1 | 2 | 926250 | 2034300 |
| Q99L43 | Cds2 | 51.313 | 4.6857 | 1 | 1 | 862850 | 779230 |
| Q9D554 | Sf3a3 | 58.841 | 4.6485 | 4 | 4 | 2381000 | 2890700 |
| Q6ZWV7 | Rpl35 | 14.552 | 4.6315 | 3 | 4 | 14530000 | 15282000 |
| P11103 | Parp1 | 113.1 | 4.6179 | 1 | 3 | 197550 | 483220 |
| Q9R233 | Tapbp | 49.736 | 4.6119 | 3 | 3 | 2642500 | 4203200 |
| Q8CG48 | Smc2 | 134.24 | 4.5379 | 4 | 3 | 216380 | 307770 |
| P01942 | Hba | 15.085 | 4.4644 | 5 | 5 | 125000000 | 120450000 |
| P14824 | Anxa6 | 75.884 | 4.4565 | 3 | 3 | 173440 | 190940 |
| Q9JIF7 | Copb1 | 107.06 | 4.4561 | 4 | 5 | 352650 | 867470 |
| P43274 | H1-4 | 21.977 | 4.45 | 5 | 7 | 9287600 | 21726000 |
| Q8CG47 | Smc4 | 146.89 | 4.422 | 2 | 3 | 88371 | 134490 |
| Q9CWX9 | Ddx47 | 50.638 | 4.32 | 6 | 6 | 2006500 | 3499700 |
| Q7TSF1 | Dsg1b;Dsg1a | 114.45 | 4.3153 | 2 | 3 | 867130 | 2245800 |
| Q9CRB2 | Nhp2 | 17.247 | 4.2911 | 1 | 1 | 2004500 | 3672100 |
| P42208 | Septin2 | 41.525 | 4.2762 | 3 | 3 | 1851700 | 2964100 |

| | | | | | | | |
|---|---|---|---|---|---|---|---|
| Q9DCS9 | Ndufb10 | 21.024 | 4.2689 | 0 | 1 | 0 | 998110 |
| Q8K019 | Bclaf1 | 106 | 4.2679 | 7 | 4 | 852520 | 1095100 |
| P84084 | Arf5 | 20.529 | 4.234 | 3 | 2 | 7222400 | 7504700 |
| P55258 | Rab8a | 23.668 | 4.2325 | 2 | 2 | 1043200 | 2051900 |
| P06745 | Gpi | 62.766 | 4.2094 | 2 | 1 | 188450 | 224930 |
| D3Z2H9 | Tpm3-rs7;Tpm2; | 28.991 | 4.2033 | 4 | 4 | 2305800 | 3240800 |
| Q8K2F8 | Lsm14a | 50.545 | 4.1766 | 2 | 0 | 280220 | 0 |
| P54071 | Idh2 | 50.906 | 4.1698 | 0 | 3 | 0 | 546690 |
| Q9Z2I0 | Letm1 | 82.988 | 4.1569 | 3 | 3 | 784230 | 1320500 |
| P01901 | H2-K1 | 41.301 | 4.1495 | 6 | 5 | 2406800 | 2114200 |
| Q99L47 | St13 | 41.655 | 4.1485 | 3 | 3 | 2099700 | 4455700 |
| Q9JHI7 | Exosc9 | 48.936 | 4.1389 | 1 | 1 | 247370 | 390520 |
| P97379 | G3bp2 | 54.087 | 4.1371 | 3 | 3 | 406430 | 811940 |
| P19253 | Rpl13a | 23.464 | 4.133 | 5 | 7 | 10812000 | 21313000 |
| Q9CQR2 | Rps21 | 9.1413 | 4.1248 | 2 | 2 | 9125100 | 8170700 |
| Q922P9 | Glyr1 | 59.715 | 4.0822 | 1 | 1 | 289420 | 394370 |
| P46638;P62492 | Rab11b;Rab11a | 24.489 | 4.0725 | 3 | 3 | 983220 | 3539000 |
| Q9WVJ2 | Psmd13 | 42.809 | 4.0655 | 1 | 4 | 267150 | 2896000 |
| Q9JKX6 | Nudt5 | 23.984 | 4.0614 | 4 | 4 | 1953700 | 3875800 |
| A9R9W0 | Mup15;Mup14; | 20.647 | 4.0414 | 3 | 2 | 2127700 | 1269200 |
| Q61074 | Ppm1g | 58.727 | 4.0411 | 1 | 3 | 0 | 1582000 |
| Q8BMC4 | Nop9 | 70.046 | 4.0289 | 1 | 1 | 115930 | 0 |

| | | | | | | | |
|---|---|---|---|---|---|---|---|
| Q8K224 | Nat10 | 115.42 | 4.0123 | 3 | 4 | 486340 | 1037500 |
| Q6NS46 | Pdcd11 | 207.78 | 4.0064 | 5 | 8 | 220740 | 438800 |
| Q61033 | Tmpo | 75.167 | 4.0033 | 2 | 3 | 614660 | 968600 |
| Q8BFY9;Q99LG2 | Tnpo1;Tnpo2 | 102.36 | 3.9685 | 2 | 4 | 308290 | 660020 |
| Q3UGC7;Q66JS6 | Eif3j1;Eif3j2 | 29.343 | 3.9561 | 3 | 3 | 1591700 | 3939000 |
| P61358 | Rpl27 | 15.798 | 3.9445 | 6 | 6 | 23565000 | 34161000 |
| Q9JJ80 | Rpf2 | 35.363 | 3.9382 | 2 | 3 | 396260 | 3182900 |
| Q3TUH1 | Tamm41 | 37.827 | 3.9368 | 1 | 1 | 979160 | 1590700 |
| Q3THS6 | Mat2a | 43.688 | 3.929 | 5 | 6 | 4068500 | 11001000 |
| Q9D8U8 | Snx5 | 46.797 | 3.8942 | 0 | 1 | 0 | 0 |
| P17751 | Tpi1 | 26.712 | 3.8857 | 0 | 2 | 0 | 56927 |
| Q9Z1R2 | Bag6 | 121.04 | 3.8598 | 2 | 0 | 166420 | 0 |
| Q922Q4;Q922W5 | Pycr2 | 33.659 | 3.8451 | 1 | 3 | 782320 | 3720400 |
| P61982 | Ywhag | 28.302 | 3.8261 | 3 | 3 | 904030 | 1064100 |
| P24452 | Capg | 39.24 | 3.8157 | 2 | 2 | 1443200 | 2302100 |
| Q91VX2 | Ubap2 | 117.96 | 3.7456 | 2 | 0 | 691850 | 0 |
| Q3UX10 | Tubal3 | 49.988 | 3.7266 | 3 | 2 | 0 | 0 |
| P25799 | Nfkb1 | 105.61 | 3.6635 | 3 | 4 | 638900 | 815970 |
| Q05D44 | Eif5b | 137.61 | 3.6614 | 4 | 3 | 251130 | 413080 |
| Q9CQN1 | Trap1 | 80.208 | 3.656 | 5 | 6 | 1157400 | 1836400 |
| Q05144 | Rac2 | 21.441 | 3.6513 | 4 | 5 | 24296000 | 35028000 |

| | | | | | | | |
|---|---|---|---|---|---|---|---|
| Q61696;P17879 | Hspa1a;Hspa1b | 70.078 | 3.6395 | 5 | 6 | 0 | 346340 |
| Q922F4 | Tubb6 | 50.09 | 3.6275 | 10 | 11 | 454290 | 1070400 |
| P10107 | Anxa1 | 38.734 | 3.6199 | 4 | 3 | 338840 | 1046800 |
| P23198 | Cbx3 | 20.855 | 3.6001 | 1 | 1 | 2003300 | 3236400 |
| Q9WTK5 | Nfkb2 | 96.831 | 3.5953 | 1 | 2 | 297280 | 141380 |
| Q61703 | Itih2 | 105.93 | 3.5627 | 2 | 2 | 770200 | 798090 |
| Q08024 | Cbfb | 22.03 | 3.5406 | 1 | 1 | 599600 | 1271800 |
| P35279;P61294;Q8BHD0 | Rab6a;Rab6b | 23.59 | 3.5258 | 5 | 5 | 4066200 | 5408800 |
| Q99LC5 | Etfa | 35.009 | 3.5238 | 3 | 3 | 3095500 | 7439100 |
| Q99PT1 | Arhgdia | 23.407 | 3.5231 | 4 | 3 | 4889100 | 2997000 |
| O55143;Q8R429 | Atp2a2 | 114.86 | 3.5003 | 4 | 5 | 782130 | 1654000 |
| Q80UG5 | Septin9 | 65.574 | 3.4999 | 6 | 5 | 2729100 | 3909800 |
| Q8BJY1 | Psmd5 | 55.971 | 3.4962 | 1 | 3 | 174770 | 935360 |
| Q810V0 | Mphosph10 | 78.734 | 3.4924 | 1 | 1 | 521190 | 1204800 |
| Q8HWB2 | H2-Q4;H2-Q2; | 39.617 | 3.4622 | 3 | 4 | 0 | 908320 |
| Q9JJA4 | Wdr12 | 47.346 | 3.4616 | 1 | 1 | 847080 | 1219400 |
| Q9CU62 | Smc1a | 143.23 | 3.4526 | 4 | 4 | 299420 | 490230 |
| Q8VDF2 | Uhrf1 | 88.303 | 3.4461 | 5 | 6 | 458300 | 738220 |
| Q9D0L8 | Rnmt | 53.291 | 3.4378 | 3 | 2 | 1031100 | 896180 |
| P62141 | Ppp1cb | 37.186 | 3.4329 | 2 | 3 | 276070 | 672100 |
| Q8BH95 | Echs1 | 31.474 | 3.4319 | 0 | 2 | 0 | 428000 |
| Q9D883 | U2af1 | 27.815 | 3.4286 | 1 | 1 | 2660300 | 4373900 |

| | | | | | | | |
|---|---|---|---|---|---|---|---|
| P42228 | Stat4 | 85.94 | 3.4136 | 2 | 2 | 502360 | 842150 |
| Q6P5E4 | Uggt1 | 176.43 | 3.3838 | 1 | 2 | 150820 | 250570 |
| P62996 | Tra2b | 33.665 | 3.3812 | 3 | 3 | 5516000 | 6425100 |
| D3Z3N4 | Hnrnph3 | 36.869 | 3.3639 | 3 | 3 | 3266500 | 2568400 |
| O35326 | Srsf5 | 30.891 | 3.3611 | 1 | 3 | 816400 | 1696000 |
| Q8C0C7 | Farsa | 57.598 | 3.3493 | 4 | 6 | 1227200 | 3466000 |
| Q505F5 | Lrrc47 | 63.589 | 3.3472 | 3 | 4 | 388790 | 1515300 |
| Q99LE6 | Abcf2 | 71.781 | 3.3462 | 6 | 9 | 2582500 | 4466900 |
| Q9QY81 | Nup210 | 204.1 | 3.3446 | 2 | 2 | 92848 | 180350 |
| Q91VR2 | Atp5f1c | 32.886 | 3.3291 | 5 | 4 | 6355800 | 4970200 |
| Q9Z2I9 | Sucla2 | 50.113 | 3.3258 | 1 | 2 | 122190 | 229090 |
| F8VQC1 | Srp72 | 74.656 | 3.2999 | 1 | 3 | 132720 | 999460 |
| Q9DCW4 | Etfb | 27.623 | 3.2555 | 1 | 2 | 714330 | 1453200 |
| Q99JY9;Q641P0 | Actr3 | 47.357 | 3.2356 | 3 | 5 | 1418900 | 4261900 |
| Q3THK7 | Gmps | 76.723 | 3.2349 | 4 | 6 | 706240 | 1407000 |
| P27048;P63163 | Snrpb;Snrpn | 23.656 | 3.2286 | 2 | 2 | 1870300 | 1943900 |
| P62751 | Rpl23a | 17.695 | 3.2088 | 3 | 3 | 26346000 | 42917000 |
| Q8BG81 | Poldip3 | 46.132 | 3.2068 | 5 | 5 | 1523600 | 1729800 |
| O35465 | Fkbp8 | 43.528 | 3.1866 | 1 | 2 | 667350 | 0 |
| Q8BGS0 | Mak16 | 35.185 | 3.1619 | 1 | 1 | 1197800 | 2267800 |
| Q61187 | Tsg101 | 44.123 | 3.1509 | 0 | 1 | 0 | 0 |
| P19783 | Cox4i1 | 19.53 | 3.1484 | 2 | 3 | 1427300 | 2866800 |
| P16045 | Lgals1 | 14.866 | 3.1304 | 1 | 2 | 1248100 | 1020500 |
| P50247 | Ahcy | 47.688 | 3.129 | 4 | 3 | 1469700 | 2596800 |

| | | | | | | | |
|---|---|---|---|---|---|---|---|
| P14576 | Srp54;Srp54c | 55.72 | 3.1099 | 3 | 4 | 152800 | 410030 |
| Q99P88 | Nup155 | 155.12 | 3.1051 | 2 | 4 | 194770 | 598530 |
| Q60710 | Samhd1 | 75.892 | 3.0723 | 4 | 3 | 995240 | 1311900 |
| Q61699 | Hsph1 | 96.406 | 3.0595 | 2 | 4 | 309970 | 498980 |
| Q8BSL7 | Arf2;Arf1; | 20.746 | 3.0513 | 2 | 2 | 3396200 | 2706900 |
| P13864 | Dnmt1 | 183.19 | 3.0403 | 3 | 4 | 65452 | 279690 |
| P50171 | Hsd17b8 | 26.588 | 3.0339 | 1 | 1 | 0 | 776610 |
| Q8CCF0 | Prpf31 | 55.429 | 3.0273 | 1 | 3 | 367830 | 1267500 |
| Q3UTA4 | Eif1ad10;Eif1ad3; | 16.568 | 3.0173 | 2 | 2 | 9312800 | 6801900 |
| Q6PHQ9 | Pabpc4 | 72.242 | 3.0134 | 4 | 6 | 336320 | 665690 |
| O08749 | Dld | 54.272 | 3.0088 | 0 | 2 | 0 | 0 |
| Q9WUM5 | Suclg1 | 36.154 | 2.9936 | 4 | 4 | 2465100 | 5407100 |
| Q6P5B0 | Rrp12 | 143.13 | 2.9916 | 0 | 3 | 0 | 0 |
| Q60931 | Vdac3 | 30.752 | 2.9909 | 1 | 1 | 503780 | 1197000 |
| P62320 | Snrpd3 | 13.916 | 2.9899 | 2 | 2 | 8864000 | 13455000 |
| Q9DB96 | Ngdn | 35.658 | 2.956 | 2 | 2 | 862740 | 1673000 |
| O35841 | Api5 | 56.784 | 2.9521 | 2 | 2 | 930250 | 1327100 |
| Q61210 | Arhgef1 | 102.8 | 2.9307 | 3 | 4 | 770900 | 1257900 |
| Q8BTI8 | Srrm2 | 294.84 | 2.9153 | 1 | 3 | 29754 | 121680 |
| Q8VCG3 | Wdr74 | 42.636 | 2.8902 | 1 | 2 | 211440 | 1099000 |
| D3YXK2 | Safb | 105.1 | 2.8841 | 3 | 3 | 301990 | 1575400 |
| Q9CSH3 | Dis3 | 108.84 | 2.8724 | 3 | 4 | 318500 | 448900 |
| P56399 | Usp5 | 95.832 | 2.8688 | 2 | 2 | 718300 | 1168600 |
| Q9WUK4 | Rfc2 | 38.724 | 2.867 | 1 | 1 | 0 | 0 |

| | | | | | | | |
|---|---|---|---|---|---|---|---|
| P18155 | Mthfd2 | 37.863 | 2.8354 | 5 | 5 | 1630900 | 3096700 |
| O35344 | Kpna3 | 57.772 | 2.8353 | 3 | 2 | 1519200 | 2017300 |
| A0A2R8VHP3 | Gm5478 | 57.919 | 2.8345 | 7 | 8 | 7149800 | 99468000 |
| P43404 | Zap70 | 70.111 | 2.8003 | 4 | 2 | 363530 | 576430 |
| Q8C5R8 | Prps1l1 | 34.82 | 2.7955 | 2 | 3 | 1092100 | 2791300 |
| P97384 | Anxa11 | 54.079 | 2.7814 | 3 | 3 | 385150 | 1212000 |
| O35691 | Pnn | 82.435 | 2.7756 | 2 | 2 | 197490 | 645590 |
| Q80X50 | Ubap2l | 116.8 | 2.7712 | 2 | 1 | 817930 | 519950 |
| P62843 | Rps15 | 17.04 | 2.7633 | 2 | 2 | 13131000 | 17413000 |
| Q5SWD9 | Tsr1 | 92.104 | 2.7525 | 2 | 3 | 590840 | 1124800 |
| Q8BVK9 | Sp110 | 50.14 | 2.7515 | 2 | 2 | 831640 | 1544400 |
| Q3UM45 | Ppp1r7 | 41.291 | 2.7259 | 0 | 1 | 0 | 0 |
| Q60864 | Stip1 | 62.581 | 2.692 | 4 | 6 | 1652900 | 2862500 |
| Q62465 | Vat1 | 43.096 | 2.6899 | 2 | 2 | 1041800 | 1396800 |
| Q8CIE6 | Copa | 138.43 | 2.6805 | 2 | 4 | 216450 | 346050 |
| O35685 | Nudc | 38.358 | 2.6758 | 5 | 6 | 3384900 | 5823100 |
| Q9CYL5 | Glipr2 | 17.09 | 2.6633 | 2 | 2 | 3299500 | 5709300 |
| Q8K4L0 | Ddx54 | 97.747 | 2.6625 | 1 | 3 | 117770 | 311230 |
| Q9EST5 | Anp32b | 31.078 | 2.6619 | 1 | 1 | 1261900 | 1639500 |
| Q8CFQ3 | Aqr | 170.29 | 2.6398 | 2 | 3 | 52411 | 182210 |
| A0A1D5RLD8 | Gapdhrt2 | 35.812 | 2.6336 | 18 | 19 | 0 | 0 |
| Q9EPU4 | Cpsf1 | 160.82 | 2.6142 | 2 | 3 | 93745 | 135620 |
| P67984 | Rpl22 | 14.759 | 2.6082 | 3 | 3 | 23073000 | 31105000 |
| P22892 | Ap1g1 | 91.349 | 2.607 | 0 | 1 | 0 | 0 |
| Q8BGQ7 | Aars1 | 106.91 | 2.5948 | 1 | 4 | 0 | 674780 |

| | | | | | | | |
|---|---|---|---|---|---|---|---|
| O70310 | Nmt1 | 56.888 | 2.5889 | 0 | 1 | 0 | 0 |
| Q9QZ85 | Iigp1 | 47.571 | 2.5889 | 2 | 2 | 468550 | 847690 |
| B1AUN2 | Eif2b3 | 50.488 | 2.5762 | 2 | 2 | 649890 | 1117700 |
| P35700 | Prdx1 | 22.176 | 2.5653 | 2 | 3 | 3960300 | 10147000 |
| Q5XJY5 | Arcn1 | 57.229 | 2.5552 | 4 | 3 | 591340 | 539410 |
| Q9D1H7 | Get4 | 36.525 | 2.5538 | 0 | 1 | 0 | 0 |
| Q3UHX2 | Pdap1 | 20.605 | 2.5527 | 4 | 1 | 7711000 | 271230 |
| Q8K0C4 | Cyp51a1 | 56.775 | 2.5493 | 3 | 3 | 640030 | 1266600 |
| Q8C3J5 | Dock2 | 211.7 | 2.5418 | 1 | 2 | 84275 | 168540 |
| Q61166 | Mapre1 | 30.016 | 2.5281 | 3 | 4 | 1074000 | 1240100 |
| Q9DCC4 | Pycr3 | 28.721 | 2.5076 | 1 | 1 | 695860 | 1660200 |
| Q80VD1 | Fam98b | 45.349 | 2.4899 | 3 | 3 | 1823000 | 2045000 |
| Q9JI48 | Plac8 | 12.353 | 2.4875 | 1 | 1 | 19143000 | 13622000 |
| P63037 | Dnaja1 | 44.868 | 2.4863 | 2 | 2 | 2019600 | 4444800 |
| O54824 | Il16 | 141.43 | 2.4815 | 3 | 2 | 512920 | 204860 |
| P35821 | Ptpn1 | 49.593 | 2.4811 | 2 | 3 | 245720 | 686310 |
| P62889 | Rpl30 | 12.784 | 2.4697 | 2 | 2 | 7787100 | 10512000 |
| P97496 | Smarcc1 | 122.89 | 2.4694 | 1 | 3 | 236090 | 188390 |
| Q9Z127 | Slc7a5 | 55.872 | 2.4555 | 1 | 1 | 824250 | 1666900 |
| Q3TAQ9 | Wdr36 | 99.719 | 2.4489 | 0 | 3 | 0 | 675880 |
| Q9JLF6 | Tgm1 | 89.825 | 2.411 | 2 | 2 | 160870 | 472860 |
| Q91V12 | Acot7 | 42.536 | 2.3991 | 1 | 2 | 222940 | 1197700 |
| Q9DBJ1;O70250 | Pgam1 | 28.832 | 2.3898 | 2 | 3 | 969740 | 1455900 |
| Q9Z1Z2 | Strap | 38.442 | 2.3894 | 4 | 4 | 1207700 | 3001400 |
| Q6PGF5 | Bms1 | 145.41 | 2.3846 | 2 | 3 | 112480 | 356220 |

| | | | | | | | |
|---|---|---|---|---|---|---|---|
| Q60930 | Vdac2 | 31.732 | 2.3703 | 3 | 4 | 3505500 | 7095500 |
| Q05CL8;A2AMD0 | Larp7;Larp7-ps | 64.802 | 2.3683 | 1 | 2 | 0 | 1063600 |
| Q9DBY8 | Nvl | 94.475 | 2.3625 | 1 | 1 | 0 | 213000 |
| Q9Z2U1 | Psma5 | 26.411 | 2.3602 | 2 | 3 | 3022900 | 3496700 |
| P43277 | H1-3 | 22.099 | 2.3596 | 6 | 7 | 308030 | 14861000 |
| Q9Z1F9 | Uba2 | 70.568 | 2.3273 | 1 | 2 | 270160 | 866480 |
| Q62189 | Snrpa | 31.835 | 2.3235 | 1 | 2 | 2133100 | 4561300 |
| Q921I9 | Exosc4 | 26.249 | 2.3072 | 0 | 2 | 0 | 3067700 |
| O55135 | Eif6 | 26.511 | 2.3044 | 2 | 4 | 695450 | 1492800 |
| Q99P31 | Hspbp1 | 39.166 | 2.3008 | 0 | 2 | 0 | 533350 |
| P41241 | Csk | 50.716 | 2.299 | 2 | 4 | 525600 | 1613800 |
| P45952 | Acadm | 46.481 | 2.2759 | 3 | 3 | 1291500 | 2072800 |
| P08249 | Mdh2 | 35.611 | 2.2748 | 2 | 3 | 276150 | 651970 |
| Q3UJB0 | Sf3b2 | 98.197 | 2.2722 | 3 | 4 | 175380 | 606920 |
| Q9JLJ2 | Aldh9a1 | 53.514 | 2.2722 | 0 | 2 | 0 | 0 |
| Q9CW46 | Raver1 | 79.381 | 2.2652 | 1 | 1 | 393150 | 401000 |
| P59235 | Nup43 | 41.989 | 2.2507 | 1 | 1 | 1065600 | 1312500 |
| P70349 | Hint1 | 13.777 | 2.2451 | 2 | 3 | 2160700 | 3891900 |
| Q7TNP2 | Ppp2r1b | 65.934 | 2.2418 | 2 | 2 | 175960 | 326920 |
| Q9D0M3 | Cyc1 | 35.327 | 2.2373 | 3 | 1 | 1110800 | 772680 |
| Q9CR16 | Ppid | 40.742 | 2.2354 | 0 | 2 | 0 | 479560 |
| Q9D1R9 | Rpl34 | 13.293 | 2.2187 | 1 | 2 | 3647400 | 10867000 |
| P70698 | Ctps1 | 66.682 | 2.2162 | 4 | 4 | 987970 | 1362000 |
| Q9Z1N5 | Ddx39b | 49.035 | 2.1955 | 8 | 8 | 1218000 | 2443300 |
| O08915 | Aip | 37.605 | 2.1881 | 3 | 3 | 926590 | 1400800 |

| | | | | | | | |
|---|---|---|---|---|---|---|---|
| Q8K4Z5 | Sf3a1 | 88.544 | 2.1878 | 1 | 2 | 301980 | 665970 |
| O08528;Q91W97;P17710 | Hk2 | 102.53 | 2.183 | 2 | 4 | 514170 | 1202000 |
| Q99J77 | Nans | 40.024 | 2.1823 | 0 | 1 | 0 | 0 |
| Q9D8B3 | Chmp4b | 24.936 | 2.1811 | 1 | 0 | 0 | 0 |
| Q3UEB3 | Puf60 | 60.248 | 2.1619 | 2 | 2 | 558590 | 598980 |
| P62071 | Rras2 | 23.399 | 2.1451 | 1 | 1 | 347420 | 658420 |
| Q922H2 | Pdk3 | 47.922 | 2.1383 | 1 | 2 | 432870 | 826390 |
| P08003 | Pdia4 | 71.982 | 2.1197 | 1 | 1 | 294870 | 283450 |
| Q9WUA2 | Farsb | 65.696 | 2.1179 | 3 | 3 | 376740 | 847010 |
| P47753 | Capza1 | 32.939 | 2.1169 | 5 | 4 | 2787600 | 2388300 |
| P62305 | Snrpe | 10.803 | 2.1091 | 1 | 1 | 6877200 | 13564000 |
| O55201 | Supt5h | 120.66 | 2.1069 | 1 | 1 | 421670 | 443570 |
| Q99LB7 | Sardh | 101.68 | 2.1067 | 1 | 0 | 134650 | 0 |
| Q6Y7W8 | Gigyf2 | 149.19 | 2.1056 | 1 | 1 | 0 | 0 |
| P35486 | Pdha1 | 43.231 | 2.0986 | 1 | 1 | 0 | 0 |
| P68368 | Tuba4a | 49.924 | 2.0955 | 13 | 15 | 4529800 | 7948600 |
| Q8VDM6 | Hnrnpul1 | 96.001 | 2.0953 | 3 | 2 | 608170 | 353960 |
| Q91VK1 | Bzw2 | 48.063 | 2.0948 | 2 | 2 | 175380 | 1131000 |
| P49722 | Psma2 | 25.926 | 2.0806 | 1 | 1 | 540270 | 911890 |
| Q9QYA2 | Tomm40 | 37.895 | 2.0724 | 3 | 3 | 1468400 | 2019400 |
| P17897 | Lyz1 | 16.794 | 2.0655 | 1 | 1 | 14865000 | 24134000 |
| P63323 | Rps12 | 14.515 | 2.0582 | 3 | 3 | 19801000 | 6277700 |
| Q99P72 | Rtn4 | 126.61 | 2.0515 | 1 | 1 | 130700 | 145610 |
| Q6A026 | Pds5a | 150.33 | 2.0481 | 1 | 1 | 71547 | 111640 |
| E9PWZ3 | Rpl3l | 46.512 | 2.0477 | 1 | 1 | 103830 | 517710 |

| | | | | | | | |
|---|---|---|---|---|---|---|---|
| Q9D7X8 | Ggct | 21.166 | 2.0376 | 0 | 1 | 0 | 1284600 |
| Q6PHN9 | Rab35 | 23.025 | 2.0341 | 1 | 1 | 1059900 | 1317000 |
| Q8CGR7 | Upp2 | 35.755 | 2.0288 | 1 | 1 | 9458500 | 20370000 |
| Q8VE37 | Rcc1 | 44.93 | 2.0269 | 0 | 1 | 0 | 923520 |
| Q922V4 | Plrg1 | 56.937 | 2.0152 | 1 | 1 | 401870 | 385420 |
| Q8BH74 | Nup107 | 106.72 | 2.0146 | 0 | 1 | 0 | 128950 |
| Q9DBG7 | Srpra | 69.622 | 2.0097 | 1 | 0 | 0 | 0 |
| Q9D0R4 | Ddx56 | 61.211 | 1.9856 | 2 | 4 | 213110 | 1243500 |
| Q8CCS6 | Pabpn1 | 32.296 | 1.9846 | 1 | 1 | 0 | 687370 |
| Q8VCY6 | Utp6 | 70.429 | 1.9541 | 1 | 1 | 158270 | 133190 |
| O09106;P70288 | Hdac1;Hdac2 | 55.074 | 1.9409 | 3 | 2 | 1090500 | 2380100 |
| P26040;P26043 | Ezr;Rdx | 69.406 | 1.9232 | 6 | 7 | 8035200 | 10358000 |
| P05201 | Got1 | 46.247 | 1.915 | 0 | 1 | 0 | 0 |
| Q8K4L4 | Pof1b | 67.784 | 1.8949 | 0 | 3 | 0 | 1125500 |
| Q9D358 | Acp1 | 18.192 | 1.8885 | 1 | 1 | 720950 | 1101200 |
| P62874 | Gnb1 | 37.377 | 1.8875 | 1 | 4 | 0 | 958330 |
| Q9CZW5 | Tomm70 | 67.589 | 1.885 | 0 | 2 | 0 | 354390 |
| Q9R1P1 | Psmb3 | 22.965 | 1.8814 | 1 | 1 | 1123300 | 1559700 |
| Q9JLJ5 | Elovl1 | 32.677 | 1.8801 | 1 | 1 | 949010 | 2465100 |
| Q8CFI7 | Polr2b | 133.91 | 1.8601 | 0 | 2 | 0 | 181290 |
| Q9CQ65 | Mtap | 31.062 | 1.8559 | 1 | 2 | 785450 | 2284700 |
| Q9DAW6 | Prpf4 | 58.369 | 1.8549 | 1 | 2 | 0 | 770950 |
| P53986 | Slc16a1 | 53.267 | 1.8439 | 1 | 2 | 559070 | 2639200 |
| Q63850 | Nup62 | 53.254 | 1.8407 | 0 | 1 | 0 | 435870 |

| | | | | | | | |
|---|---|---|---|---|---|---|---|
| Q6PB66 | Lrpprc | 156.61 | 1.8291 | 4 | 4 | 50671 | 401150 |
| Q99JI6;P62835 | Rap1b;Rap1a | 20.825 | 1.8285 | 3 | 3 | 4724100 | 5633800 |
| Q9D7Z3 | Nol7 | 28.97 | 1.8241 | 1 | 1 | 1441900 | 3200500 |
| Q9DBG5 | Plin3 | 47.262 | 1.8208 | 1 | 1 | 279840 | 228290 |
| E9PVA8 | Gcn1 | 293.02 | 1.8191 | 0 | 2 | 0 | 33766 |
| Q99JR8;Q6P9Z1;Q61466 | Smarcd2; | 59.084 | 1.8172 | 2 | 2 | 259860 | 391920 |
| P57784 | Snrpa1 | 28.357 | 1.8108 | 2 | 2 | 1543300 | 3433400 |
| P63085 | Mapk1 | 41.275 | 1.8077 | 1 | 2 | 157360 | 512870 |
| Q9DAR7 | Dcps | 38.988 | 1.7994 | 1 | 2 | 340670 | 736870 |
| Q9CRB9 | Chchd3 | 26.334 | 1.7982 | 0 | 2 | 0 | 892690 |
| Q9CQT1 | Mri1 | 39.41 | 1.797 | 3 | 5 | 661970 | 2005400 |
| Q9CR67 | Tmem33 | 28.031 | 1.7898 | 1 | 1 | 0 | 1121400 |
| O54734 | Ddost | 49.027 | 1.7842 | 3 | 3 | 1576000 | 1994600 |
| Q3UJB9 | Edc4 | 152.48 | 1.7836 | 0 | 1 | 0 | 131480 |
| Q08943 | Ssrp1 | 80.859 | 1.7783 | 1 | 2 | 190440 | 458150 |
| G3X987 | Gimap9 | 33.298 | 1.7782 | 1 | 1 | 265990 | 493820 |
| Q9JIF0 | Prmt1 | 42.435 | 1.775 | 3 | 2 | 499050 | 1521300 |
| P13379 | Cd5 | 53.849 | 1.7715 | 2 | 3 | 906010 | 1134400 |
| P39054;Q8BZ98 | Dnm2 | 98.144 | 1.7657 | 3 | 5 | 263410 | 527730 |
| Q9DC51 | Gnai3 | 40.538 | 1.7647 | 2 | 3 | 1134800 | 1552300 |
| Q8BHX3 | Cdca8 | 32.214 | 1.7527 | 1 | 1 | 263430 | 619380 |
| Q8BJ71 | Nup93 | 93.28 | 1.7509 | 2 | 3 | 239710 | 645260 |
| Q9JJT0 | Rcl1 | 40.84 | 1.7496 | 1 | 1 | 181230 | 380890 |

| | | | | | | | |
|---|---|---|---|---|---|---|---|
| Q7TQI3 | Otub1 | 31.27 | 1.7457 | 1 | 1 | 558500 | 1124700 |
| Q8BH59 | Slc25a12 | 74.569 | 1.7361 | 0 | 1 | 0 | 549640 |
| P68134 | Acta1;Actc1; | 42.051 | 1.7329 | 13 | 18 | 2295100 | 4019000 |
| P10852 | Slc3a2 | 58.336 | 1.7279 | 1 | 2 | 764570 | 2230000 |
| Q91YH5 | Atl3 | 60.574 | 1.7225 | 1 | 1 | 477680 | 621910 |
| Q9D6T0 | Nosip | 33.209 | 1.7186 | 1 | 1 | 515930 | 881590 |
| Q6A0A9 | FAM120A | 121.64 | 1.7175 | 2 | 2 | 223940 | 276680 |
| Q9CX34 | Sugt1 | 38.158 | 1.7154 | 4 | 4 | 3085600 | 4180600 |
| P32233 | Drg1 | 40.512 | 1.7142 | 2 | 3 | 0 | 0 |
| A2AJ72 | Fubp3 | 61.447 | 1.7109 | 1 | 0 | 0 | 0 |
| Q6PFR5 | Tra2a | 32.316 | 1.7025 | 1 | 1 | 1129200 | 1781700 |
| Q9CQA3 | Sdhb | 31.814 | 1.6966 | 0 | 1 | 0 | 1019200 |
| Q9JM76 | Arpc3 | 20.524 | 1.6963 | 2 | 2 | 1439700 | 2555900 |
| Q9R190 | Mta2;Mta3; | 75.029 | 1.6842 | 1 | 1 | 366080 | 0 |
| Q9WUP7 | Uchl5 | 37.616 | 1.6623 | 1 | 2 | 0 | 0 |
| Q91V64 | Isoc1 | 32.032 | 1.6588 | 0 | 1 | 0 | 708660 |
| Q6PDG5 | Smarcc2 | 132.6 | 1.6536 | 0 | 2 | 0 | 199050 |
| Q9QUM9 | Psma6 | 27.372 | 1.6425 | 2 | 2 | 758120 | 1813100 |
| Q62418 | Dbnl | 48.699 | 1.6401 | 2 | 3 | 610620 | 1016200 |
| Q99KQ4 | Nampt | 55.446 | 1.6338 | 3 | 2 | 676990 | 832670 |
| Q9CQC6 | Bzw1 | 48.043 | 1.6324 | 4 | 4 | 1987800 | 3288600 |
| P35293 | Rab18 | 23.035 | 1.6289 | 1 | 1 | 621940 | 854500 |
| P62858 | Rps28 | 7.8409 | 1.6264 | 2 | 1 | 23226000 | 12468000 |
| Q62186 | Ssr4 | 18.936 | 1.6244 | 2 | 1 | 994020 | 1803200 |
| Q9CPQ1 | Cox6c | 8.4689 | 1.6202 | 1 | 1 | 1062900 | 1706800 |
| O35490 | Bhmt | 45.02 | 1.6163 | 1 | 1 | 1768500 | 1114400 |

| | | | | | | | |
|---|---|---|---|---|---|---|---|
| Q9Z0X1 | Aifm1 | 66.765 | 1.5782 | 2 | 4 | 196180 | 806130 |
| Q9CPT5 | Nop16 | 21.139 | 1.5716 | 1 | 2 | 887880 | 2737800 |
| Q8C4J7 | Tbl3 | 88.265 | 1.565 | 1 | 1 | 165370 | 333250 |
| P30681 | Hmgb2 | 24.162 | 1.5607 | 1 | 1 | 616400 | 836850 |
| Q60848 | Hells | 95.125 | 1.5526 | 1 | 2 | 221400 | 366230 |
| Q64511 | Top2b | 181.91 | 1.5477 | 3 | 5 | 102970 | 197800 |
| Q6P1Y8 | Inpp4b | 104.53 | 1.5452 | 1 | 1 | 44366 | 73847 |
| Q9Z2U0 | Psma7 | 27.855 | 1.5443 | 4 | 3 | 2894800 | 2272500 |
| Q08189 | Tgm3 | 77.308 | 1.5381 | 1 | 1 | 508470 | 1018500 |
| Q8BGA5 | Krr1 | 43.537 | 1.5378 | 0 | 1 | 0 | 518340 |
| F7ABZ6 | Dnah10 | 522.69 | 1.5367 | 0 | 1 | 0 | 0 |
| Q9WV32 | Arpc1b | 41.063 | 1.535 | 1 | 1 | 1215200 | 1835400 |
| Q9CZ44 | Nsfl1c | 40.709 | 1.5342 | 1 | 0 | 248180 | 0 |
| Q9QYJ3 | Dnajb1 | 38.167 | 1.5072 | 2 | 1 | 696160 | 537150 |
| Q8QZY9 | Sf3b4 | 44.355 | 1.4972 | 1 | 1 | 633560 | 988280 |
| E9Q5C9 | Nolc1 | 73.697 | 1.4947 | 1 | 2 | 1122800 | 2209100 |
| Q60766 | Irgm1 | 46.551 | 1.4867 | 1 | 1 | 464410 | 585870 |
| Q9Z2X8 | Keap1 | 69.552 | 1.4831 | 0 | 1 | 0 | 0 |
| Q69ZQ2 | Isy1 | 32.989 | 1.4831 | 0 | 1 | 0 | 0 |
| E9Q6L7 | Gsta5;Gm37 76; | 25.506 | 1.4831 | 1 | 1 | 505970 | 156810 |
| Q9JJ28 | Flii | 144.8 | 1.4797 | 1 | 1 | 112810 | 0 |
| G5E870 | Trip12 | 224.13 | 1.4787 | 0 | 1 | 0 | 54143 |
| Q9EQP2 | Ehd4 | 61.48 | 1.4766 | 2 | 3 | 582400 | 563390 |
| P61202 | Cops2 | 51.596 | 1.4661 | 3 | 2 | 403500 | 528900 |
| Q3U821 | Wdr75 | 94.036 | 1.4512 | 2 | 2 | 221010 | 563620 |

| | | | | | | | |
|---|---|---|---|---|---|---|---|
| Q60611 | Satb1 | 85.879 | 1.4459 | 2 | 1 | 603180 | 251900 |
| P55194 | Sh3bp1 | 74.172 | 1.4392 | 1 | 0 | 0 | 0 |
| Q3TTA7 | Cblb | 109.09 | 1.4383 | 1 | 1 | 0 | 49907 |
| Q61176 | Arg1 | 34.807 | 1.4322 | 1 | 1 | 1144000 | 3218400 |
| O08784 | Tcof1 | 135 | 1.4288 | 2 | 2 | 352170 | 592300 |
| Q99PU8 | Dhx30 | 136.67 | 1.4227 | 1 | 2 | 0 | 64310 |
| P34884 | Mif | 12.504 | 1.4197 | 1 | 1 | 1926200 | 4007400 |
| P63158 | Hmgb1 | 24.893 | 1.4179 | 0 | 1 | 0 | 0 |
| Q62193 | Rpa2 | 29.44 | 1.4089 | 1 | 1 | 610780 | 869810 |
| Q61550 | Rad21 | 72.082 | 1.4076 | 1 | 0 | 0 | 0 |
| O35226 | Psmd4 | 40.703 | 1.4001 | 1 | 1 | 1276900 | 1488800 |
| Q8R2Y8 | Ptrh2 | 19.526 | 1.3986 | 2 | 2 | 454580 | 644930 |
